# Archaic ancestry inference in imputed ancient human genomes

**DOI:** 10.1101/2025.07.23.664192

**Authors:** M.R. Capodiferro, L. Planche, E.M. Breslin, L. Ongaro, M.C. Ávila-Arcos, F. Jay, L.M. Cassidy, E. Huerta-Sanchez

## Abstract

When modern humans expanded from Africa into Eurasia, they interbred with archaic hominins such as Neanderthals and Denisovans. This groundbreaking discovery, made in part possible through the genomic analysis of archaic remains, reshaped our understanding of human origins, and opened new research avenues to study the effects and consequences of archaic introgression. While significant progress has been made in identifying and quantifying archaic introgression, most studies have focused on contemporary individual genomic data. As a result, relatively little is known about the evolution of archaic variants within modern human populations shortly after interbreeding occurred. While analysis of ancient DNA from modern humans offers the potential to address this scientific gap, its poor quality has hindered its exploration. However, recent studies have shown that imputation using contemporary reference panels can accurately infer missing genotypes in ancient genomes. Here, we investigate the feasibility of using imputation to improve both global and local archaic ancestry inference in ancient genomes, by downsampling to different low-coverage values and imputing 20 high-coverage (>10X) ancient genomes, representing individuals from diverse temporal and geographical contexts.

We tested the reliability of detecting and quantifying archaic introgression using *D-statistics* and the *f4-ratio*. We identified consistent results from the imputed and the original ancient genomes, suggesting that genomic estimates to detect and quantify introgression work well in imputed genomes. Regarding Local Ancestry Inference we find that we can identify more introgressed segments in imputed genomes than in non-imputed genomes. Surprisingly, we find that imputation accuracy is even higher in regions of archaic ancestry compared to other regions of the genome, facilitating the detection of introgressed segments in imputed genomes. Imputation also allows detection of Denisovan segments in Siberian and Alaskan individuals. We show that segments identified even at 0.0625X coverage can be used to reconstruct the history of introgressed haplotypes. For example, comparisons with archaic segments in contemporary humans reveal that the oldest individual analyzed, Ust’Ishim, carried segments that are now found exclusively in either European or East Eurasian populations, indicating that these segments co-occurred in a shared ancestral population. By comparing only ancient individuals, we demonstrate that clustering based solely on the identified archaic segments effectively groups individuals into genetic clusters that correspond to populations defined by geography and time. This analysis shows a clear distinction between individuals with Denisovan introgression and those without, as well as a more detailed separation between Mesolithic and Neolithic Europeans, with the latter closely resembling Central Eurasian individuals. Furthermore, despite the small sample size in this study, we are able to reconstruct the origins of genes identified as candidates for adaptive introgression in contemporary populations, such as the *BCN2* gene, which is an adaptively introgressed gene identified in contemporary West Eurasians. Additionally, we identify new candidates for adaptive introgression including *LEMD2* and *MLN* in Europe. In this gene-region, imputation helps resolve the introgressed haplotype, which is closest to the Vindija Neanderthal haplotype.

## Introduction

Despite recent advances in identifying and quantifying archaic introgression in contemporary humans^1^, its inference in ancient genomes has been relatively neglected due to the poor quality of ancient DNA (aDNA). Analysis of ancient genomes could provide valuable insights into the number, magnitude and timing of introgression events and the post-introgression (~40,000 years ago) evolution of archaic variants within modern human genomes, including the identification of adaptive introgression signals.

This deficiency is mainly attributed to low coverage and sequencing errors^2^ leading to large gaps in genomic coverage. Consequently, many studies on ancient genomes resort to pseudohaploidization (i.e. selecting a single random allele or read at each site) rather than using genotypes. These limitations often render approaches developed for analyzing contemporary genomes unsuitable for aDNA. While archaic Global Ancestry Inference (GAI) may still be suitable for ancient genomes with sufficient data, archaic Local Ancestry Inference (LAI) necessitates high-quality genotype data and is highly susceptible to missing data. Indeed, the only software specifically developed for this task, Admixfrog^3^, provides a useful option for working with aDNA. However, depending on the coverage of the ancient genomes, the amount of archaic segments may be underestimated.

A promising solution is genotype imputation, which allows for the inference of missing or incorrect genotypes—caused by sequencing at low depth or damage in aDNA— by using a reference panel of present-day haplotypes, facilitating haplotype-based inference and increasing the depth of analysis. The quality of the inferred genotypes is influenced not only by the coverage of ancient genomes, but also by the frequency of alleles in the reference dataset^4^. Given the small amounts of introgressed archaic segments in modern humans (0-6%), it is unclear whether imputation is particularly challenging in archaic segments, and whether introgressed regions can be reliably inferred in imputed ancient genomes.

Here, we investigate the feasibility of inferring archaic introgression in imputed ancient genomes and measure our ability to reconstruct introgressed fragments within ancient individuals using a present-day reference panel. Our strategy is based on a comparative approach (Figure 1A), in which we evaluate the results of specific analyses across six different versions of previously published high-coverage shotgun-sequenced genomes. These include the original high-coverage data (original), the imputed version of the same Original high-Coverage genome (imputed OC), and imputed versions obtained after downsampling to various low-coverage (2×, 1×, 0.5×) and a very low-coverage level (0.0625×). We selected 20 high-coverage (>10x) genomes representing ancient individuals from Eurasia, the Americas, and Africa, spanning a broad temporal range from approximately 44,000 to 5,000 years Before Common Era (BCE) (Table S1, Figure 1B)^5–18^. Imputation was performed using GLIMPSE^19^ with the 1000 Genomes Project dataset as the modern reference panel.

**Figure 1.**
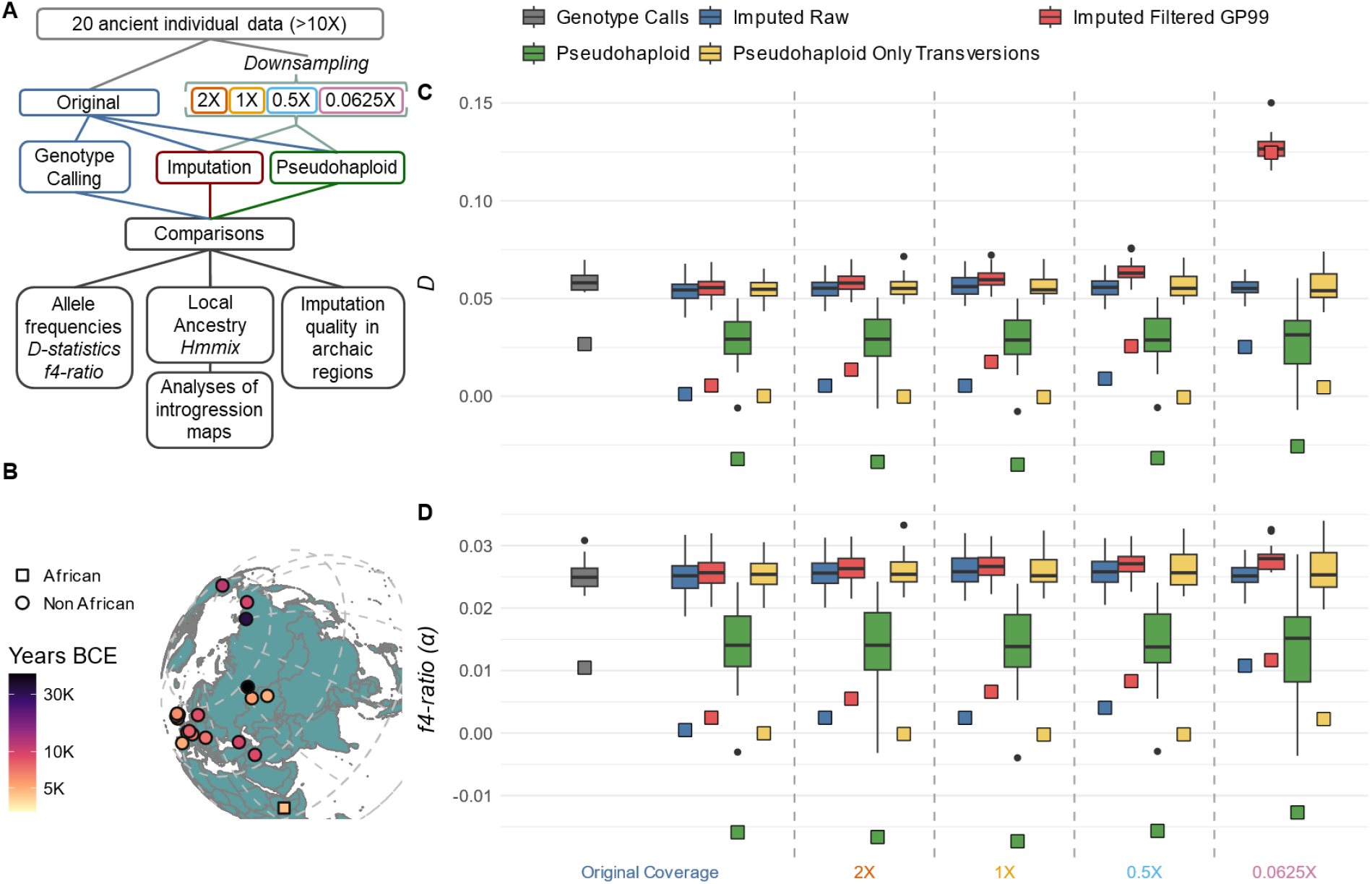
**A)** Workflow illustrating the methods used to test the inference of archaic introgression in imputed ancient DNA. **B)** Map showing the geographical locations of the samples included in this study. The colors represent the estimated date of each individual in thousands of years before present (BCE). **C)** Boxplot of D-statistics calculated using AdmixTools in the format D(Africa, Ancient Sample; Altai Neanderthal, Chimp), plotted against the sequencing depth before imputation (x-axis). **D)** Boxplot of α values obtained from the f4-ratio in the format f4(Altai, Chimp; Ancient Sample, Africa) /f4(Altai, Chimp; Vindija, Africa), also plotted against sequencing depth before imputation. For panels (C) and (D), each coverage level includes four boxplots, which depict the distribution of the respective statistics measured at the individual level, representing different data types. Two are based on imputed data: Blue, without quality filtering; Red, filtered for genotype probability (GP ≥ 99%). The other two reflect results from pseudohaploid data, where one random allele is selected per position: Green, without filtering; Yellow, retaining only transversions, to reduce the impact of post-mortem damage. The highlighted squares indicate the African individual Mota, used as a negative control.

To evaluate imputation, we examined three key aspects: the reliability of identifying archaic introgression, the ability to quantify the amount of introgression, and the precision in pinpointing the genomic locations of introgressed haplotypes in imputed ancient genomes. We validate the viability of utilizing imputed genomes for inferring archaic introgression and find that imputation accuracy is higher in archaic introgressed regions compared to the average in non-introgressed regions. Even in scenarios where high-coverage genomes are available, imputation offers a valuable avenue for refining and expanding our understanding of archaic introgression in ancient genomes.

## Results and discussion

### Using allele sharing to identify and quantify archaic introgression

#### D-statistics to identify archaic introgression

To test the ability to detect allele sharing between archaic and ancient (imputed and original) genomes, we employed *D-statistics* in the form *D(Africa, Test Individual; Archaic, Outgroup)* using the *qpDstat* software from the AdmixTools package^20^. Our results show comparable outcomes (*D*=0.05-0.06) among the imputed genomes and the original ancient genome (i.e. “the truth”, Figure 1). The significance of the signals, as indicated by the *Z-score* in Admixtools, further supports the results (Figure S1). The pseudohaploid data show an underestimation of the *D* value when all positions are considered, however, when only transversions are used, it is very consistent with the imputed and original data (Figure 1).

As in previous studies^4^, we applied filters for genotype probability (GP ≥ 99%) and minor allele frequencies in the reference panel (MAF ≥ 5%), which are expected to yield the most reliable calls. While overall consistency is maintained down to 0.5× coverage, applying the GP filter at 0.0625× coverage causes the *D-statistic* to become unreliable, leading to an increase in the *D* value (Figure 1C). In contrast, applying only the MAF filter does not produce this bias (Figure S1). One way to correct this is to weight the frequency of each allele by its GP values rather than being represented as a discrete frequency of 0, 0.5, or 1 (Supplementary Material). Figure S2 shows that this approach brings complete consistency across coverages to the values of the D-statistic in imputed genomes, even as low as 0.0625×, and better matches the results obtained from original genomes.

In the overall analysis, the African individual Mota—used as a negative control—shows some unexpected signals, when only a reduced amount of SNPs can be explored, such as positive *Ds* and significant *Z-scores* (see Supplementary Materials). This could reflect either traces of archaic ancestry from back migrations or the well-known limitations in imputation accuracy for African genomes^4^.

#### Global Ancestry Inference (GAI) using the f4-ratio

To assess imputation’s ability to estimate the overall proportion of archaic introgression in ancient genomes, we applied the *f4-ratio* approach^21^ in the form *f4(Archaic1, Chimp; Test, Africa) /f4(Archaic2, Chimp; Test, Africa)*, using the *qpF4Ratio* software from the Admixtools package^20^.

We observed consistent results across different coverages, with values ranging between 2% and 3%, comparable to those obtained from original genomes (Figure 1D). Applying a GP ≥ 99% filter revealed a slight increase in archaic ancestry, which becomes more pronounced as sequencing depth decreases. However, the bias is small and we observe similar proportion results across individuals. The only exceptions to the overall consistency of the *f4-ratio* results arise with Mota and pseudohaploid data when using all sites. Mota shows a slight overestimation of archaic ancestry in both original and very low-coverage imputed genomes (0.0625×), while pseudohaploid data can even produce negative *α* values and more consistent estimates are obtained when restricting the analysis to transversions.

To test whether GAI is influenced by sample age or original coverage, we assessed their correlation with *α* in non-African individuals (excluding Mota) (Figure S3). As expected, *α* correlates with age but not with original coverage in any downsampled dataset.

### Local Ancestry Inference

#### Identification of archaic segments in imputed vs non imputed ancient genomes

We evaluated the ability of imputation to improve the identification of archaic segments in ancient genomes. We use *hmmix*^*22*^ to detect introgressed segments because i) it was developed for high-quality data and we want to show that such methods can be applied to imputed ancient genomes and ii) it is reference-free making it more suitable for detecting Denisovan ancestry^23^. We applied *hmmix*^*22*^ to identify archaic segments in all original and imputed (filtered for GP ≥ 99%) individual genomes. We observed that imputation significantly enhances the identification of introgressed segments in imputed aDNA. Both the number of the introgressed region and the percentage of the genome identified as introgressed are higher in imputed genomes than in the original data, except in cases of very low-coverage (Figure 2A, S4).

**Figure 2.**
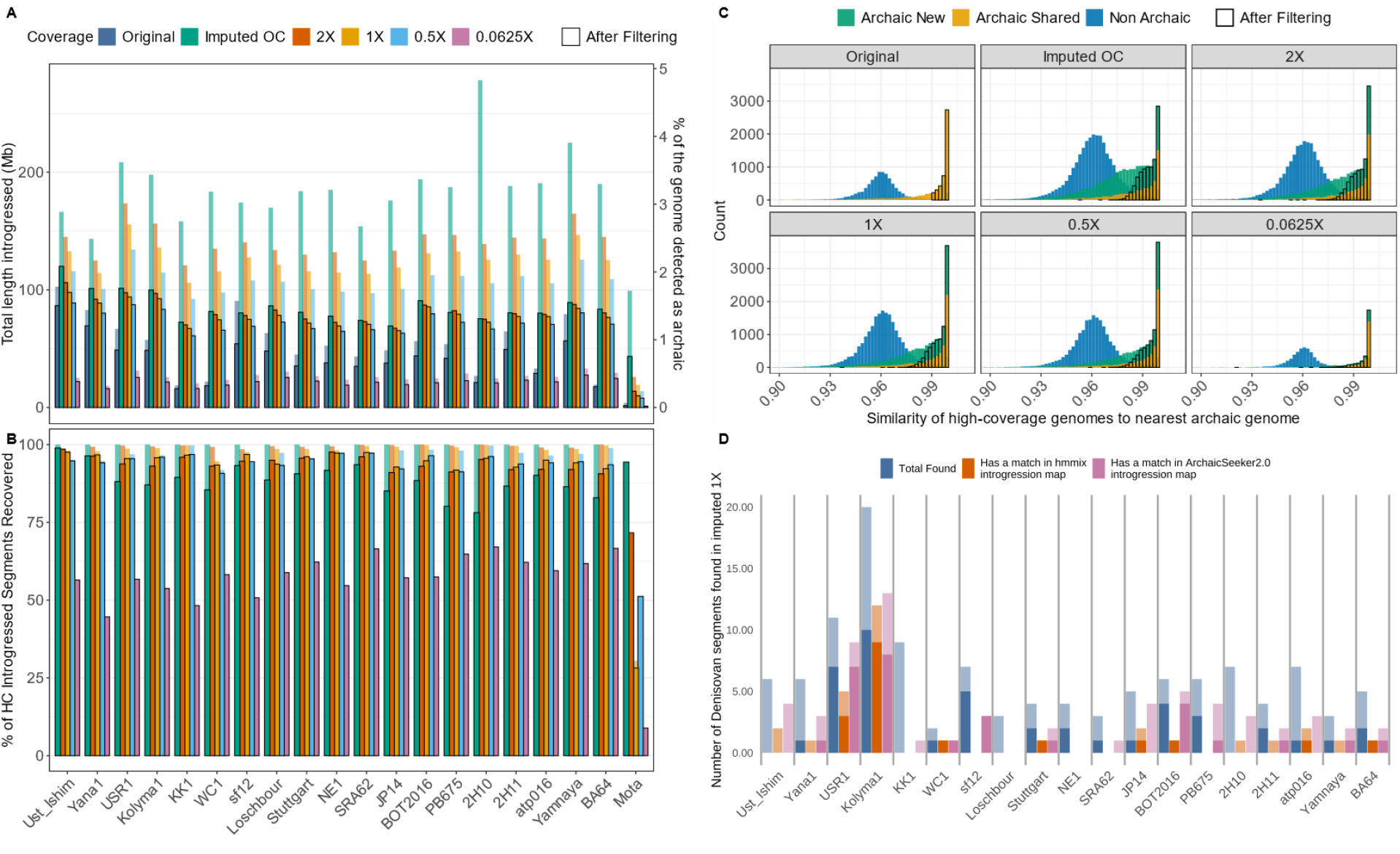
**A)** Sum of the length of the segments identified in each individual for each of the dataset expressed in megabases (Mb) and also expressed on the right axis as the percentage of the genome detected as archaic. The bordered bars for A and B represent the respective measures after filtering for quality, considering a minimum length of 40Kb and a similarity measured with the imputed data above 0.99 **B)** Proportions of the archaic segments detected using the original genome, that are also detected using imputed dataset. Over all the samples, respectively 100%, 99%, 99%, 97% and 50% of the segments detected using the original genome are also detected using respectively the Imputed OC, 2×, 1×, 0.5× and 0.0625× genomes. The bordered bars for A and B represent the respective measures after filtering for quality, considering a minimum length of 40Kb and a similarity measured with the imputed data 0.99 **C)** Similarity to the nearest archaic genome (between Vindija and Denisovan) of the segments detected in Imputed OC, 2×, 1×, 0.5× and 0.0625×. The similarity is computed by looking at the corresponding individual genotype in the original genome; this step is to avoid taking into account archaic variants which would have been wrongly introduced with imputation. In green the segments that are only detected in the imputed genome (Archaic New), in orange the segments that are detected both in the imputed and original genome (Archaic Shared). In these plots the box represents the proportion of each value after the quality filtering **D)** Denisovan segments identified in the 1X dataset. In blue the total numbers of inferred Denisovan segments, in orange the number of such segments which matched a Denisovan segment found in hmmix^22^ introgression map and in pink the number of such segments which matched a Denisovan segment found in ArchaicSeeket2.0^24^ introgression map. The dark shade corresponds to the segments also found in the original genome, the light shade represents segments only found after imputation.

The imputed genomes exhibit less variation between samples (except for Mota) in the detected amount of introgression compared to original genomes where it is proportional to the sequencing coverage of the genome (Figure S4). Remarkably, more than 97% of the archaic segments detected in original genomes are still identified when the data are downsampled to 0.5× and then imputed, and even at 0.0625× coverage, half of the archaic segments remain detectable (Figure 2B).

#### Quality of the inferred archaic introgressed segments

To confirm the introgressed segments identified are of archaic origin, we measured the genetic similarity of the original genome (not imputed) to the nearest archaic genome (Vindija Neanderthal or Denisova3) in genomic regions classified as introgressed in the imputed genomes (Figure 2C). We categorized these archaic regions into two groups: Archaic Shared, which includes regions detected in both imputed and original genomes, and Archaic New, consisting of regions identified only in the imputed ones.

While the Archaic Shared segments exhibit the highest mean similarity to the archaic reference genome, the Archaic New segments also show significantly higher similarity compared to regions classified as non-archaic (Figure 2C). This suggests that the original genomes do carry those introgressed segments, but they failed to be detected. In this case, it is important to consider that despite the confidence in the genotype calls from original genomes, sequencing errors, missing data and regions of low-coverage can still result in incorrect genotypes, which may impact the detection of introgressed segments. We further expanded this concept by introducing a new distance measure, providing a more principled framework for classification, normalizing the similarity by dividing it by the average similarity of modern African individuals (Biaka from HGDP) to the archaic genomes in each segment. When evaluating segments using both similarity and distance, we observe a clear separation between the distributions of archaic and non-archaic segments (Figure S5).

Building on this, we applied additional stringent quality filters to exclude segments (potential false positives) that are too short (<40 kb) and lack strong genetic affinity to archaic genomes (similarity <0.99). With these filters we observed that the proportion of introgressed segments identified in the imputed genomes, compared to the original, more closely aligns with the *f4-ratio* estimates obtained using original data (Figure S6). Moreover, considering only the filtered results (the bar with the black border in Figure 2A), the older individuals (shown on the left) have a higher proportion of their genome inferred to be of archaic origin across all coverage levels, consistent with expectations.^23,25^.

#### Identification of Denisovan origin introgressed segments

We assessed the potential of imputation to improve the detection of Denisovan segments in ancient genomes, a task complicated by the reliance on a single high-coverage Denisovan reference and the complex history of Denisovan introgression^23,25^. To evaluate imputation performance, we applied conservative filters, classifying a segment as Denisovan if (i) it was detected as archaic, (ii) had a distance <0.2 to the Denisovan genome, and (iii) was at least 0.3 closer to Denisovan than Neanderthal.

We observed that Kolyma1 (Siberia) and USR1 (Alaska) harbored the most Denisovan segments in both original and imputed data, reflecting their Ancient North Siberian (ANS) and East Asian ancestry^18^, while Yana1 (ANS) showed fewer segments (Figure 2D, S7). Comparison with introgression maps from contemporary individuals (*hmmix*,^22^; *ArchaicSeeker*^*24*^) showed that Kolyma1’s Denisovan segments aligned with previous studies. Therefore using imputed genotypes, we recovered most original segments (down to 0.5× coverage) and identified additional Denisovan segments (Figure 2D, S7).

#### Impact of imputation on the reconstruction of the archaic introgressed segments’ full length

Introgressed segment length in original data correlates with sample age, as expected^25,26^, with Ust’Ishim and Yana1 encompassing the longer segments, but imputation reduces this signal, making ancient genomes distribution of segment length more similar to that of modern ones (see Supplementary Materials, Figure S8). To further explore this limitation, we analyzed segment length based on (i) similarity to the archaic genome and (ii) whether segments were Archaic Shared or Archaic New. Shared segments display length distributions more similar to original data (Figure S9A) and longer than Archaic New. Nevertheless, this only marginally affects similarity: length and similarity distributions remain comparable between shared and new segments (Figure S9B).

### Quality of Imputation in archaic vs not archaic genomic regions

#### Imputation accuracy in introgressed regions

In the previous sections we have shown that we can detect and identify introgressed regions in low coverage imputed genomes and estimate the proportion of introgression using *f-statistics*. Encouraged by this, we wanted to check whether introgressed regions and variants are easier or harder to impute and understand why we do not detect some introgressed segments in the original genomes.

We first assessed how well imputation recovers archaic variants. We defined a variant as archaic if it is absent in African populations (Yoruba, Mende and Esan from the 1000 Genome Project) and homozygous in Vindija Neanderthal, and archaic SNPs are positions carrying these variants.

For each sample, we compared archaic SNPs between original and imputed genomes (Table 1). We observe that imputation infers archaic variants in introgressed regions as expected—both when the original genome has the archaic variant and when the position is missing, and assigns non-archaic variants in non-introgressed regions. For example, in both original and imputed OC genomes, in non-archaic regions, archaic variants appear at a rate of 0.3%. For Archaic Shared regions, the rates are 88% and 89% for original and imputed OC respectively, while for Archaic New regions, these values are 75% and 79%. Notably, the Archaic New segments contain more missing positions in the original data. This pattern confirms that the missingness of archaic variants in the original data could prevent *hmmix* from classifying these regions as introgressed. Finally, we measured the error rate of imputation in archaic SNPs, which is remarkably low at approximately 0.2%, even at very low-coverage such as 0.0625X.

**Table 1.** Rate of Archaic Variant Imputation in Imputed Ancient Individuals. A variant is defined as archaic if it is absent in African populations (Yoruba, Mende and Esan from the 1000 Genomes Project), present in non-Africans and Vindija Neanderthal is homozygous for that variant. We look at the variants inferred by imputation in archaic SNPs positions, separately for regions detected as Non Archaic, Archaic Shared and Archaic new. The first column, in the form “x->y” gives the variant in the original genome (x) and after imputation (y). The variant can take the value “0” (non archaic), “1” (archaic) and “.” (missing). We then count for each coverage the number of sites corresponding to each possible value and the numbers in the parenthesis indicate the absolute values for each category. The percentages in each column add up to 100% within each Segment Classification (i.e. Non Archaic, Shared Archaic and New Archaic) written in the last column. To make the numbers more comparable, the counts for “Non-Archaic” are computed from a set of random segments chosen to have the same length distribution as the Archaic segments. Results are shown for all imputed datasets.

| Segment Classification | Original -> Imputed | Imputed OC | 2X | 1X | 0.5X | 0.0625X |
| --- | --- | --- | --- | --- | --- | --- |
| Non Archaic | 0 -> 0 | 44%<br>(143262) | 44%<br>(124982) | 45%<br>(111579) | 45%<br>(96132) | 42%<br>(20371) |
| Non Archaic | . -> 0 | 55%<br>(179722) | 55%<br>(155854) | 55%<br>(137760) | 55%<br>(118485) | 57%<br>(27554) |
| Non Archaic | 0 -> 1 | 0%<br>(0) | 0.001%<br>(3) | 0.002%<br>(5) | 0.001%<br>(2) | 0%<br>(0) |
| Non Archaic | 1 -> 1 | 0.1%<br>(432) | 0.2%<br>(431) | 0.1%<br>(318) | 0.2%<br>(325) | 0.07%<br>(38) |
| Non Archaic | . -> 1 | 0.2%<br>(577) | 0.2%<br>(554) | 0.2%<br>(451) | 0.2%<br>(424) | 0.1%<br>(61) |
| Non Archaic | 1 -> 0 | 0.001%<br>(3) | 0.003%<br>(10) | 0.003%<br>(7) | 0.007%<br>(14) | 0.04%<br>(20) |
| Archaic Shared | 0 -> 0 | 6%<br>(8415) | 5%<br>(7517) | 4%<br>(6687) | 4%<br>(5564) | 1%<br>(664) |
| Archaic Shared | . -> 0 | 6%<br>(8346) | 5%<br>(8208) | 5%<br>(7394) | 4%<br>(6223) | 2%<br>(968) |
| Archaic Shared | 0 -> 1 | 0.1%<br>(166) | 0.2%<br>(364) | 0.2%<br>(372) | 0.2%<br>(352) | 0.3%<br>(140) |
| Archaic Shared | 1 -> 1 | 43%<br>(63368) | 42%<br>(64940) | 43%<br>(64647) | 44%<br>(62349) | 46%<br>(23863) |
| Archaic Shared | . -> 1 | 45%<br>(65871) | 46%<br>(70073) | 47%<br>(70219) | 48%<br>(68013) | 50%<br>(26483) |
| Archaic Shared | 1 -> 0 | 0.01%<br>(12) | 0.03%<br>(49) | 0.04%<br>(65) | 0.05%<br>(74) | 0.01%<br>(6) |
| Archaic New | 0 -> 0 | 9%<br>(4584) | 8%<br>(4625) | 7%<br>(4142) | 7%<br>(3538) | 4%<br>(437) |
| Archaic New | . -> 0 | 12%<br>(5929) | 11%<br>(6668) | 11%<br>(6102) | 10%<br>(5385) | 7%<br>(820) |
| Archaic New | 0 -> 1 | 0.2%<br>(92) | 0.2%<br>(165) | 0.3%<br>(163) | 0.3%<br>(154) | 0.3%<br>(33) |
| Archaic New | 1 -> 1 | 27%<br>(13885) | 27%<br>(15765) | 27%<br>(15411) | 27%<br>(14028) | 26%<br>(2845) |
| Archaic New | . -> 1 | 51%<br>(26003) | 53%<br>(31008) | 54%<br>(30546) | 55%<br>(28337) | 62%<br>(6878) |
| Archaic New | 1 -> 0 | 0.01%<br>(4) | 0.01%<br>(4) | 0.01%<br>(8) | 0.01%<br>(5) | 0.01%<br>(1) |

#### Accuracy of archaic variants vs MAF

Imputation accuracy has been shown to be higher for alleles with a high frequency in the reference panel^4^.

To assess the impact of MAF on the imputation of both archaic and non-archaic SNPs (independently from the classification of the region as archaic or non-archaic), we calculated the mean squared error between the imputed haplotype (using GP, see Supplementary Material) and the genotype in the original individual as a function of MAF in the 1000 Genomes. For non-archaic SNPs, our results are consistent with previous studies, showing similar values to those reported previously^4^ (Figure 3A). As expected, the error is higher in genomes downsampled at lower coverage values and for variants with lower MAF, confirming that both coverage and allele frequency significantly influence imputation accuracy. Strikingly, we observe a distinct pattern in the concordance of archaic SNPs (Figure 3A). Although slightly influenced by MAF at very low frequencies, these SNPs consistently show higher concordance than non-archaic SNPs. Even at very low coverage, the accuracy difference between archaic and non-archaic SNPs remains significant, particularly for variants with low MAF (Figure S10A). Notably, there is a significant drop in accuracy between the 0.5× and 0.0625× genomes. This explains why we were able to infer nearly all archaic segments detected in the original genome using the 0.5× imputed data, but only 50% when using the 0.0625× imputed genome.

**Figure 3.**
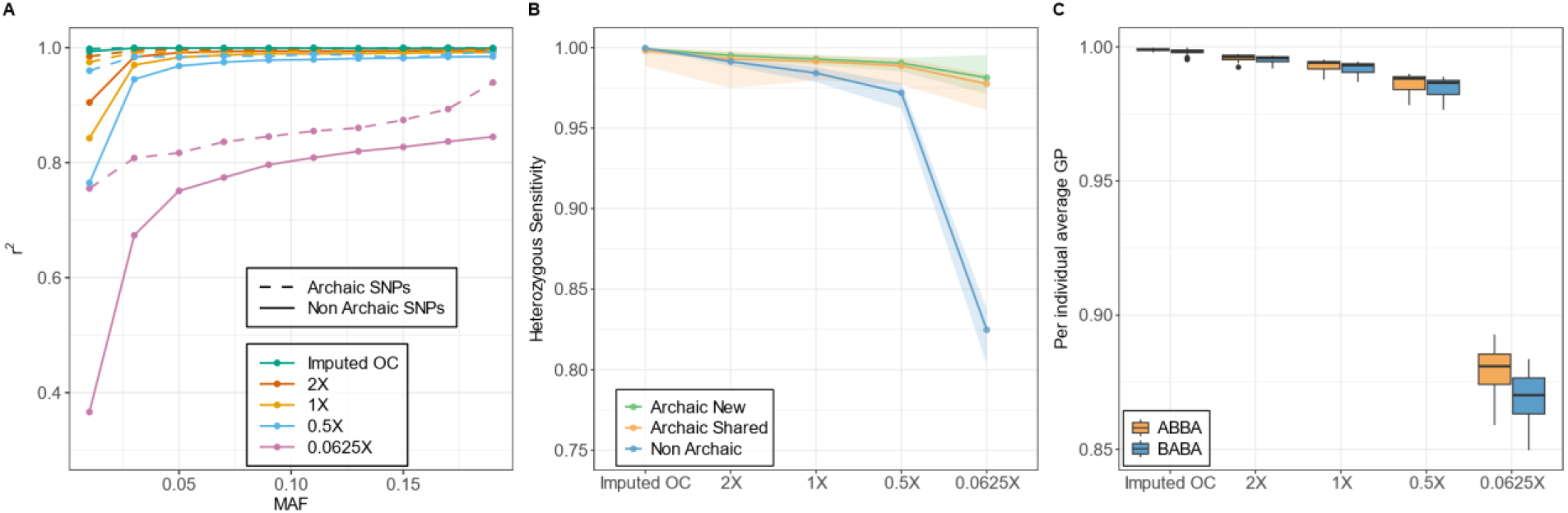
**A)** Imputation concordance for archaic and non-archaic SNPs across different downsampled datasets and Minor Allele Frequencies (MAF) in the reference panel. **B)** Mean heterozygous sensitivity in the imputed downsampled datasets, measured with Picard, across three categories: i) Archaic Shared— segments detected in both non imputed and imputed genomes; ii) Archaic New—archaic segments identified only in imputed genomes; iii) Non Archaic. Colored shades represent the minimum and maximum values. **C)** Per-individual average genotype probability for ABBA and BABA sites across different coverage levels, based on D-statistics of the form D(Africa, Ancient; Altai Neanderthal, Chimp). The individual Mota was excluded from all analyses.

When we compared the accuracy of archaic SNPs in inferred introgressed regions to their accuracy across the whole genome (Figure S10B), we observed that coverage and MAF has little to no impact at this level as the accuracy is consistently close to one. This indicates that introgressed segments are imputed with very high accuracy, and/or that LAI performs better in regions where imputation quality is high.

#### Heterozygous sensitivity in introgressed and non introgressed regions

We then expanded this analysis examining heterozygous sensitivity across all variants considering 3 groups of segments, Archaic Shared, Archaic New and Non Archaic. Once again, we observed higher sensitivity in archaic segments compared to non-archaic regions, with the difference between the two groups increasing as coverage decreases (Figure 3B). This trend reaches its maximum in the 0.0625× downsampled imputed dataset, highlighting that the accuracy of imputation in archaic segments remains superior to that in non-archaic regions, especially under more challenging conditions. We do not observe a major difference between the Archaic Shared and Archaic New.

#### Effects of imputation unbalance on the allele frequencies analyses

Building on these findings, we revisited the allele frequency analyses, where we previously observed an increase in *D* and *α* values when applying more stringent GP filters. To better understand this pattern, we examined how various factors—such as the genotype of the test individual, the ancestral alleles, and the amount of recoverable data—influence the *D-statistics* (Supplementary Material, Figure S10C). We observed that the average GP in ABBA sites (a proxy for archaic variants) is higher than in BABA sites (Figure 3C). Although this pattern is observed across all coverages, the difference becomes more pronounced at lower coverages, where imputation is inherently more challenging. Therefore, the increase in *D* values at low coverage and with high GP filter (Figure 1C, Filtered GP99) can be attributed to the fact that archaic variants are easier to impute than non-archaic variants, resulting in an enrichment of ABBA sites. This demonstrates that applying a GP filter—a standard quality control step in imputed data—can unintentionally introduce bias by over-representing ancestries that are more easily imputed, ultimately skewing the analysis.

### Use of archaic segments in ancient DNA

To further assess the quality of the identified archaic regions, we evaluated their utility in population genetic analyses by comparing them to contemporary datasets and ancient individuals to determine how well they capture population structure, introgression patterns, genetic affinities between ancient and modern populations, and changes in the frequencies of potentially adaptive introgressed regions over time and space.

#### Comparison with present-day populations

We looked in which contemporary populations we could find the archaic segments detected in our datasets, comparing, in a quantitative approach, all our detected archaic segments to the introgression maps from *hmmix*^*22*^ and *ArchaicSeeker2*.*0*^*24*^ papers. We found for all six datasets a clear and consistent relationship between the geographic origin of the individuals and the contemporary location of their archaic segments (Figures 4a, S11-13, Table S3). European Mesolithic and Neolithic samples share most of their archaic segments with contemporary Europeans, while non-European individuals like Kolyma1 and USR1 show greater sharing with East and Central Asian populations, consistent with Denisovan introgression (Figure 2D). These patterns remain consistent even at very low coverage (0.0625×), confirming that imputed genomes reliably capture geographic structure (Figures S11–S13). This supports the use of imputed introgressed segments to trace past migrations and the spread of archaic ancestry over time.

**Figure 4.**
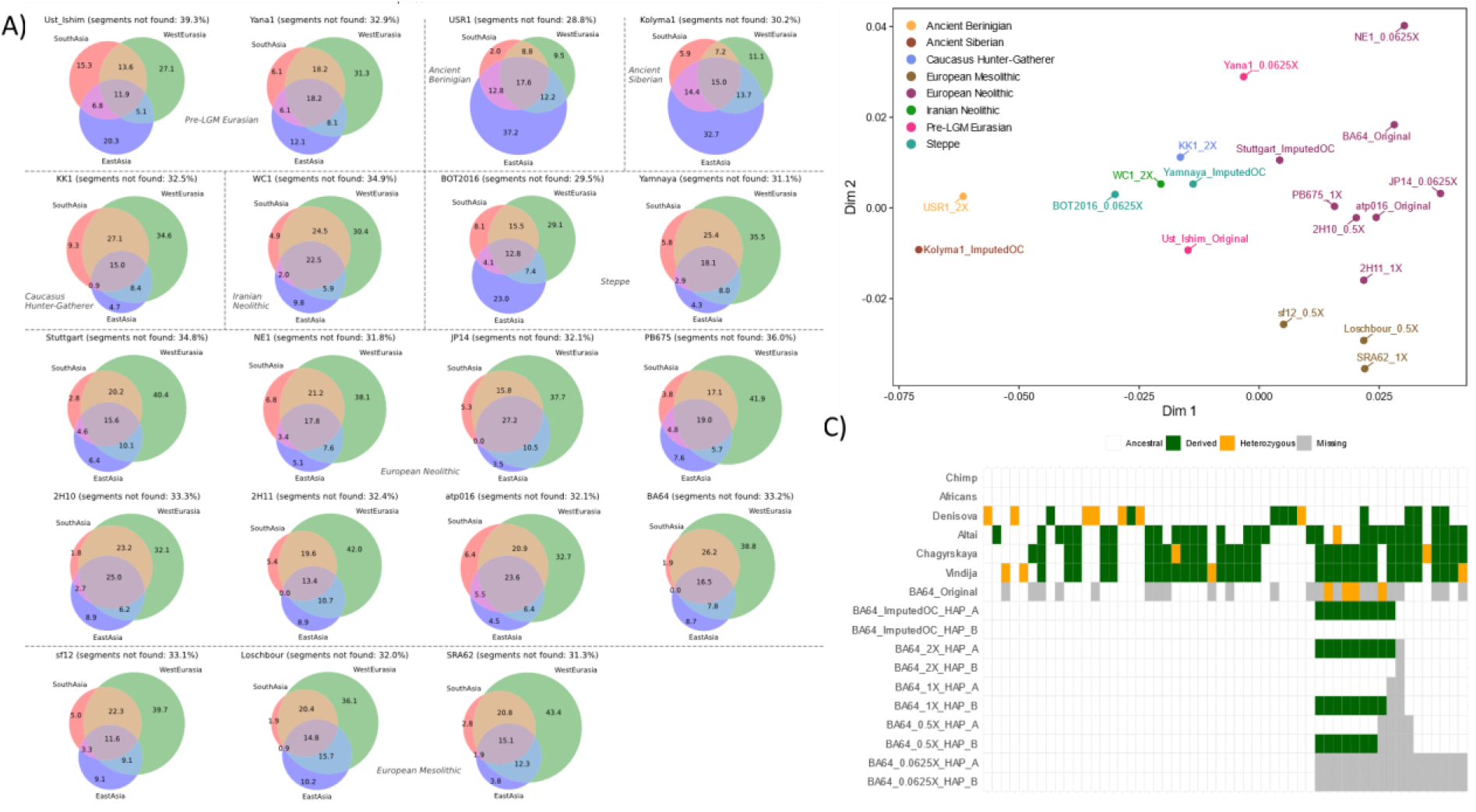
**A)** Venn diagram describing the proportion of shared archaic segments between imputed 1X ancient genomes and introgression maps from present-day individuals^22^ **B)** Multidimensional Scaling (MDS) analysis based on the count of introgressed segments among individuals from different imputed coverage datasets. **C)** Haplotype-level heatmap of archaic-derived SNPs (Table S7) in the BNC2 gene region across chimpanzee, African, archaic, and non-imputed and imputed BA64 genome after filtering for GP > 99%. Each row represents a haplotype or diploid individual, and each column corresponds to a possible derived allele. Alleles are classified as Ancestral (white), Derived (green), Heterozygous (orange) and Missing (grey). Imputed genomes are split into phased haplotypes (HAP_A and HAP_B), while archaic and original genomes are unphased.

Figure 4A shows that all ancient individuals - whether it is looking at original coverage, imputed 1× or imputed very low-coverage ancient individuals - have approximately 60% of their archaic segments shared with at least two groups (see Figure 4A, Figure S14-S15, Table S4). Notably, Ust’Ishim, who is the oldest individual considered here, shows one of the highest proportion of archaic segments unshared with any contemporary individuals, reflecting its ancient position in the Eurasian genetic landscape. Interestingly, Ust’Ishim’s exhibits similar proportions of sharing with European and East Asian populations. Unlike other individuals, Ust’Ishim lacks a strong affinity toward any specific geographic group, with only around 15% of his segments shared across all regions and more than 50% found exclusively in a single region (though not consistently the same one). A similar pattern is observed for Yana1—the second oldest individual of our dataset—although Yana shares a little more exclusively with Europeans (~24-31%, Figure 4A, S14). These results suggest that patterns of shared and unique archaic segments between Ust’Ishim and present-day population confirm the hypothesis that the population to which Ust’Ishim belonged diverged before the split of Europeans and East Asians^11^. Additionally, this suggests that differences in archaic introgression between Europeans and East Asians may not stem from multiple archaic sources but from ancient populations that once shared these variants^27^.

#### Comparison among ancient individuals

We examined shared archaic ancestry among the ancient individuals analysed in this work and found patterns consistent with those seen in modern populations, showing continuity between ancient individuals and their geographic regions. As expected, the oldest individuals do not preferentially share segments with any specific region, while European Neolithic and Mesolithic samples share more with each other (Figures S16-18). This pattern is robust across all imputed coverages, including the very low-coverage dataset, though slightly less pronounced at the lowest depths.

To mimic real-world scenarios in which datasets are composed of samples imputed from varying coverage levels, we created a mixed dataset of the 20 individuals analysed, each randomly selected from either the imputed or original datasets available in this project. Using an MDS based on shared segment counts in this dataset, we observed consistent population structure, comparable to that of uniform-coverage datasets (Figures 4C, S19-21). This structure aligns well with the geographic and temporal origins of individuals. Removing outliers (e.g., Mota and very low-coverage) further clarifies this structure (Figure S20-S21), revealing a distinct East–West pattern. Eastern individuals carrying Denisovan segments cluster together, while Neolithic western samples show proximity to Central Eurasian groups, reflecting their intermediate genetic profile. Overall, our results align with Iasi et al. 2024^23^, confirming that archaic segments alone can recapitulate population structure—here demonstrated using a quantitative approach.

#### Overrepresented regions and evolution of candidate adaptive introgression signals

After demonstrating that imputed ancient genomes can be used to study genome-wide patterns of archaic ancestry, we assessed imputation performance at the gene level. Focusing on 14 candidate genes for adaptive introgression in modern humans^25^, we detected archaic segments overlapping 9 of these genes in one or more individuals from one or more coverage datasets (Table S5). The number of carriers and the proportion of gene length covered by introgressed segments remain relatively consistent across coverages from 0.5× to imputed original (Figure 4c, Table S6). Notably, imputation increased gene recovery from 6 (in non-imputed data) to 9. The *OAS1/2/3* genes are less prevalent in the past compared to other genes. Among the genes analyzed, *TLR6-1-10* show high similarity to Neanderthal haplotypes and are found at relatively high frequencies in Europe since the Mesolithic (2 out of 3) and Neolithic (5 out of 8), with the exception of the ancient Siberian Kolyma1 individual. *BNC2* exhibits the highest frequency, particularly in post-LGM West Eurasians. It is found in 2 out of 3 European Mesolithic individuals and 7 out of 8 European Neolithic individuals, and outside Europe, it appears in Caucasus Hunter-Gatherers and Steppe populations.

We directly examined the derived alleles in *BNC2* (Table S7) and confirmed that the high proportion of missing positions prevents the identification of archaic introgression in the original genome even in the presence of derived alleles (Figure 4C). This suggests that even in genomes sequenced at relatively high coverage, imputing missing genotypes can help resolve haplotypes in specific regions of interest.

To explore the potential for identifying new adaptive introgression candidate genes—despite the limited number of individuals analysed—we examined regions statistically over-represented in the 11 European individuals included in our study. Significantly shared archaic segments were identified for each coverage level using z-scores of segment presence frequencies across individuals, with one-tailed p-values adjusted via Bonferroni correction. Regions with an adjusted p-value < 0.01 were considered over-represented and were expanded to include adjacent intervals with adjusted p-values < 0.05, to account for broader signals and the uncertainty in segment length typical of ancient DNA. These regions contain 17 protein-coding genes (Table S8), mostly grouped into two main clusters (Figure S22). The peak distributions are consistent across coverage levels, though statistical significance varies depending on the number of segments per dataset. Datasets with fewer segments—Original coverage and 0.0625X—may still reach significance due to missing data and reduced segment counts (Table S9). One of these two clusters, located on chromosome 15, contains genes already known to be introgressed, such as *SLC28A1* ^28^, and *WDR73. AKAP13*^*29*^ is significant in the original data, but not in the imputed data. Specifically, AKAP13 is present in 7 out of 8 Neolithic Europeans but only in 1 out of 3 Mesolithic individuals, failing to meet the minimum threshold for significance in imputed datasets.

Another cluster, located on chromosome 6 at the very beginning of the 6p21.3 band, is not present in the *Original* data. When analyzing the genes in the region, *LMED2* and *MLN*, as with *BNC2*, we observe the presence of archaic alleles (Table S10) in the Original data, but there is a substantial amount of missing data (Figure S23). In the imputed genomes, we are able to resolve this issue, retaining heterozygous sites in all samples except for individual sf12, who is homozygous in both the original and imputed data. The introgression is present in all the European individuals, except for Stuttgart. With imputation, we can shift the analysis to the haplotype level, where the introgressed haplotype is closer to the Vindija Neanderthal haplotype than to those present in the other high-coverage archaic genomes. This haplotype is completely absent in the 9 non-European individuals analyzed in this study (Figure S23B). These analyses suggest *LMED2* and *MLN* as promising candidates for adaptive introgression, that provide an example of the benefits of imputation.

## Discussion

In this study, we evaluate the effectiveness of inferring archaic ancestry from imputed low-coverage ancient genomes, showing a surprising higher effectiveness of the imputation in recovering archaic haplotypes compared to non-archaic haplotypes. This opens the possibility of using, in the context of ancient DNA, methods initially designed to study archaic introgression in high coverage genomes. Specifically, we downsampled 20 high-coverage (>10X) ancient genomes to four low-coverage levels (2×, 1×, 0.5×, and 0.0625×), imputed them as well as the original high coverage data, and compared patterns of archaic introgression — using *D-*statistics, *f4-ratio and hmmix* — between non imputed and imputed datasets. The results obtained with the imputed dataset were consistent across all five coverage values for *D- and f4-ratio* statistics (Figure 1, S1). The results on the imputed data are also comparable with the one obtained from pseudohaploidized data when using a transversion-only filter. We show that applying stringent genotype probability filters to the imputed data (GP > 99%) biases the *D-statistic* values and produces a slightly over estimation of the *α* at very low-coverage (0.0625X). We discover that this overestimation occurs because the quality of imputation of archaic variants is higher (higher GP) than non-archaic variants, especially at low-coverage (Figure 3C). To solve this problem, we extended the *D-statistics* to directly use GP as an input instead of the most likely genotype. This shows very stable results across coverages (Figure S2) and is a promising direction to adapt methods to imputed data. These results suggest that imputation can be effectively used in allele frequency analyses, providing an alternative approach to pseudohaploidization methods. Notably, unlike pseudohaploid data, imputation enables the use of all SNPs, rather than being restricted to transversions only.

Using *hmmix* to identify archaic segments, we find that we can identify nearly all archaic fragments in imputed individuals (that are present in the original individual data) even when we impute from as low as 0.5X and as much as 50% of the segments when we impute from 0.0625X (Figure 2B). Notably, we identify more archaic fragments in imputed individuals than in the original data, and we confirm the archaic origin of these “new” segments by looking at the similarity of the original genomes in these regions with the archaic references. We showed for LAI the importance of imputation in resolving some of the missing genotypes that make it harder to detect these segments in original genomes (Table1, Figure 4C). Moreover, we also checked the ability to identify Denisovan segments in Siberians and Arctic Americans in our sample, and inferred all the original segments and identify new ones in the imputed genomes (Figure 2). Despite the effectiveness of imputation, we acknowledge that it underperforms in reconstructing the full length of archaic segments, especially in the oldest samples (Ust’Ishim and Yana1), where the expected pattern of longer segments is reduced compared to original data (Figure S8). A similar bias is seen when comparing Archaic New and Archaic Shared segments (Figure S9). Therefore, we caution against using segment length to estimate the timing of introgression using imputed genomes, and future work accounting for this bias may lead to more accurate estimates of the length of introgressed fragments.

One surprising result is that imputation performs better in archaic versus non-archaic regions and variants, and imputation correctly assigns archaic variants within archaic regions and non-archaic variants within non-archaic regions (Table 1). Previous studies have shown that imputation accuracy improves with stricter GP filters (e.g., GP > 80%) and when focusing on common variants (MAF > 5%). We therefore tested the genotype accuracy inference in function of MAF as well heterozygous sensitivity—i.e., the imputation quality at heterozygous sites, which are typically more challenging to impute, we find that accuracy is higher for both archaic regions and archaic SNPs.

We also observed similar heterozygous sensitivity at both Archaic New and Archaic Shared segments, suggesting that SNP density in the original data has a reduced impact within archaic regions (Figure 3B). In sum, our results highlight the potential of imputation as a reliable approach for analyzing archaic introgression, especially in the growing number of ancient genomes available from recent studies.

Finally, we assessed the potential applicability of the archaic segments identified in imputed genomes in real-case scenarios. First, we compared the introgression maps generated in this study with those obtained from present-day populations in previous works^22 24.^Across all six datasets (one non-imputed and five imputed), we observed consistent geographic patterns: ancient European individuals share more archaic segments with contemporary Europeans, while Kolyma1 and USR1 align more with East Asians (Figure 4, S11-15). Ust’Ishim, as expected, shows no regional affinity and has a high proportion of unshared segments, supporting its placement before the West–East Eurasian split. This suggests that regional differences in archaic ancestry between Europeans and East Asians may have emerged from a shared ancestral pool rather than multiple admixture events. We found similar patterns of geographic structure among ancient individuals, stable across all coverage levels. To mimic real-world datasets, we assembled a dataset of 20 individuals, randomly selected from the 6 coverage datasets, and performed MDS on the shared segments (Figure 4B, S19-21). The resulting structure aligns with temporal and geographic origins, demonstrating that population structure can be identified using only archaic segments and confirming that imputation preserves population signals, even in mixed-coverage scenarios.

We also assessed whether imputation could help detect regions of overrepresented archaic introgression, potentially reflecting adaptive events. Of 14 candidate genes, 9 were recovered in imputed genomes—3 of which were not detected in the original data (Table S5). We show that Imputation helps to identify more archaic haplotypes, especially in Archaic New segments missed in original genomes due to lower SNP density (Figure 4C). Focusing on Neolithic and Mesolithic Europe, we identified two overrepresented gene clusters consistently across coverage levels, with significance varying by segment count. We provide an example of a gene region (e.g. *LEMD2* and *MLN*) of how imputation improves the detection of a putative selected archaic haplotype present in 10 out of 11 Europeans, absent in the non-European individuals, and most similar to the Vindija Neanderthal haplotype (Figure S23).

In summary, we show that using imputed genomes, we can obtain reliable measures of the introgression proportion, identify introgressed fragments, and that archaic ancestry recapitulates broad temporal and geographic patterns observed using all genetic variants. Imputation also helps resolve archaic haplotypes in ancient genomes making it a promising strategy to study fine-scale signatures of adaptive introgression through time. As the number of low-coverage ancient genomes rapidly increases, imputation will vastly reduce the amount of missing data, and as imputation works better in introgressed regions it will be useful to characterize archaic ancestry over time and space.

## Acknowledgments

M.R.C. and L.P. contributed equally to this study, and their names may be listed in either order as co-first authors. The authors thank David Peede for fruitful discussions that contributed to improving this study. This study was supported by the European Research Council (ERC), Horizon 2018 starting grant (ERC-2018-STG, 804994), by the Human Frontier Science Project grant (number RGY0075/2019) and by ANR-20-CE45-0010. M.R.C. was also supported by Taighde Éireann – Research Ireland under Grant number 24/PATH-S/12372.

## Author Contributions

M.R.C., L.P., and E.H.S. conceived and designed the study. M.R.C. and L.P. analyzed the data. M.R.C., E.M.B., L.O., and L.M.C. performed imputation. L.P. and E.H.S. conceived and developed the software.

M.C.A.A. and F.J. helped with the interpretation of results and provided feedback on the methodological approaches. This work was supervised by E.H.S. M.R.C., L.P., and E.H.S. wrote the manuscript, with input from all authors. All authors have reviewed and approved the manuscript.

## Conflict of Interest Statement

The authors declare no competing interests.

## Data and Code availability

The unfiltered imputed ancient genomes are available in Zenodo (add link)

The scripts we used to obtain the main results can be found in the following github repository: https://github.com/LeoPlanche/arcAncientImputed

## Notes

### Competing Interest Statement

The authors have declared no competing interest.

### Summary of Updates

We corrected Planche's first name and removed the high-quality figures from the bottom of the PDF to make it easier to access.

