## SupplementaryInfo for "Archaic ancestry inference in imputed ancient human genomes"

### Supplementary Material

*Material and Methods*

#### Dataset

Our strategy to assess the feasibility of using imputed ancient genomes to study archaic introgression in humans is based on a comparative approach, evaluating results from specific analyses using imputed downsampled genomes alongside those obtained directly from non imputed original genomes (Figure 1A). We selected and downloaded 20 previously published high-coverage (>10X in ancient DNA, aDNA) shotgun genomes (Table S1) from public repositories, representing ancient individuals from Eurasia, the Americas, and Africa, and spanning a broad temporal range—from approximately 44,000 to 5,000 years Before Common Era (BCE) (Figure 1B)[5–18].

The genomic data were realigned with *bwa*[30] to the same reference genome (*hs37d5*), and filtered for a mapping quality of 25 and a minimum read length of 34. These genomes were then downsampled to four lower coverage levels (2X, 1X, 0.5X, and 0.0625X) using *samtools*[31].
Genotype likelihoods were called using bcftools[31] for approximately 78 million biallelic SNPs from the 1000 Genomes Project Phase III[32], both for the downsampled and original genomes. These served as input for imputation with GLIMPSE[19], using the 1000 Genomes Project Phase III panel as a modern reference. Each individual was imputed independently and after imputation they were merged based on the coverage in five, one from each of the four downsampled coverages, plus imputation of the original data.
Various filters, such as Genotype Probability (GP) and Minor Allele Frequency (MAF) in the reference panel, were applied in different analyses to minimize potential biases introduced by imputation, as shown in previous studies (Sousa da Mota et al. 2023). Additionally, genotype likelihoods were called from the original genomes at the same SNP positions and filtered using a Genotype Quality (GQ) ≥ 30, Depth of Coverage (DP) ≥ 10, and Allele Balance ≥ 0.4. These high-confidence genotypes were treated as the truth set for comparison with the imputed data to evaluate imputation accuracy.
From the same original and downsampled BAM files, we also generated pseudohaploid calls at the same ~78 million positions using the *Haploid Calling* method implemented in ANGSD[33]. This allowed us to compare the imputation results with the most commonly used data type in low-coverage aDNA studies.

Other published data were also included in various analyses. The four available high-coverage archaic genomes—three Neanderthals (Altai, Vindija 33.19, and Chagyrskaya 8) and one Denisovan (Denisova 3)—were used as archaic references[34–37]. African individuals were represented by 292 genomes from the 1000 Genomes Project Phase 3, specifically from the Yoruba (YRI), Esan (ESN), and Mende (MSL) populations (Table S2). The chimpanzee genome (panTro6) was used as an outgroup.

#### Allele frequencies analysis - archaic Global Ancestry Inference

*D-statistics*

To identify archaic introgression, the most widely used approach is based on the *D-statistics* (also known as ABBA-BABA or Patterson's D-statistics[20]). This method compares the allele sharing between a test

individual or population and an archaic individual, relative to the sharing observed in an unadmixed modern population (Africans) with the same archaic individual.
To evaluate the ability to detect allele sharing between archaic and imputed genomes, we applied *D-statistics* using the qpDstat software from the AdmixTools package [20], in the form:
*D(Unadmixed modern humans, Test individual; Archaic individual, Outgroup).*
For this analysis, we used the Altai Neanderthal genome as the archaic reference, a set of 292 unadmixed African individuals from the 1000 Genomes Project Phase 3 (Table S2) as the modern reference population, and the chimpanzee genome as the outgroup.
The *D-statistics* were computed for the imputed, pseudohaploid, and original genotypes (Figure 1B, C, Figure S1). For the imputed data, we applied three different filtering strategies i) Raw (no filtering), ii) Filtered for GP ≥ 0.99, iii) Filtered for Minor Allele Frequency (MAF ≥ 0.5%) in the reference panel, following the approach of Sousa de Mota et al. 2023[4]. For the pseudohaploid data, we used two sets of variants: i) all SNP positions, ii) only transversions.
The transversion-only filter was applied to all samples, regardless of whether UDG treatment had been used in library preparation to mitigate post-mortem damage (PMD) effects.

Despite the overall consistency of results across datasets, a notable bias emerges when analyzing the very low-coverage dataset (0.0625×) filtered with GP ≥ 99%. In this case, we observed an inflated *D* value, indicating that stringent genotype probability filtering may distort allele sharing patterns at extremely low coverage. To address this bias, we implemented a method to compute the *D-statistics* directly from the GP given by imputation (Figure S2). For each position, with the GP of the imputed genome as ($GP_{0|0}$,$GP_{0|1}$,$GP_{1|1}$), its allele frequencies at the position is $GP_{0|0} + \frac{1}{2} GP_{0|1}$ and $GP_{1|1} + \frac{1}{2} GP_{0|1}$. The rest of the D statistics is computed as usual. The script used can be downloaded from github: https://github.com/LeoPlanche/arcAncientImputed/src/dstat.py.

To better understand the patterns that may bias the estimation of the *D-statistic* across different coverage levels, we measured the proportion of various features—such as total positions and genotype types (homozygous reference, homozygous alternative, and heterozygous)—across genotype probability (GP) bins of width 10%. In addition, we modified the RIPTA script (Peede et al. 2022) to extract the GP values associated with each ABBA and BABA site. By plotting how these patterns vary across GP bins—considering the average values across samples and the observed variability—we found a consistent trend across coverages, with the notable exception of the lowest coverage (0.0625x) (Figure S10C). In general, all features show increasing proportions with higher GP values. However, at 0.0625x, both the homozygous and heterozygous alternative genotypes tend to decrease in the highest GP bin (≥ 90%), while the proportion of homozygous reference genotypes (grey) and the number of positions increase.
Although both ABBA and BABA counts decrease at high GP values, the difference between them (i.e., the numerator of the *D-statistic*) increases slightly, whereas the denominator—defined as the total number of ABBA and BABA sites (ABBA + BABA)—decreases (Figure S10C). As a result, the ratio between the two, and thus the *D* value, tends to increase. This suggests that, overall, ABBA sites are more likely than BABA sites to pass a stringent GP filter.
To further investigate this, we calculated the average GP for ABBA and BABA sites per sample and coverage level (Figure 3C). We found that, across coverages, GP values were generally high, except at 0.0625x, where they dropped below 85%. Surprisingly, across all coverage levels, there was a consistent difference in GP between ABBAs and BABAs, with ABBA sites—representing the introgressed (archaic)

positions—showing higher genotype probabilities. This difference increased as coverage decreased, reaching its maximum at 0.0625x.
These results indicate that filtering on stringent GP thresholds can systematically favor ABBA over BABA sites, introducing a bias when comparing archaic and non-archaic SNPs.

Aside from the bias observed with a very stringent GP filter, we observe that the ancient African individual Mota—included as a negative control— consistently displays outlier positions in terms of *D* in each dataset, with values usually, but not always, closer to zero. However, when we apply filters for GP >99%, the *D* values tend to increase. If we apply the MAF filter in the reference populations, then *D*s stay close to zero with the exception of when Mota is imputed from the lowest coverage considered here (0.0625x) (Figure S1B). Also the value we consider the truth, computed directly from the original genome shows a deviation from 0, even if it is still an outlier in the original genome *Ds* distribution. Statistical significance of these signals, represented in Admixtools by the *Z-score* for each interaction, is less consistent. For example, we detect signals of introgression in Mota in some cases (Figure S1), and this may be due to the fact that Mota may actually have some archaic ancestry from back migrations from Europe or because imputation does not work as well in African individuals. In a previous study, it was shown that this African genome had the second lowest imputation accuracy among the ancient high-coverage genomes that were downsampled and imputed[4]. Also in this case we observe signals of introgression in Mota using the original data, using only transversion we do not see this signal anymore (data not shown). For all other interactions, the *Z-score* remains below |3.3|, allowing us to confidently exclude the presence of introgression in those samples.

*f4-ratio*
For global ancestry inference, specifically to estimate the proportion of archaic introgression in each individual genome, we applied the f4-ratio method from the AdmixTools package[20]. The level of Neanderthal introgression was estimated using the *f4-ratio* in the form: *f4(Altai, Chimp; Test Individual, Africa) / f4(Altai, Chimp; Vindija, Africa).* Across all coverage levels, we observed that applying a GP ≥ 99% filter tended to increase the estimated archaic proportion. For instance, while the average proportion in the original genome, imputed raw and pseudohaploid transversion data is around 2.5%, the estimates increase with stricter filtering in the imputed data. This effect is particularly evident as coverage decreases. For the 0.0625× dataset, the distributions of *f4-ratio* estimates with and without GP filtering showed minimal overlap, highlighting a clear discrepancy introduced by stringent filters at very low coverage. To assess whether the GAI correlates with sample-specific features and is independent of data processing variables, we tested correlations between α (*f4-ratio* values) and: i) the age of each ancient individual (in thousands of years BCE) and the original sequencing coverage (before downsampling).
For this analysis, the Mota individual was excluded.

#### Local archaic Ancestry Inference

For archaic local ancestry inference we use our custom version of *hmmix*[22] that is a reference free method, meaning that it does not require archaic genomes as references, but it does require an outgroup composed of individuals without archaic ancestry. We used the same unadmixed outgroup of 292 African genomes from the 1000 Genomes Project used in the allele frequencies analyses. This analysis was performed individually for each genome across all six datasets analyzed in this study—one corresponding to the original dataset and five to the imputed datasets. For the imputed data, we used

unphased genotypes filtered for GP ≥ 99%. The resulting introgression maps, which include both similarity scores and distances to archaic reference genomes (Vindija Neandertha and Denisova3, see below), are available at: https://github.com/LeoPlanche/arcAncientImputed/maps.
For Hidden Markov Models (HMMs) methods, missing information in one part of the genome does not affect the quality of inference in other distant regions with better coverage, missing data at one position affects inference only locally. This allows for the detection of archaic segments even in cases of very low-coverage data where there is a higher proportion of missing positions overall.
To estimate the proportion of the genome identified as introgressed, we summed the total length of introgressed segments, without any filters, for each individual and divided it by the length of the hg19 reference genome (Figure 2A). The amount of genome identified as introgressed depends on the downsampled coverage, with higher coverage resulting in a greater number of detected segments (Figure S4). Overall, we observed a probable overestimation of the proportion of introgressed genomes in some cases, which we then fixed using filters based on similarity to the archaic genome. To assess the recovery of introgressed regions in the imputed data, we compared the segments identified in the original non imputed dataset to those in each imputed dataset. We say that a segment found in the original dataset has also been found in an imputed dataset, if at least 80% of the segment overlaps a segment found with the imputed dataset. Across all individuals, we found that 100%, 99%, 99%, 97%, and 50% of the segments identified in the original data were also detected in the imputed original coverage, 2×, 1×, 0.5×, and 0.0625× datasets, respectively (Figure 2B).
Remarkably, even with the very low-coverage (0.0625×) imputed genomes, we were still able to recover half of the introgressed segments originally identified in the original data.

#### Similarity and distance to Archaic genomes

To evaluate the segments detected as archaic and non archaic for all datasets, we first compute their similarity to archaic reference genomes (Vindija Neandertha and Denisova3). The similarity for a segment is computed as such: we look at all the SNPs within the segment present in the archaic and in the sample's original genome. Then, there is a match if the archaic and the sample's original genome share an allele at this position. The similarity is the proportion of matches along all the SNPs. More precisely, let $a_1, a_2$ be archaic reference haplotypes and $s_1, s_2$- original sample haplotypes. Define the following sets of variable positions for a segment,

$$S = \{ \text{position } i : \exists k, j : a_k[i] = s_j[i] \}$$
$$\underline{S} = \{ \text{position } i : \forall k, j : a_k[i] \neq s_j[i] \}$$

Then similarity is simply the ratio,

$$\text{similarity} = \frac{|S|}{|S|+|\underline{S}|}.$$

By comparing the similarity of the original genomes in the regions identified as introgressed in the imputed genomes, we mitigate the potential bias introduced by imputation and confirm that the identified archaic segments exhibit higher similarity compared to the non-archaic ones (Figure 2C). Additionally, we classified the identified archaic segments into two groups: Archaic Shared for segments identified in both the imputed and original data (with overlap between the two segments), and Archaic New for segments identified only in the imputed data. Using this approach, we observed a higher similarity to the nearest archaic genome in the Archaic Shared segments compared to the Archaic New segments, which, however, still show greater similarity to the archaic genome than the non-archaic segments (Figure 2C).

We then also define a normalized version of the similarity. For a given segment, we compute the similarity as defined above between the sample's original haplotype and the archaic reference haplotypes, but also the similarity between individuals from Biaka in the HGDP dataset and the archaic reference haplotypes. Finally we compute the normalized distance as:

$$\text{distance} = \frac{1 - similarity}{1 - average\_similarity\_hgdp}$$

This approach offers a more principled way to post-processing classification of segments, returning a normalized distance between each ancient individual and the archaic reference genomes (Figure S5).
We also measured the similarity between the imputed genomes themselves (in the regions identified as archaic) and the archaic genomes. This allows us to assess how closely the actual imputed data used to build the introgression maps resemble the archaic references. Rather than solely confirming the absence of bias introduced by imputation, this similarity measure serves as a valuable metric for evaluating the quality of archaic segment calls. It can be extended to larger datasets of ancient individuals, beyond artificially downsampled genomes.

#### Filtering of introgressed segments

To reduce the possible bias/false positives and to keep only the candidate segments identified as introgressed in *hmmix* that have strong features of introgression, we proposed filters on the segments. We kept segments with a similarity to the closest ancient archaic genome (Vindija for Neanderthal and Denisova3 for Denisova) above 0.99 and with a length above 40Kb; that is above the length needed to reject incomplete lineage sorting for regions with moderate recombination rates[38]. With these filters we obtained results that are consistently close to the proportion measured with the *f4-ratio* (2-3%), for all coverages and individuals down to 0.5X (Figure S6), and in line with expectations, with the most ancient individual (on the left in Figure 2A) showing the highest proportion of archaic ancestry. Even with these quality filters the amount of introgressed segments that can be identified using imputed genomes is much higher than with the original non imputed data, and the variation among the different imputed coverage, with the exception of 0.0625x, is more stable (Figure S4).

#### Identification of Denisovan origin introgressed segments

We then assessed the potential of imputation to aid in the detection of Denisovan segments. This task is particularly challenging in ancient genomes when using a model that relies solely on the available high-coverage Denisovan genome[23], especially given the complex history of multiple Denisovan population introgression events[25].
We expect to identify Denisovan segments in the few individuals among those analyzed here who are not from West Eurasia or Africa. A segment was classified as Denisovan if it met three criteria: (i) it was detected as archaic with segment length of a least 100,000bp, (ii) it had a distance of less than 0.2 to the Denisovan genome (Figure S5), and (iii) its distance to Neanderthal was at least 0.3 greater than its distance to Denisovan. This very conservative filtering approach ensures high confidence in Denisovan segment identification, as our objective is to analyze the performance of imputation on genomic regions that can be reliably classified as Denisovan.
When analyzing the introgressed Denisovan segments inferred from the original dataset, we find, as expected, the highest number of candidate Denisovan segments in Kolyma1 (Siberia), followed by USR1 (Alaska) (Figure 2D, S7). This is consistent with the known ancestral history of these individuals, who, in addition to the Ancient North Siberian (ANS) ancestry, also have East Asian contributions[18]. In our

dataset, Yana1 represents ANS ancestry[18] and shows a limited number of Denisovan segments (Figure 2D).

We then compared the Denisovan introgressed segments we identified in ancient genomes with two introgression maps for contemporary individuals: one generated using *hmmix*[22], and the other using *ArchaicSeeker*[24]. All Denisovan segments, but one, detected in Kolyma1 were also identified in Skov et al. 2018[22], whereas only 20% of the segments found in other samples matched those in Skov et al. 2018[22]. When comparing them to the ArchaicSeeker introgression map, we found a match for all but two of Kolyma1's segments, and this time, 70% of the segments detected in other samples were also identified.

We then verified, as before, whether the introgressed segments obtained using imputed genotypes contained the Denisovan segments identified in the original dataset, as well as additional ones. Expanding the Archaic Shared and Archaic New categories to Denisovan Shared and Denisovan New.

Using imputed genotypes, we not only identified the segments present in the original dataset (at least down to 0.5× coverage) but also detected additional Denisovan segments (Figure 2D, S7). Most of these newly identified segments were also present in introgression maps based on present-day individuals. However, a small excess of segments was identified exclusively in the ancient DNA and was absent from both the original dataset and the Skov et al. introgression maps, which are based on 1000 Genomes Project individuals—the same dataset used for imputation.

These results demonstrate that imputation enhances the detection of Denisovan segments. It not only helps identify segments that persist in contemporary individuals but also enables the reconstruction of segments that were introgressed in ancient samples but are no longer present in the present-day reference panel. Although the analysis was done on a much smaller set of segments compared to all archaics and might require further investigation.

#### Quality of the introgressed segments

*Length*

The length of introgressed segments is expected to correlate with the age of the samples, with older individuals having longer segments, as they are closer to the original admixture events[25,26]. When examining segment length we observe that imputation has a deleterious effect, leading to ancient humans having values of length closest to what is expected in contemporary humans, i.e. 100,000-120,000bp for an introgression time 2400-2000 generations ago (Figure S8). Specifically, the two oldest individuals in our dataset, Ust'Ishim and Yana1, exhibit longer average segment lengths in the original dataset, +150,000bp and +74,000bp respectively compared to the more recent individuals. However, this trend diminishes in both Ust'Ishim and Yana1 when using imputed data (Figure S8). For individuals closer to the present, the differences in length between non imputed and imputed genomes is more similar, only differing by on average +9,722bp for Imputed OC, to -13,384bp for imputed 0.0625X. Considering the two groups, Archaic Shared and Archaic New, we observed that the decrease in segment length is more pronounced in the Archaic New group, while the Archaic Shared segments tend to be even longer (starting from 0.5X and above) compared to the original data. In contrast, the Archaic New segments make up the largest portion and are generally slightly shorter than the shared ones. To investigate whether this difference in segment length is associated with similarity to the archaic genome, we analyzed the closest archaic match (Vindija Neanderthal or Denisovan3) for each segment. Even in this case, the length and similarity distributions of the Archaic Shared and Archaic New categories remain quite similar (Figure 9B).

#### Accuracy of archaic alleles vs MAF

We assessed both how well imputation recovers archaic variants and how well it recovers archaic segments. The main question of interest is the ability of imputation to recover archaic segments, but the location of those segments is unknown and using the results of LAI methods in that context creates obvious biases. On the other hand we are able to define archaic variants independently of the ancient samples, eliminating such issues. A variant is defined as archaic if it is absent in African populations (Yoruba, Mende and Esan from the 1000 Genome Project) and homozygous in Vindija Neanderthal. Archaic SNPs are positions carrying these variants.

For each sample, we compared archaic positions between non imputed and imputed genomes (Table 1) in function of their state: non-archaic, Archaic New or Archaic Shared. In order for the raw numbers to be comparable, for non archaic, we picked a set of random non-archaic segments chosen to have the same length distribution as the archaic segments. We then looked at every SNPs in the genome and computed their imputation accuracy depending on if they were archaic (as defined in the previous paragraph) or not (Figure 3A). To do so, we directly use GP in the imputed genome and define a genotype between 0 and 2 as $GP_{1|1} + \frac{1}{2} GP_{0|1}$ and compute the mean square error to the original genotype at that position. We then plot 1-MSE, which is equivalent to $r^2$ correlation.

We also compared the imputation ability of the Archaic New and Archaic Shared groups, this time considering the results from the archaic LAI, using a different metric: heterozygous sensitivity. This metric measures the sensitivity to imputing heterozygous sites, which are common in archaic positions and the most challenging to impute. To assess heterozygous sensitivity, we used the GenotypeConcordance tool in Picard 3.0, comparing the original and imputed VCF files for each sample. In this case, we observed higher heterozygous sensitivity in the archaic segments compared to the non-archaic ones, with no significant differences between the Archaic New and Archaic Shared groups.

#### Analysis of Shared Introgressed Regions with Contemporary Individuals

For contemporary datasets we use *hmmix*'s map, containing individuals from Simons Genome Diversity Project (SGDP), 40 Papuans from [39] and 35 Papuans from [40] as well as *ArchaicSeeker2.0*'s map containing individuals from the 1000 Genome Project and Papuans from SGDP[41].

We look for each individual at the amount of archaic segments they share with each contemporary population. In *hmmix*'s map, contemporary individuals are assigned to one of four regions: East Asia, West Eurasia, South Asia, or Central Asia–Siberia. In *ArchaicSeeker2.0*'s map, individuals first belong to a local population as defined in the 1000 Genomes Project, which in turn belongs to a continental region: East Asia, West Eurasia, or South Asia.

To compute heatmaps (Figures S11-S13, Table S3), for each individual we count the number of their archaic segment that is found in each contemporary region. We consider that two segments are matching if they overlap for 80% of their length. If a segment is found multiple times within a region, we count it as many times as it is found. As within introgression maps some contemporary regions contain more individuals than others - especially in *hmmix*'s map - we then divide the match count for each region by the number of individuals in that region. Finally, we normalize by row.

To compute Venn diagrams (Figures 4A, S14-15, Table S4), for each individual and for each of their archaic segments, we look for each region if there are at least two matches in contemporary individuals present in *hmmix*' map, if so we mark that the segment is present in the region. If there is no match in any region, we mark the segment as not found. The Venn diagram is then the summary statistic for all archaic segments in the individual.

#### Analysis of shared introgressed regions across ancient individuals

To investigate patterns of haplotype sharing among ancient individuals, we applied a multi-intersection strategy to the set of introgressed fragments previously inferred by archaic local ancestry analysis. For each coverage level, segments were grouped by chromosome and divided into discrete, non-overlapping genomic intervals defined by unique start and end positions (breakpoints) across all individuals. This allowed us to standardize the genomic coordinate system and create binary matrices recording the presence or absence of archaic fragments per individual in each interval, analogous to the multiintersect function in bedtools. Adjacent segments that were contiguous on the genome and shared by overlapping individuals were merged into larger blocks to reduce redundancy.

Using this collapsed matrix, we computed a pairwise segment-sharing matrix by counting the number of introgressed regions shared between each pair of individuals. This matrix was then row-normalized to represent, for each individual, the proportion of their introgressed segments shared with all others.

The resulting matrix was visualized as a clustered heatmap using the pheatmap package in R. Row clustering was based on Spearman correlation with Ward's linkage, and the same order was applied to columns to ensure symmetry. Population labels were added as annotations, and matrix values were displayed as percentages representing the proportion of shared introgressed segments relative to each row individual's total (Figure S16-18).

To ensure comparability across coverages and reduce sampling bias, we randomly selected a balanced subset of individuals from each coverage level. For this subset, we computed the total length of introgressed fragments shared between every pair of individuals. Shared length was defined as the sum of minimum overlapping lengths across all intervals where both individuals carried a fragment. To account for differences in the total amount of introgressed sequence per individual, we normalized the resulting pairwise matrix row-wise by each individual's total introgressed length. This normalization reflects, for each individual, the proportion of their archaic segments that are shared with others, providing a more interpretable measure of relative haplotype similarity. To visualize population structure and shared ancestry patterns based on archaic content, we applied multidimensional scaling (MDS) to the normalized distance matrix derived from the pairwise sharing values (Figure 4B, S19-21). The resulting two-dimensional coordinates were used to generate scatter plots, in which individuals were color-coded by population and annotated by sample. This allowed us to explore clustering patterns and potential affinities among individuals based on the structure of their shared introgressed genome. These plots highlight both inter-individual similarity and broader group-level trends, offering insight into the extent to which archaic fragments are shared across populations and coverage levels.

#### Analysis of Known Candidate Genes

To assess the presence of archaic introgressed fragments at previously proposed candidate loci, we curated a list of 14 genes of interest[25] and retrieved their genomic coordinates. Coordinates were mapped to the GRCh37 reference genome, and gene boundaries were extended to include their full transcriptional span. Using Ensembl BioMart, we confirmed the identity of overlapping protein-coding genes within these intervals. We then intersected these gene regions with archaic fragments previously identified through LAI, after quality filtering. This allowed us to determine whether individual fragments overlapped candidate genes, and to compute per-sample, per-gene summary statistics (Table S5). For each gene, we recorded the average similarity and genetic distance to the archaic references across all overlapping fragments, stratified by coverage level and sample (Table S5-6). Genes with no detected overlap were retained in the final output to distinguish true negatives from missing data. Results were summarized in a table indicating which individuals carried archaic segments at each gene, and how often such overlaps occurred across different sequencing coverages.

We extracted a subset of informative SNPs from the *BNC2* locus based on allele state comparisons among archaic, African, and chimpanzee genomes. Using vcfR, we read phased (imputed genotypes after filtering for GP > 99%) and unphased VCF files for archaic individuals (Altai Neanderthal, Vindija Neanderthal, Denisova, Chagyrskaya), African samples (YRI, ESN, MSL), and a chimpanzee reference genome (panTro6). SNPs were retained if they fulfilled two conditions: (i) at least one archaic genome carried the derived allele, and (ii) all African samples were homozygous for the ancestral allele. Ancestral and derived alleles were inferred based on chimpanzee genotypes (Table S7). We then extracted genotypes for these filtered sites from the original ancient genomes and in the five imputed datasets for the individual BA64. Genotypes were converted into haplotype labels ("Ancestral", "Derived", "Heterozygous", "or "Missing") based on their match to the inferred allele states.

To visualize haplotype structure, we constructed a matrix where each row represented a haplotype or diploid individual and each column a filtered SNP. In the figure each allele states is encoded by color. Sample rows were grouped by coverage and origin (chimpanzee, African, archaic, original non imputed, imputed) to allow comparison across conditions (see Figure S4C).

#### Selection Scan on Imputed and Non Imputed European Genomes

To investigate signals of selection in introgressed regions, we conducted a genome-wide scan for overrepresented archaic fragments in ancient (Mesolithic and Neolithic) European individuals for the original genome and the five imputed datasets. We focused on segments previously identified with the LAI as likely archaic in origin after filtering for the length (>40Kb) and the similarity to reference Neanderthal and Denisovan genomes above 0.99. Introgressed fragments were projected across the genome and divided into discrete non-overlapping intervals based on their breakpoints. For each coverage level, we quantified the frequency of archaic segment presence across individuals, producing a binary presence/absence matrix per genomic interval, like the output of multiintersect in bedtools. Z-scores were computed for each interval to measure deviations from the mean segment-sharing frequency, and one-tailed p-values were calculated to identify intervals with significantly elevated archaic segment sharing. Bonferroni correction was applied to account for multiple testing. We retained regions where at least one interval showed significant enrichment (adjusted p-value < 0.01) and subsequently expanded these to include adjacent intervals with suggestive evidence (adjusted p-value < 0.05), allowing for the identification of broader genomic regions potentially affected by selection and due to the length uncertain that there is with ancient DNA. This "anchored" expansion strategy aimed to capture extended signals beyond single isolated peaks. To annotate the candidate regions, we queried the Ensembl database (GRCh37) and identified overlapping protein-coding genes. We tracked whether genes overlapped "core" significant regions (strict threshold) or extended regions (anchored by core segments), and assessed coverage-specific recurrence of gene hits (Table S8-9). Finally, we summarized the presence of these genes across coverage levels and visualized the results in Manhattan plots with Z-scores and gene annotations stratified by coverage (Figure S22).

We then directly tested the genotypes of one of the two clusters where we found a significant overrepresentation of archaic segments in Europeans. Since the cluster on chromosome 15 contains genes already identified as candidates for adaptive introgression, we focused on the cluster on chromosome 6, as this region was not significant in the original dataset. We concentrated on the region from the beginning of the *LEMD2* gene and the end of *MLN*. This region is located in the 6p21.3 band, which includes the MHC, a region known for its very high variability. As for *BCN2*, we reconstructed a heatmap showing positions where Africans are homozygous for the ancestral allele (defined using the Chimp genome), and at least one of the four high-coverage archaic genomes has a derived allele. We identified 20 positions, of which only three are not present in the 1000 Genomes Project strict mask (Table S10). In the plot, the chimpanzee and African rows represent the ancestral allele, while the archaic high-coverage and original non-imputed genotypes are colored based on the presence of the ancestral (white), derived (green), heterozygous (yellow), and missing (grey) alleles. If the cell is green or white, it indicates homozygosity for the derived or ancestral allele, respectively. Next, we plotted phased imputed genomes from European and non-European individuals. Each row represents a haplotype (single allele, not the genotype), so when an individual is heterozygous, both haplotypes are considered. What we observed (Figure S23A) is that in the original data, despite a large number of missing positions, all European samples have at least one derived allele, except for Stuttgart. Only the individual sf12 is homozygous for the derived allele. We observed the same situation in all imputed datasets with coverage up to 0.5X, with the missing positions being resolved as haplotypes from Vindija Neanderthal, but not from the other archaic humans. All samples were heterozygous, except for sf12, and no derived alleles were present in Stuttgart. This means that 10 out of 11 Europeans in our dataset show a derived allele in the original data, which we can resolve as an archaic haplotype in the imputed data. In contrast, when considering the non-European individuals (Figure S23B), none of them has a derived allele in the original data or an archaic haplotype in the imputed genomes. This illustrates that without imputation, this signal is not significant, primarily due to missing positions. However, with imputation, we can detect the signal and reconstruct the haplotypes.

*Supplementary Figures*

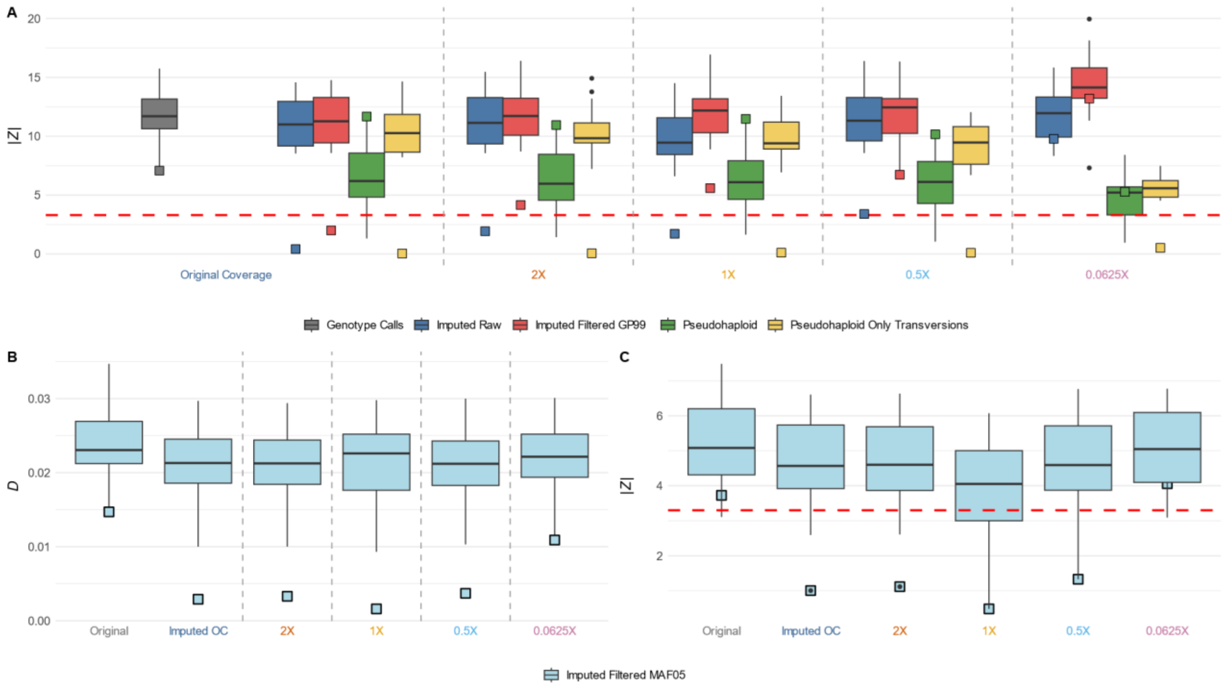


**Figure S1.** (A) Absolute Z-scores (|Z|) for different data processing methods and coverages, with highlighted "Mota" samples (black outlined points). The red dashed line represents the |Z| threshold of 3.3. (B) Distribution of D-statistics across coverages after filtering for Minor Allele Frequencies of 5% in the reference panel. (C) Absolute Z-scores (|Z|) after filtering for Minor Allele Frequencies of 5% in the reference panel, with "Mota" samples highlighted (black outlined points).

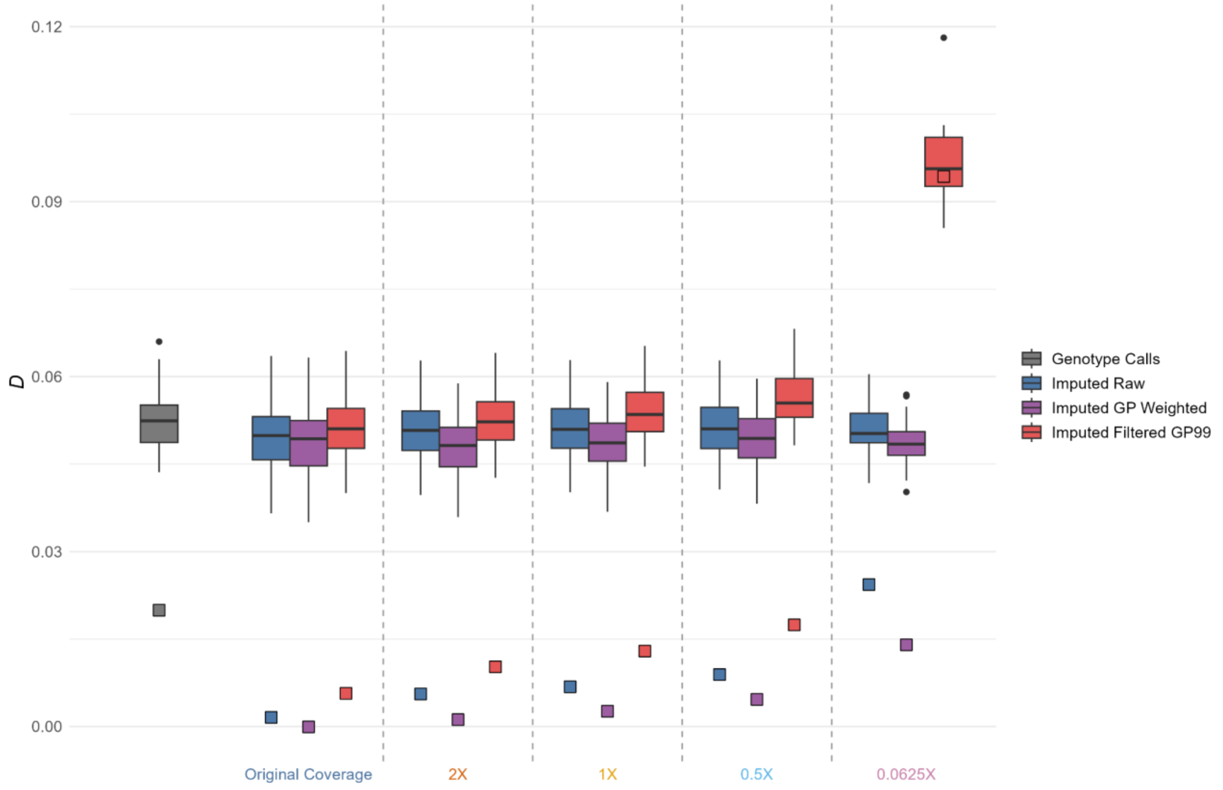

**Figure S2.** D values obtained using the new developed tool to include the Genotype Probability in the calculation of the allele frequencies.

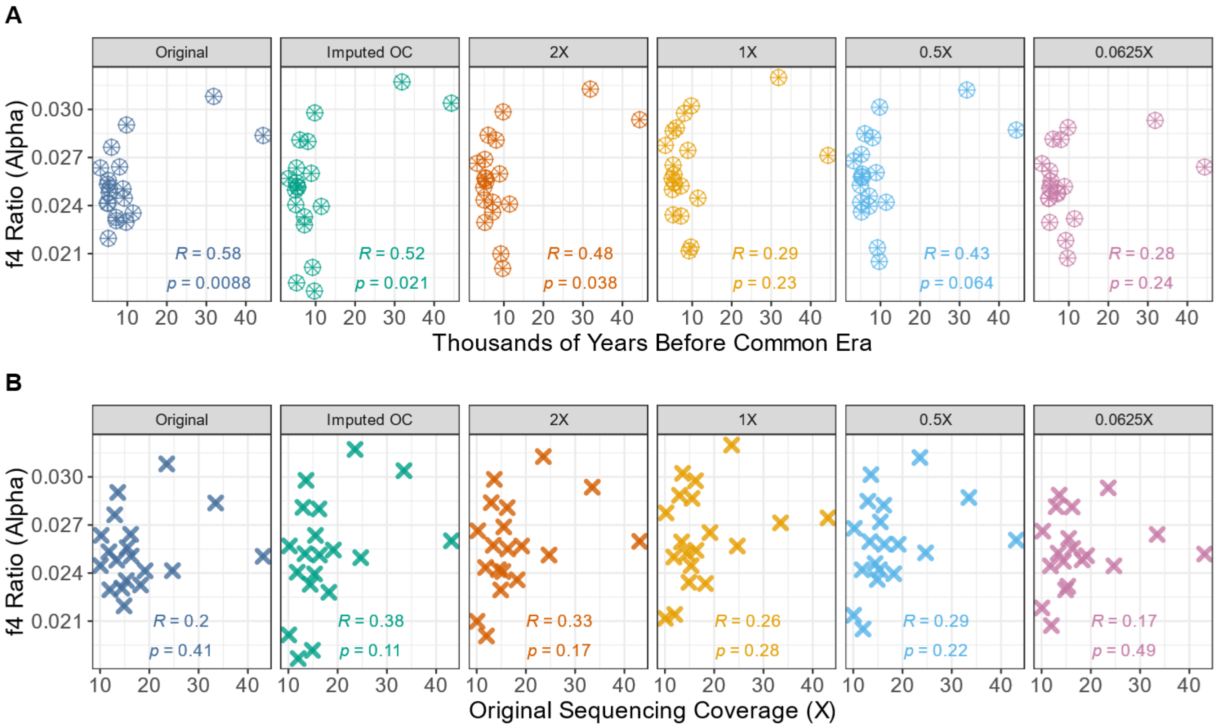


**Figures S3.** Relationship between the archaic proportion ($\alpha$) and A) The age of the ancient individual in Thousands years before common era, B) Coverage of the original sequencing coverage. For each of the comparisons the 6 different datasets have been analysed independently. The Pearson correlation coefficients are annotated on each plot. For this analysis, the Mota individual was excluded.

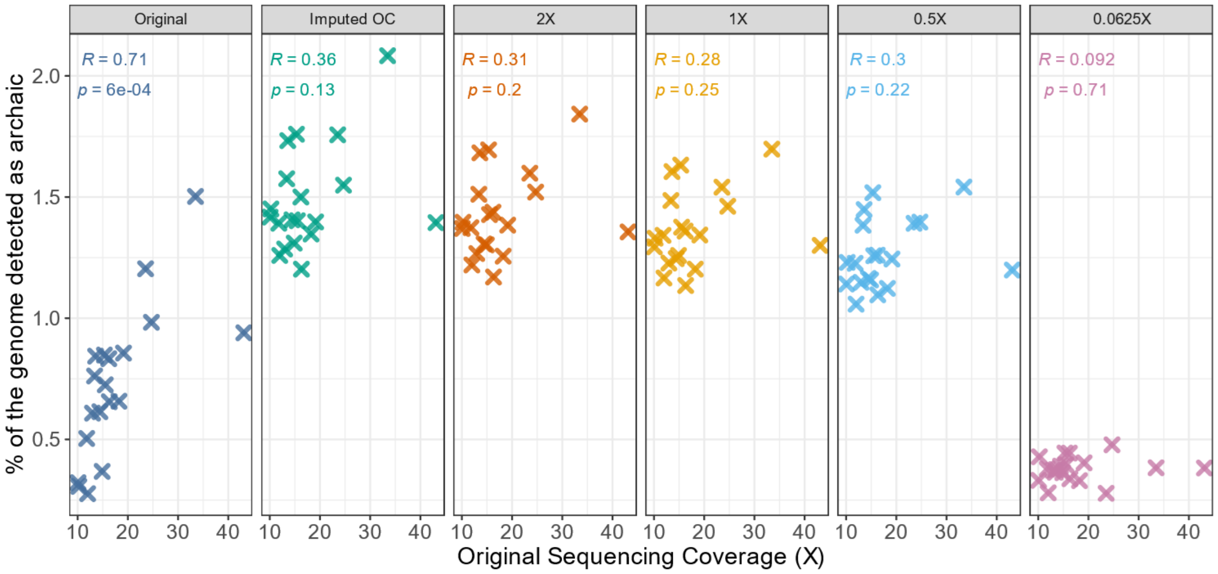


**Figures S4.** Correlation between the amount of archaic segments inferred for each individual in each dataset and its coverage. The Pearson correlation coefficients are annotated on each plot. For this analysis, the Mota individual was excluded.

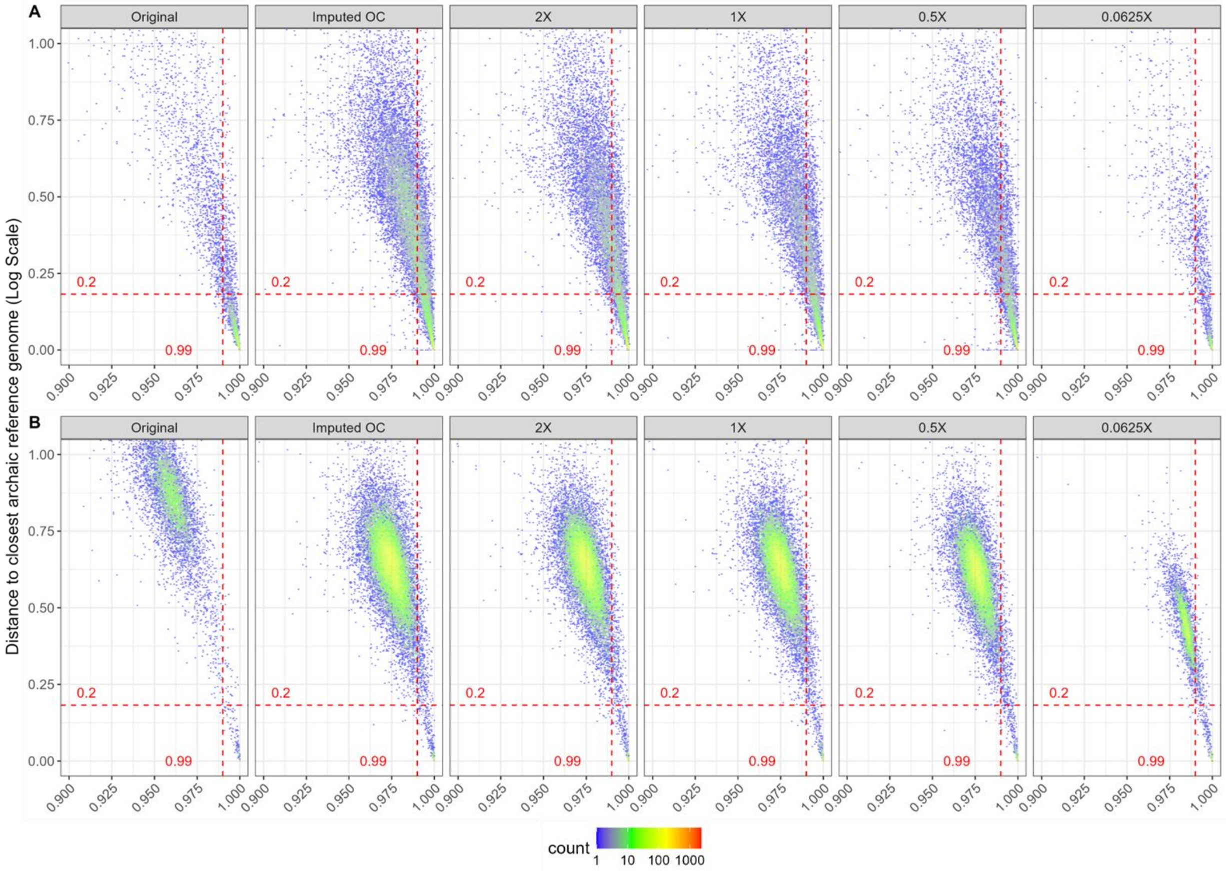


**Figures S5.** Density plots show the relationship between similarity to the closest archaic reference genome (x-axis) and distance to the closest archaic reference genome (log-transformed, y-axis) for (A) archaic segments and (B) non-archaic segments. Each hexagon's color reflects the log-scaled number of segments per bin. The vertical dashed red line marks the 0.99 similarity threshold and the horizontal line marks a distance of 0.2 used in the filtering of denisovan segments.

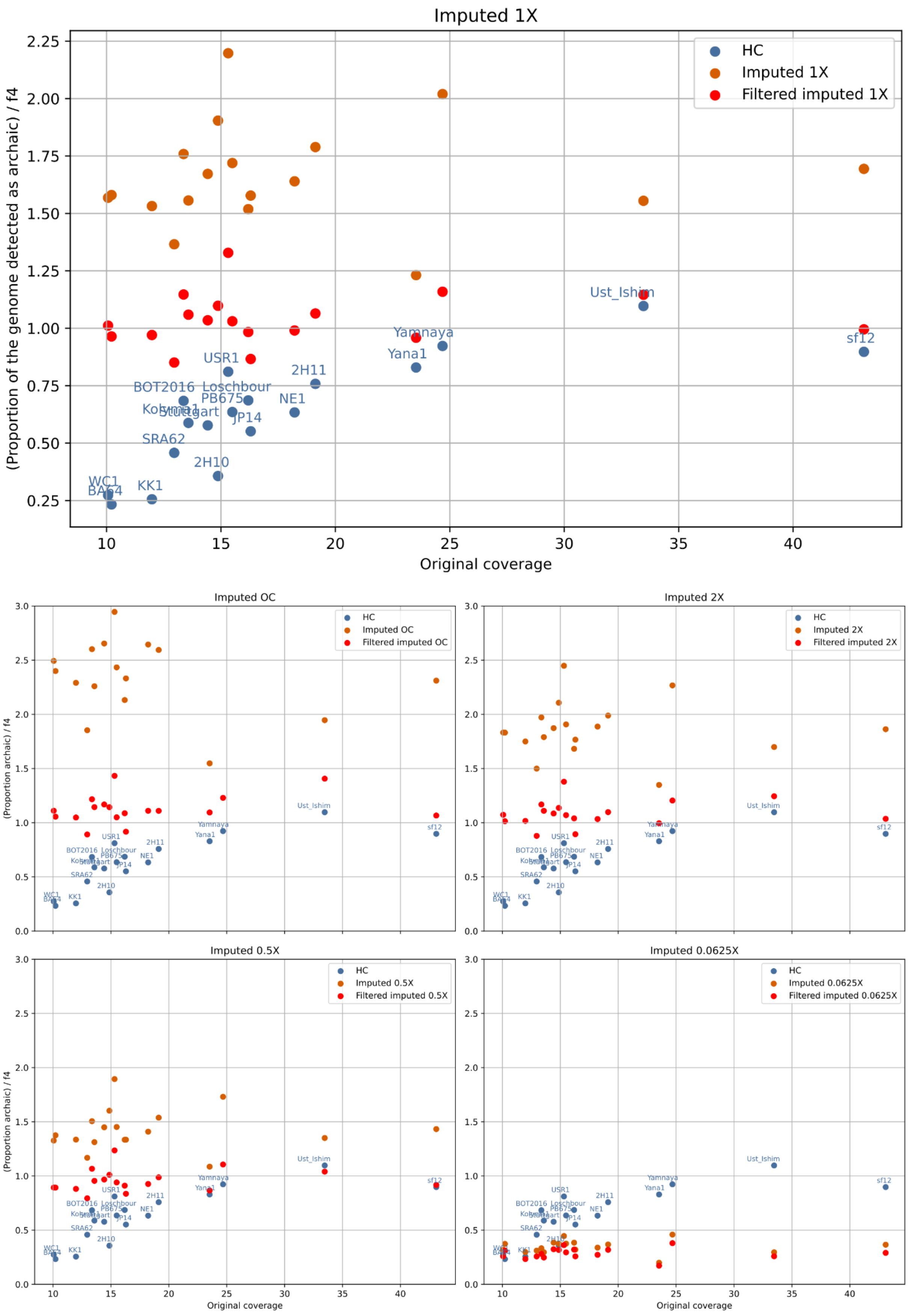


**Figure S6.** For each sample, coverage, we compute the proportion of the genome detected as archaic by summing the length of segments inferred as archaic and dividing by total genome length of

3,000,000,000. We then divide this proportion by the f4 ratio computed for the same sample. As both these values should correspond to the proportion of archaic ancestry, we expect the ratio to be 1. For the non imputed samples (blue dots), we see that this ratio depends on the original coverage of the sample, with coverage below 20X, the LAI method does not infer all archaic segments and the ratio is below 1. For imputed data (red dots), the ratio is above one for most samples, except for imputed 0.0625X, hinting at the presence of false positives. After applying filters of similarity to archaic genome greater than 0.99 and minimum segment length of 40kb (orange dots), we see that the ratio is close to one - except for imputed 0.0625X - for all individuals, independently of the original coverage,, showing that imputation can improve the inference of archaic segments even when the original coverage is already considered good for ancient DNA, i.e 10-20X. For imputed 0.0625X, we get after filtering an average ratio of 0.29, showing that we recover a significant portion of archaic segments even at such low coverages.

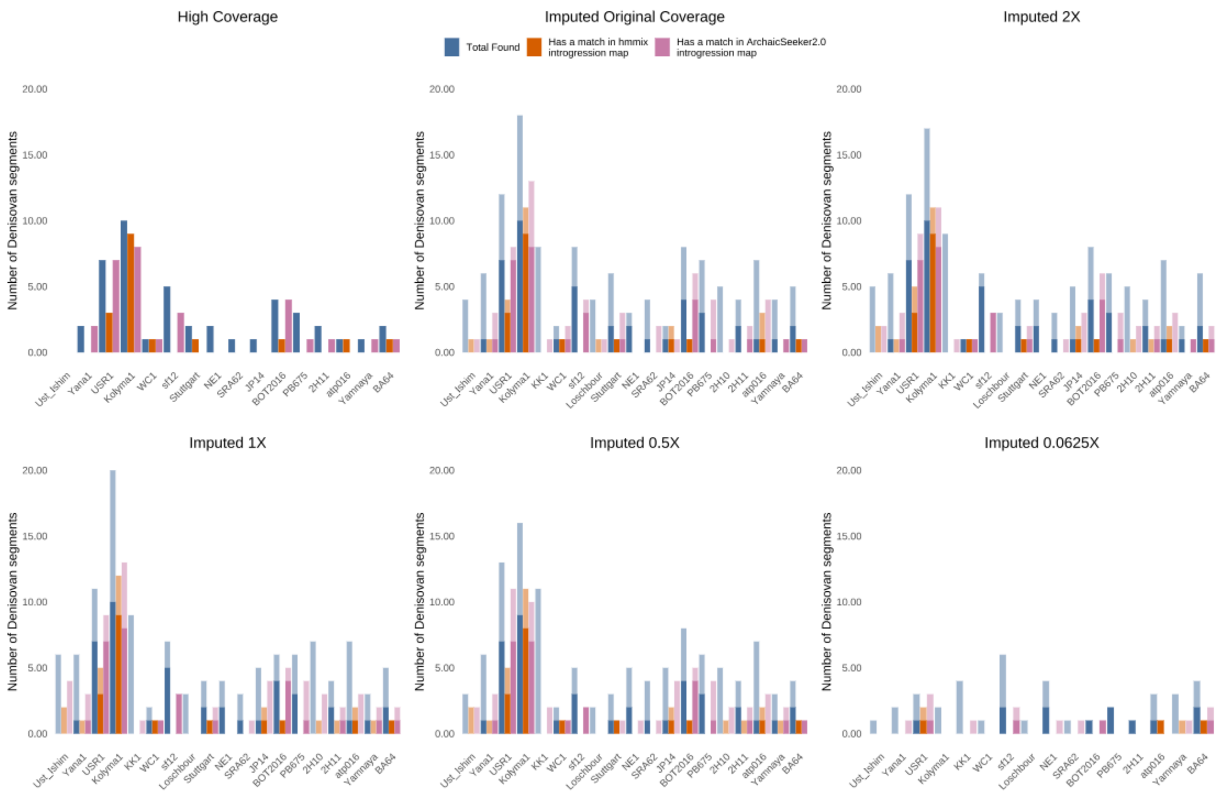


**Figures S7.** Number of Denisovan segments found in each dataset. We apply very stringent filters to classify a segment as Denisovan: minimum length of 100,000bp, distance to Denisova 3 below 0.2 and distance to Vindija Neanderthal 0.3 or more than its distance to Denisova. In blue the total number of denisovan segments found, in orange the amount of these segments also found in the *hmmix* introgression map, in pink the amount found in ArchaicSeeker 2.0 map. The light shade corresponds to Archaic New segments, the dark shade, Archaic Shared.

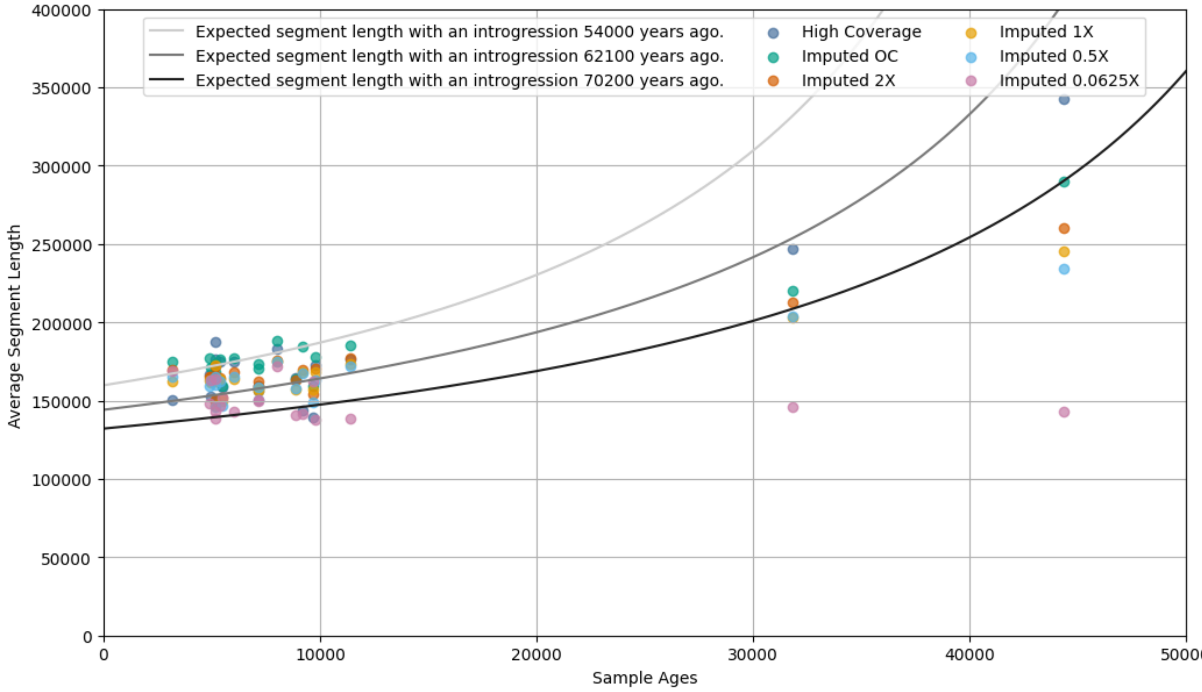


**Figure S8.** Average archaic segment length depending on the dataset used. For segments with an age below 12,000 years (all individuals but Yana1 and Ust Ishim), the average difference in length between the original coverage and imputed OC, imputed 2X, imputed 1X, imputed 0.5X and imputed 0.0625X segments are respectively +9,722bp, +988bp, -1,711bp, -3,635bp and -13,384bp. For Yana1 (31,850 years old), the differences are respectively -26,222bp, -34,241bp, -43,863 bp, -42,482bp and -100,586bp. For Ust Ishim (44,366 years old), the differences are respectively -52,731bp, -82,396bp, -97,524bp, -108,406bp and -199,563bp. For purely illustrative purposes, we give the expected segment length for three different introgression times, considering a constant recombination rate of $2.3*10^{-8}$, 27 years per generation and conditioned on a minimum segment length of 40kbp, corresponding to the filter we apply to the inferred archaic segments.

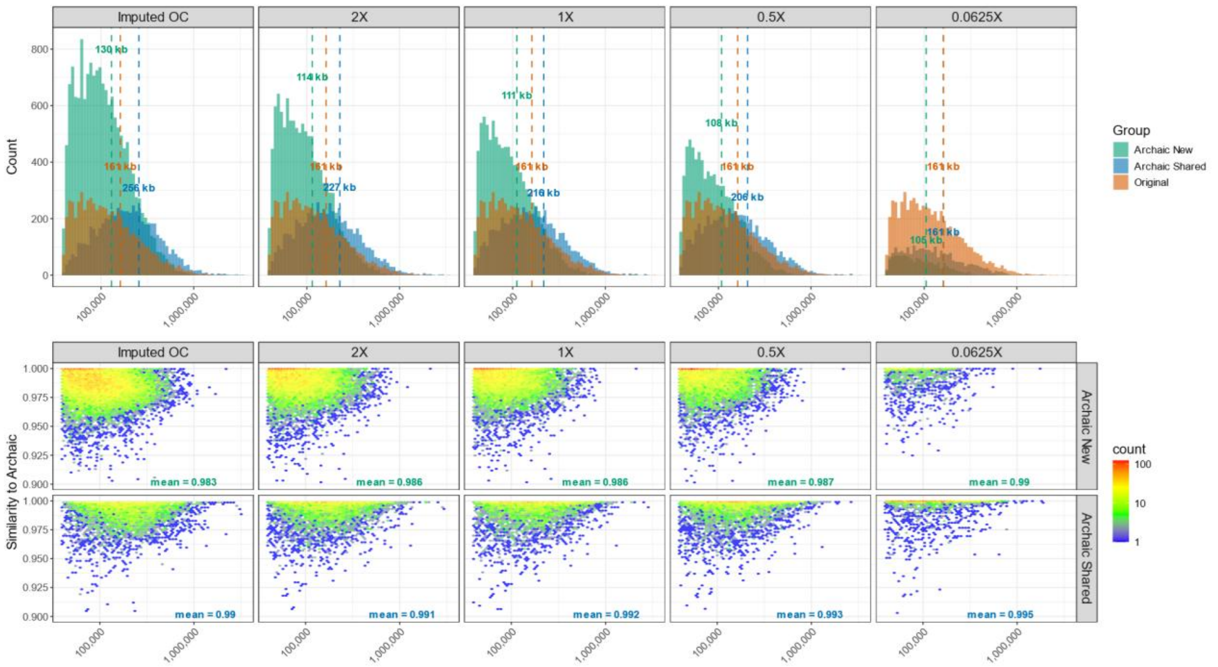


**Figure S9.** A) Histogram of archaic segment lengths (log-scaled) across different imputed coverages. Each imputed group is compared against overlaid distributions of original coverage segments. Dashed vertical lines indicate the mean segment length per group, with corresponding mean values annotated. B) Hexbin density plots showing the relationship between segment length (log-scaled) and similarity to the closest archaic genome for each imputed coverage, separately for Archaic New (top) and Archaic Shared (bottom) segments. Mean similarity values per group are shown within each facet.

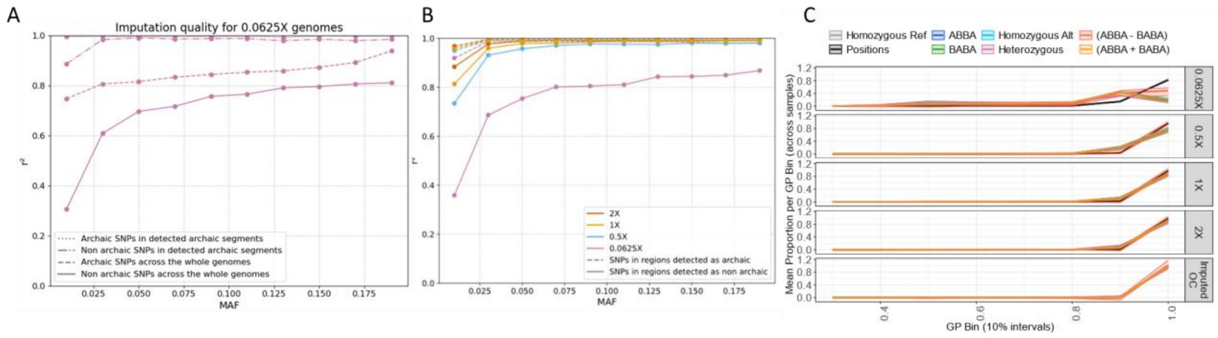


**Figure S10.** Imputation accuracy (r²) as a function of minor allele frequency (MAF). A) Imputation performance for the 20 samples downsampled to 0.06× coverage, shown separately for different SNP categories: Archaic SNPs and Non-Archaic SNPs, in both archaic regions and across the whole genome. B) Imputation accuracy across different coverage levels for SNPs located in Archaic and Non-Archaic regions. C) Distribution of site and genotype characteristics across genotype probability (GP) bins at different coverage levels. The figure shows the mean proportion (solid line) and range (shaded area) of various patterns—including genotype categories (homozygous reference, homozygous alternative, heterozygous), ABBA/BABA signals, and total positions—across GP bins of width 0.1, stratified by coverage. Each curve represents the average across samples.

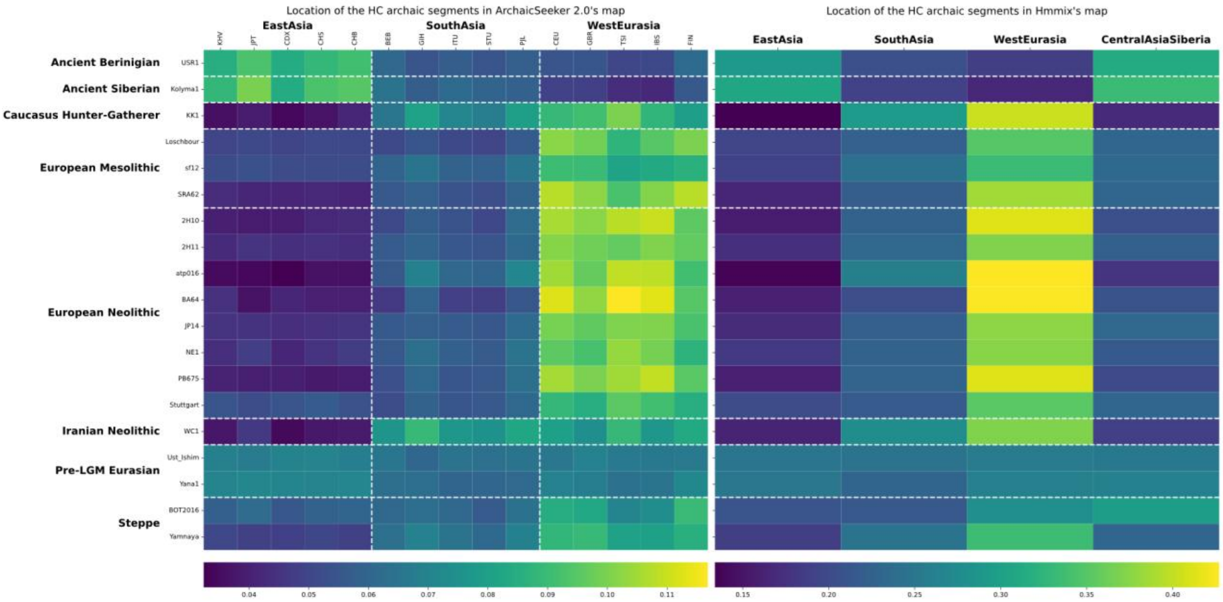


**Figure S11.** Geographical location of the archaic segments detected in the original coverage genomes within introgression maps of contemporary individuals. For each archaic segment detected in the imputed genomes, we count the number of overlaps with previously published introgression maps from the HGDP and 1000 Genomes Project. We then divide this count by the total number of segments of the same origin in the introgression map. Finally, the values are row-normalized.

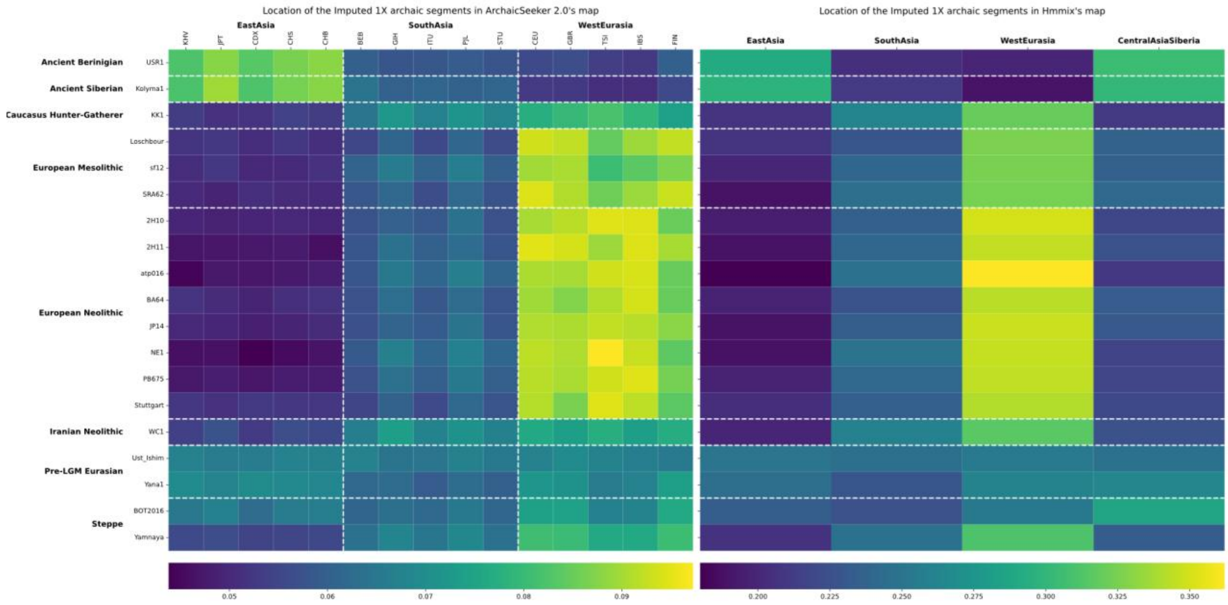


**Figure S12.** Geographical location of the archaic segments detected in the imputed 1X genomes within introgression maps of contemporary individuals. For each archaic segment detected in the imputed genomes, we count the number of overlaps with previously published introgression maps from the HGDP and 1000 Genomes Project. We then divide this count by the total number of segments of the same origin in the introgression map. Finally, the values are row-normalized.

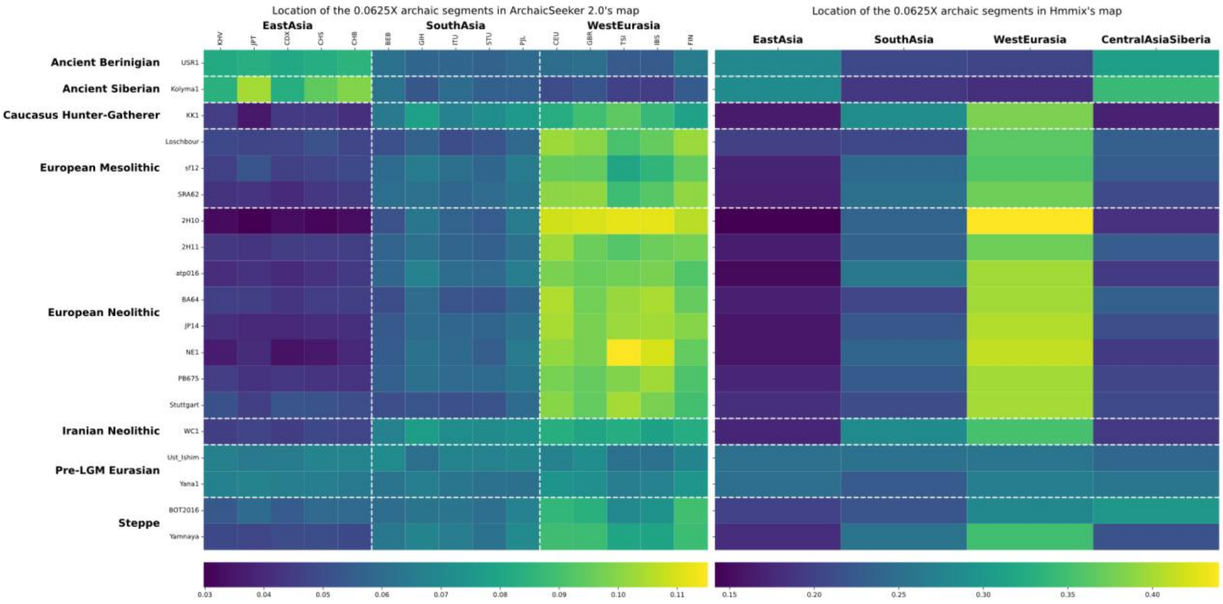


**Figure S13.** Geographical location of the archaic segments detected in the imputed 0.0625X genomes within introgression maps of contemporary individuals. For each archaic segment detected in the imputed genomes, we count the number of overlaps with previously published introgression maps from the HGDP and 1000 Genomes Project. We then divide this count by the total number of segments of the same origin in the introgression map. Finally, the values are row-normalized.

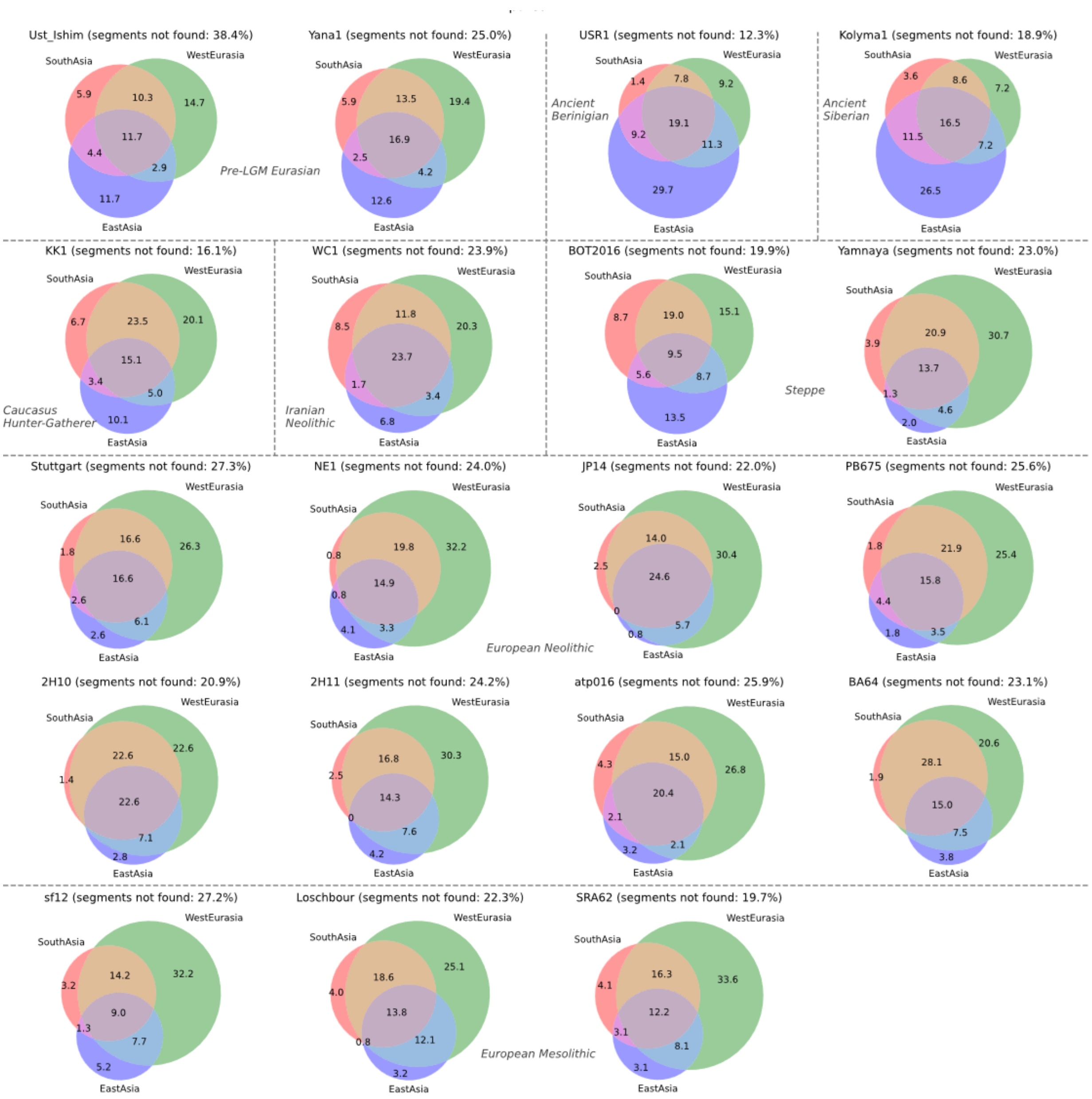


**Figure S14.** Proportion of the inferred archaic segments detected in the original coverage genomes that are uniquely found in West Eurasia, South Asia, East Asia regions, using contemporary individuals in[22] introgression maps.

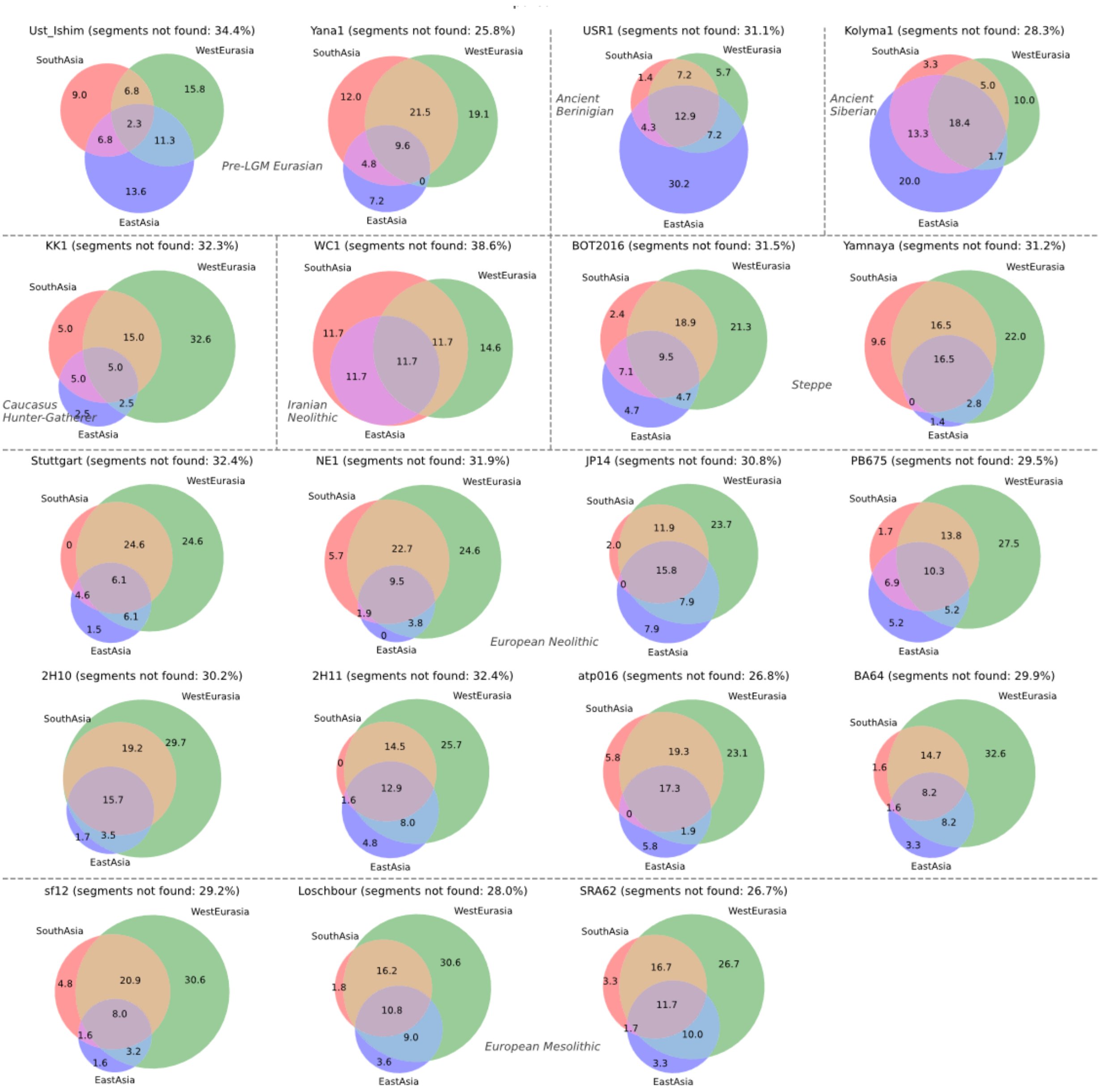


**Figure S15.** Proportion of the inferred archaic segments detected in the imputed 0.0625X genomes that are uniquely found in West Eurasia, South Asia, East Asia regions, using contemporary individuals in[22] introgression maps.

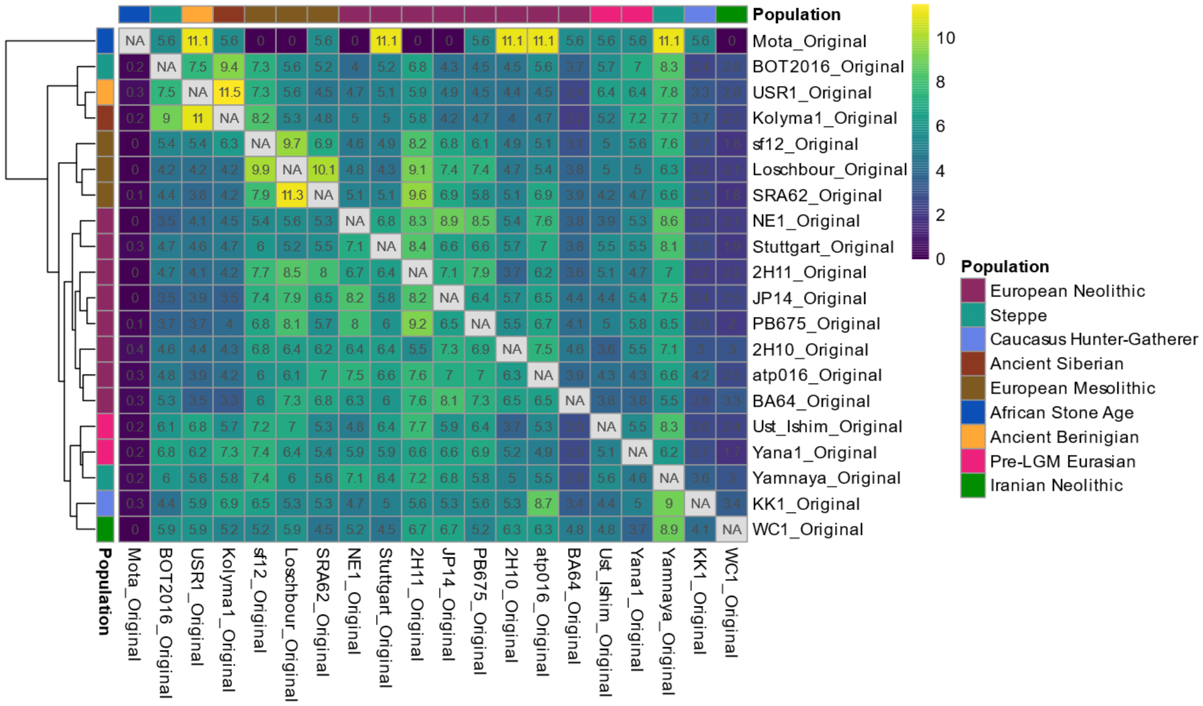


**Figure S16.** Heatmap of pairwise sharing of post-filtering introgressed genomic segments among ancient individuals for the original coverage dataset. Each cell shows the percentage of introgressed regions in the row individual that are also present in the column individual, based on collapsed shared segments. The matrix is normalized by row totals to account for differences in the amount of introgressed material per individual. Row clustering was performed using Spearman correlation and Ward's linkage, and the same order was applied to columns to preserve symmetry. Population affiliations are indicated by color-coded annotations.

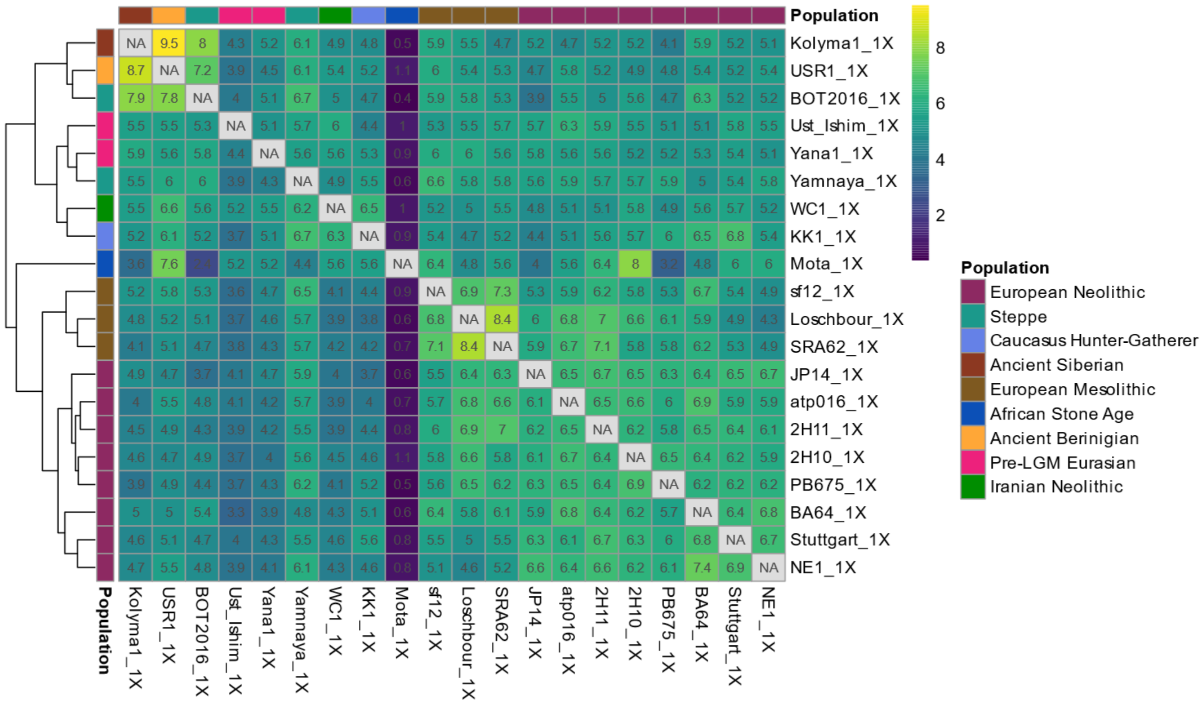


**Figure S17.** Heatmap of pairwise sharing of post-filtering introgressed genomic segments among ancient individuals for the imputed 1X dataset. Each cell shows the percentage of introgressed regions in the row individual that are also present in the column individual, based on collapsed shared segments. The matrix is normalized by row totals to account for differences in the amount of introgressed material per individual. Row clustering was performed using Spearman correlation and Ward's linkage, and the same order was applied to columns to preserve symmetry. Population affiliations are indicated by color-coded annotations.

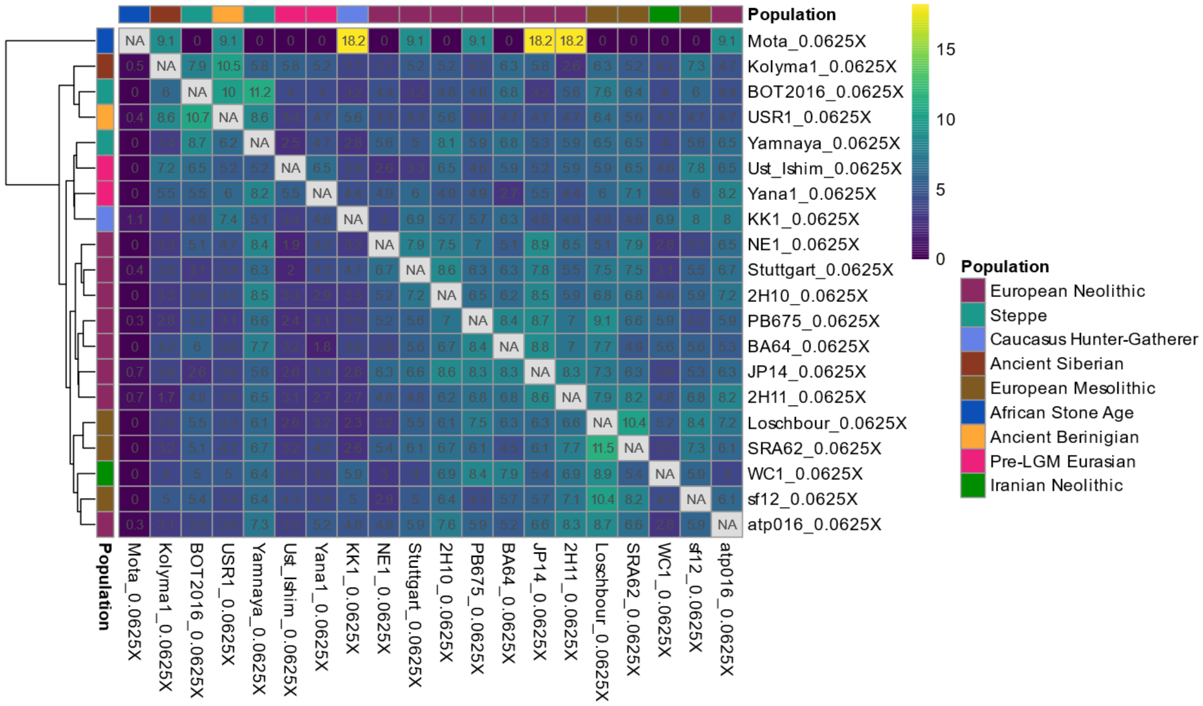


**Figure S18.** Heatmap of pairwise sharing of post-filtering introgressed genomic segments among ancient individuals for the imputed 0.0625X dataset. Each cell shows the percentage of introgressed regions in the row individual that are also present in the column individual, based on collapsed shared segments. The matrix is normalized by row totals to account for differences in the amount of introgressed material per individual. Row clustering was performed using Spearman correlation and Ward's linkage, and the same order was applied to columns to preserve symmetry. Population affiliations are indicated by color-coded annotations.

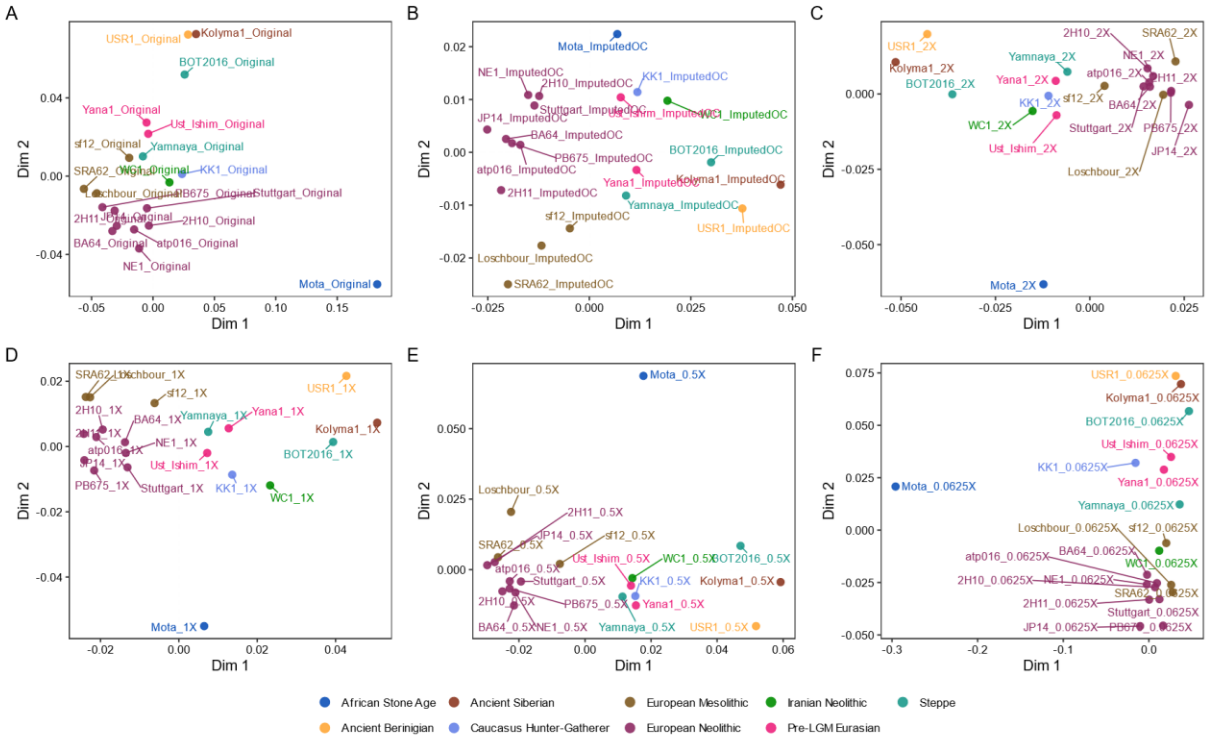


**Figure S19.** Multidimensional scaling (MDS) plots of shared introgressed segments across coverage levels, based on segment count overlap.

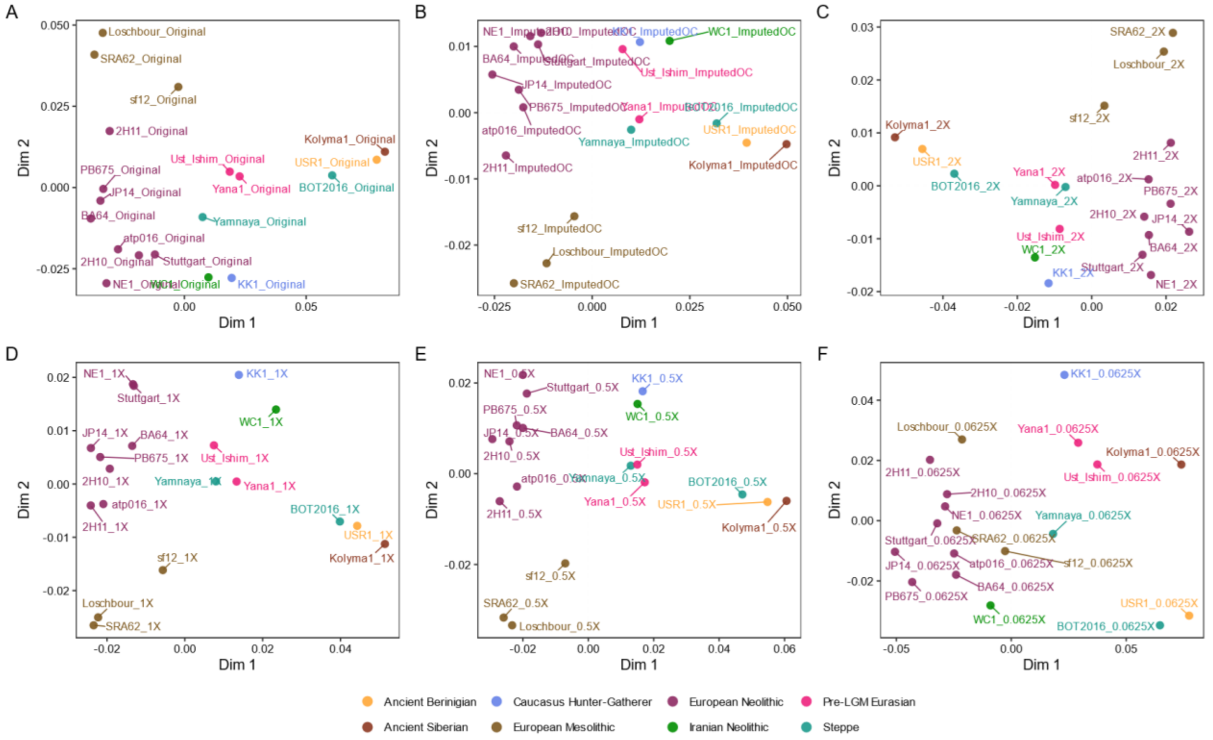


**Figure S20.** Multidimensional scaling (MDS) plots of shared introgressed segments across coverage levels, based on segment count overlap. In this analysis the individual Mota has been excluded.

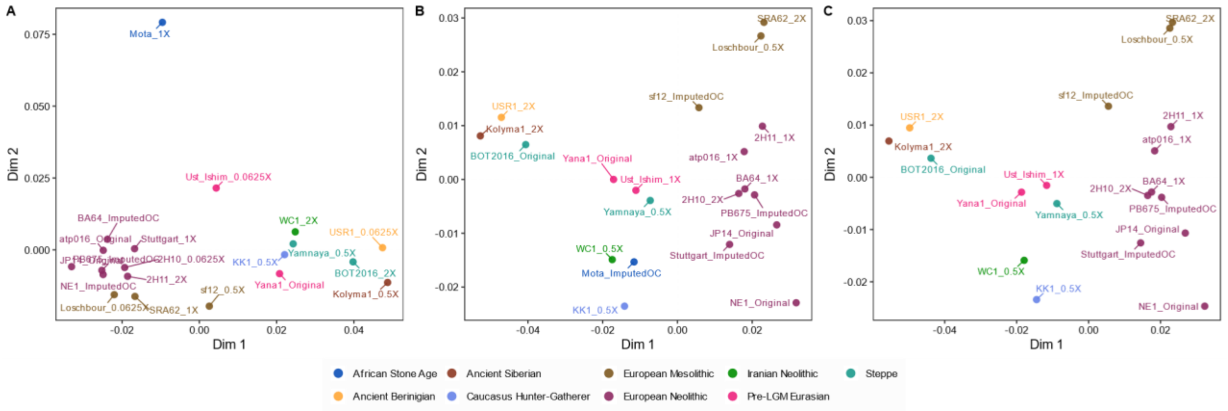


**Figure S21.** Multidimensional scaling (MDS) plots of shared introgressed segments across randomized individual-coverage combinations. (A) Includes all randomly assigned individuals and coverages. (B) Same procedure, excluding 0.0625X coverage samples. (C) Same procedure, excluding 0.0625X coverage samples and the Mota individual.

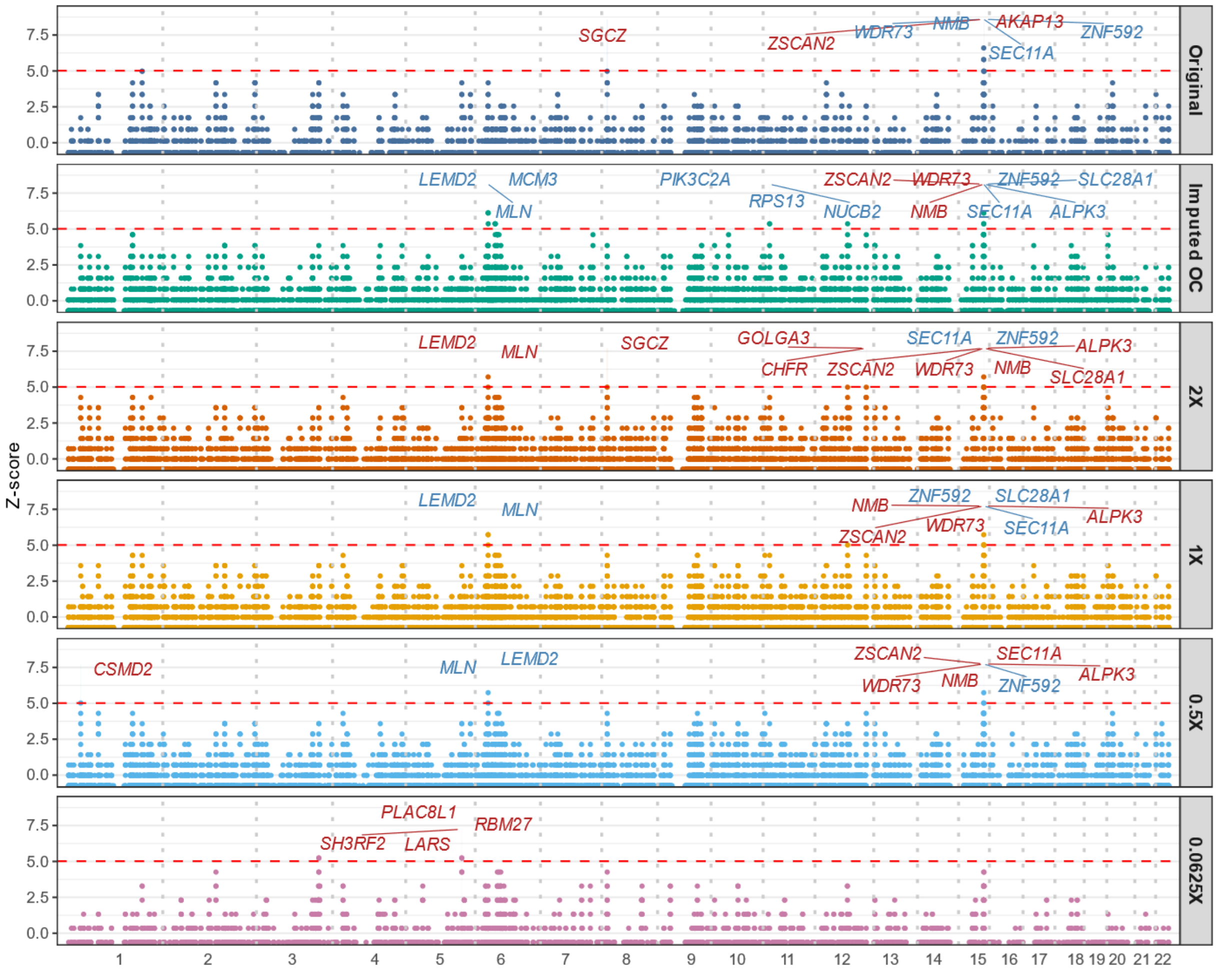


**Figure S22.** Manhattan plot of frequencies of archaic introgressed regions in ancient European individuals. The Y-axis shows Z-scores for the frequency of archaic segments that are positioned along the genome on the X-axis, with vertical dotted lines marking chromosome boundaries. The dashed red line indicates the Bonferroni-corrected significance threshold (α = 0.01, one-tailed test). Labels indicate genes detected in core regions (adjusted p < 0.01, red) or extended regions (adjusted p < 0.05, blue).

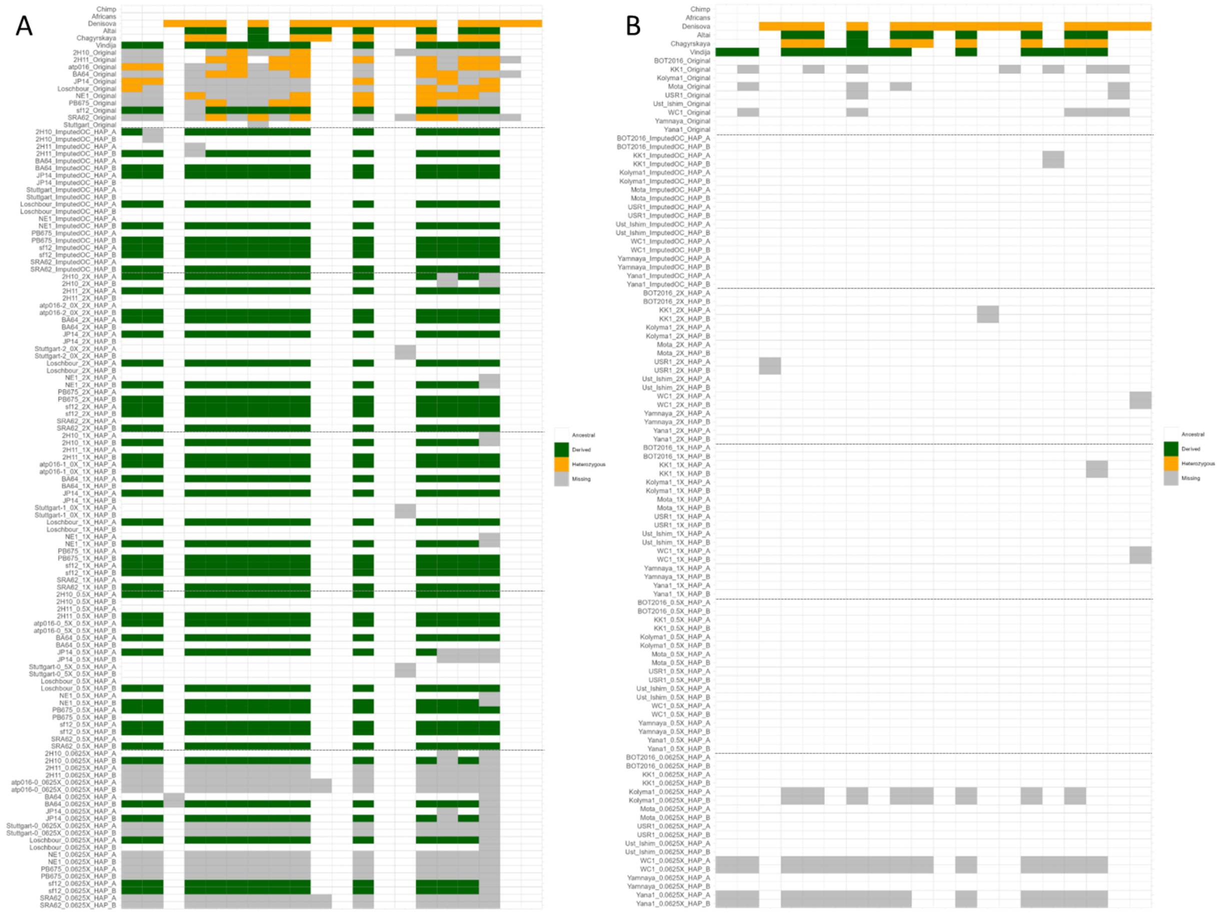


**Figure S23.** *Haplotype-level heatmap of archaic-derived SNPs from the beginning of LEMD2 to the end of MLN genes region across chimpanzee, African, archaic, and non-imputed and imputed ancient human genomes after filtering for GP > 99%, A) for European individuals and B) for not european individuals. Each row represents a haplotype or diploid individual, and each column corresponds to a possible derived allele. Alleles are classified as Ancestral (white), Derived (green), Heterozygous (orange) and Missing (grey). Imputed genomes are split into phased haplotypes (HAP_A and HAP_B), while archaic and original genomes are unphased.*

*Supplementary Tables*

**Table S1**. Metadata for the ancient individuals analyzed in this study.

**Table S2.** Metadata for the reference and modern individuals used in this study.

**Table S3**. Data used to make Venn Diagrams in figure 4A., including all coverages.

**Table S4**. Data used to make the Heatmap in figure S10-12, including all coverages.

**Table S5.** Summary of sample counts per coverage level with overlapping introgressed fragments in 14 known candidate genes.

**Table S6.** Per-sample overlap of archaic introgressed fragments with 14 known candidate genes.

**Table S7.** List of SNPs used to investigate archaic introgression at the *BNC2* locus. Each site is biallelic and annotated with the reference (REF), alternative (ALT), and inferred ancestral and derived alleles. Only SNPs where archaic genomes carry the derived allele and African genomes are homozygous for the ancestral allele were retained.

**Table S8.** Presence of significant genes across coverages in core and extended introgressed regions. The table lists all protein-coding genes overlapping significant archaic-introgressed segments, categorized into core and extended regions. Core regions are defined as segments where at least one subregion has a Bonferroni-adjusted p-value $< 0.01$. Extended regions are composed of adjacent segments that do not meet the core significance threshold individually (p_adj_bonferroni $\geq 0.01$), but are contiguous with core segments and show elevated frequencies (p_adj $< 0.05$), suggesting potential broader introgression.

**Table S9.** Segment-level annotation of introgressed genes across core and extended regions. This table lists all protein-coding genes overlapping archaic introgressed segments across all coverage levels.

**Table S10**. List of SNPs used to investigate archaic introgression from the start of *LEDM2* gene to the end of *MLN* gene. Each site is biallelic and annotated with the reference (REF), alternative (ALT), and inferred ancestral and derived alleles. Only SNPs where archaic genomes carry the derived allele and African genomes are homozygous for the ancestral allele were retained. In the last column is reported if the position is in the Strict Mask region

14. Günther, T. *et al.* Population genomics of Mesolithic Scandinavia: Investigating early

postglacial migration routes and high-latitude adaptation. *PLoS Biol* **16**, e2003703 (2018).
15. Jones, E. R. *et al.* Upper Palaeolithic genomes reveal deep roots of modern Eurasians. *Nat Commun* **6**, 8912 (2015).
16. Lazaridis, I. *et al.* Ancient human genomes suggest three ancestral populations for present-day Europeans. *Nature* **513**, 409–413 (2014).
17. Moreno-Mayar, J. V. *et al.* Terminal Pleistocene Alaskan genome reveals first founding population of Native Americans. *Nature* **553**, 203–207 (2018).
18. Sikora, M. *et al.* The population history of northeastern Siberia since the Pleistocene. *Nature* **570**, 182–188 (2019).
19. Rubinacci, S., Ribeiro, D. M., Hofmeister, R. J. & Delaneau, O. Efficient phasing and imputation of low-coverage sequencing data using large reference panels. *Nat. Genet.* **53**, 120–126 (2021).
20. Patterson, N. *et al.* Ancient admixture in human history. *Genetics* **192**, 1065–1093 (2012).
21. Moorjani, P. *et al.* The history of African gene flow into Southern Europeans, Levantines, and Jews. *PLoS Genet* **7**, e1001373 (2011).
22. Skov, L. *et al.* Detecting archaic introgression using an unadmixed outgroup. *PLoS Genet.* **14**, e1007641 (2018).
23. Iasi, L. N. M. *et al.* Neanderthal ancestry through time: Insights from genomes of ancient and present-day humans. *Science* **386**, eadq3010 (2024).
24. Yuan, K. *et al.* Refining models of archaic admixture in Eurasia with ArchaicSeeker 2.0. *Nat Commun* **12**, 6232 (2021).
25. Ongaro, L. & Huerta-Sanchez, E. A history of multiple Denisovan introgression events in modern humans. *Nat Genet* **56**, 2612–2622 (2024).
26. Reilly, P. F., Tjahjadi, A., Miller, S. L., Akey, J. M. & Tucci, S. The contribution of Neanderthal introgression to modern human traits. *Curr. Biol.* **32**, R970–R983 (2022).
27. Witt, K. E., Funk, A., Añorve-Garibay, V., Fang, L. L. & Huerta-Sánchez, E. The Impact of Modern Admixture on Archaic Human Ancestry in Human Populations. *Genome Biol. Evol.* **15**, (2023).
28. Gouy, A. & Excoffier, L. Polygenic Patterns of Adaptive Introgression in Modern Humans Are Mainly Shaped by Response to Pathogens. *Mol Biol Evol* **37**, 1420–1433 (2020).
29. Racimo, F., Marnetto, D. & Huerta-Sánchez, E. Signatures of Archaic Adaptive

Introgression in Present-Day Human Populations. *Mol Biol Evol* **34**, 296–317 (2017).
30. Li, H. & Durbin, R. Fast and accurate short read alignment with Burrows-Wheeler transform. *Bioinformatics* **25**, 1754–1760 (2009).
31. Danecek, P. *et al.* Twelve years of SAMtools and BCFtools. *Gigascience* **10**, (2021).
32. 1000 Genomes Project Consortium *et al.* A global reference for human genetic variation. *Nature* **526**, 68–74 (2015).
33. Korneliussen, T. S., Albrechtsen, A. & Nielsen, R. ANGSD: Analysis of Next Generation Sequencing Data. *BMC Bioinformatics* **15**, 356 (2014).
34. Prüfer, K. *et al.* The complete genome sequence of a Neanderthal from the Altai Mountains. *Nature* **505**, 43–49 (2014).
35. Prüfer, K. *et al.* A high-coverage Neandertal genome from Vindija Cave in Croatia. *Science* **358**, 655–658 (2017).
36. Mafessoni, F. *et al.* A high-coverage Neandertal genome from Chagyrskaya Cave. *Proc. Natl. Acad. Sci. U. S. A.* **117**, 15132–15136 (2020).
37. Meyer, M. *et al.* A high-coverage genome sequence from an archaic Denisovan individual. *Science* **338**, 222–226 (2012).
38. Huerta-Sánchez, E. *et al.* Altitude adaptation in Tibetans caused by introgression of Denisovan-like DNA. *Nature* **512**, 194–197 (2014).
39. Malaspinas, A. S. *et al.* A genomic history of Aboriginal Australia. *Nature* **538**, (2016).
40. Vernot, B. *et al.* Excavating Neandertal and Denisovan DNA from the genomes of Melanesian individuals. *Science* **352**, 235–239 (2016).
41. Mallick, S. *et al.* The Simons Genome Diversity Project: 300 genomes from 142 diverse populations. *Nature* **538**, 201–206 (2016).
