## SupplementaryFigures for "Archaic ancestry inference in imputed ancient human genomes"

FigureS1

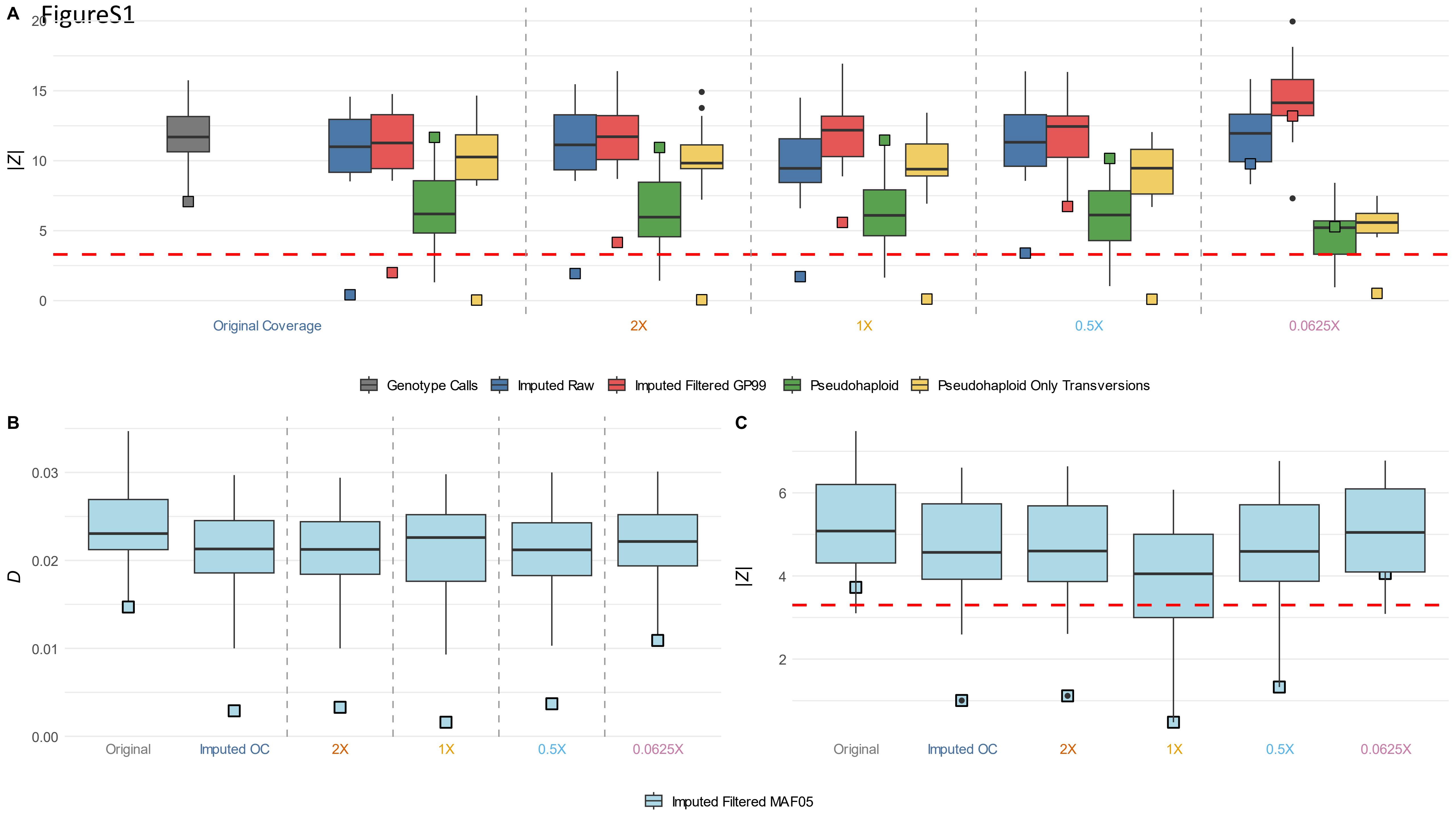

A
|Z|
20
15
10
5
0
Original Coverage
2X
1X
0.5X
0.0625X
Genotype Calls
Imputed Raw
Imputed Filtered GP99
Pseudohaploid
Pseudohaploid Only Transversions
B
D
0.03
0.02
0.01
0.00
Original
Imputed OC
2X
1X
0.5X
0.0625X
C
|Z|
6
4
2
Original
Imputed OC
2X
1X
0.5X
0.0625X
Imputed Filtered MAF05

FigureS2

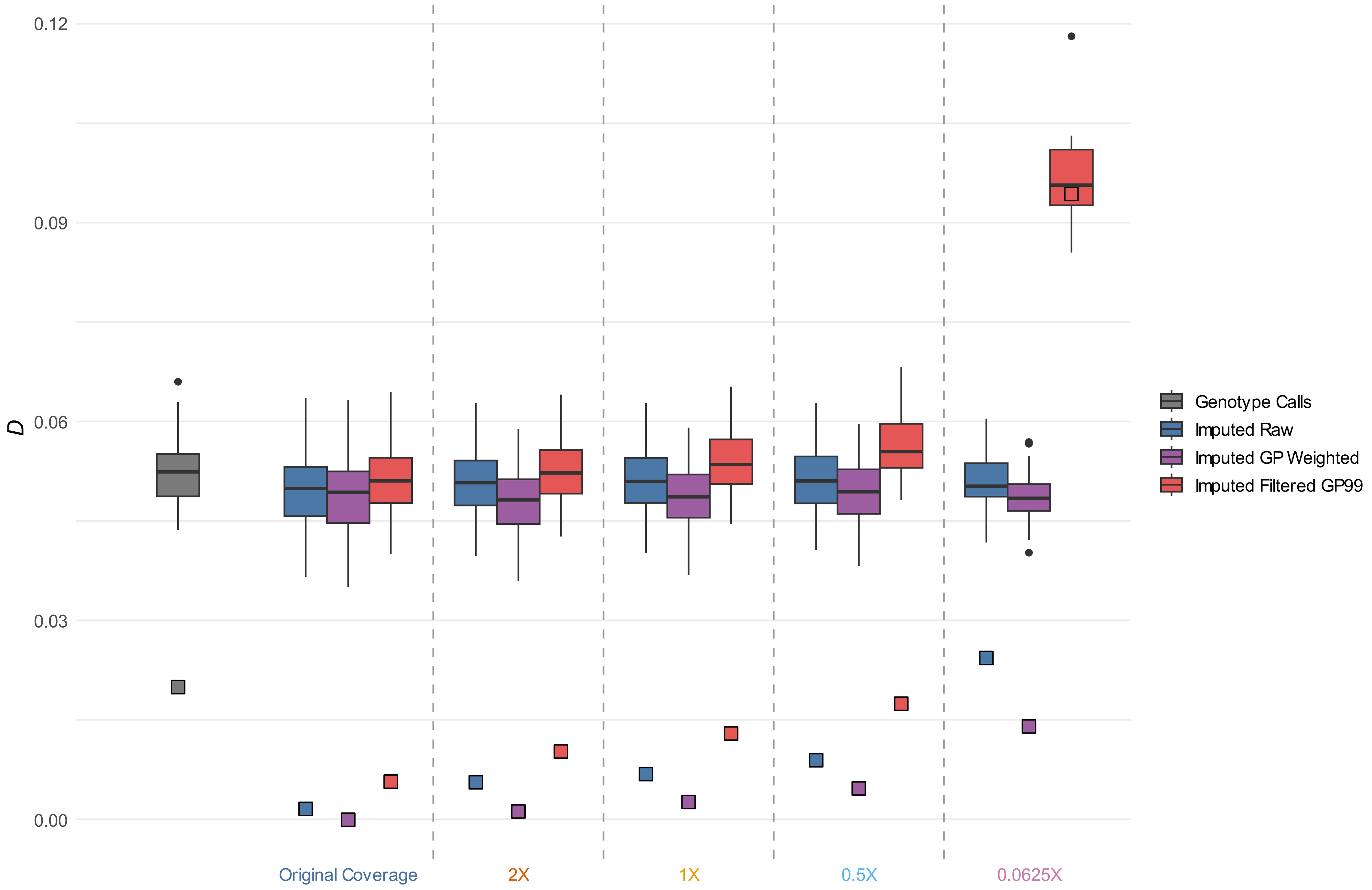
0.12
0.09
0.06
0.03
0.00
D
Original Coverage
2X
1X
0.5X
0.0625X
Genotype Calls
Imputed Raw
Imputed GP Weighted
Imputed Filtered GP99

FigureS3

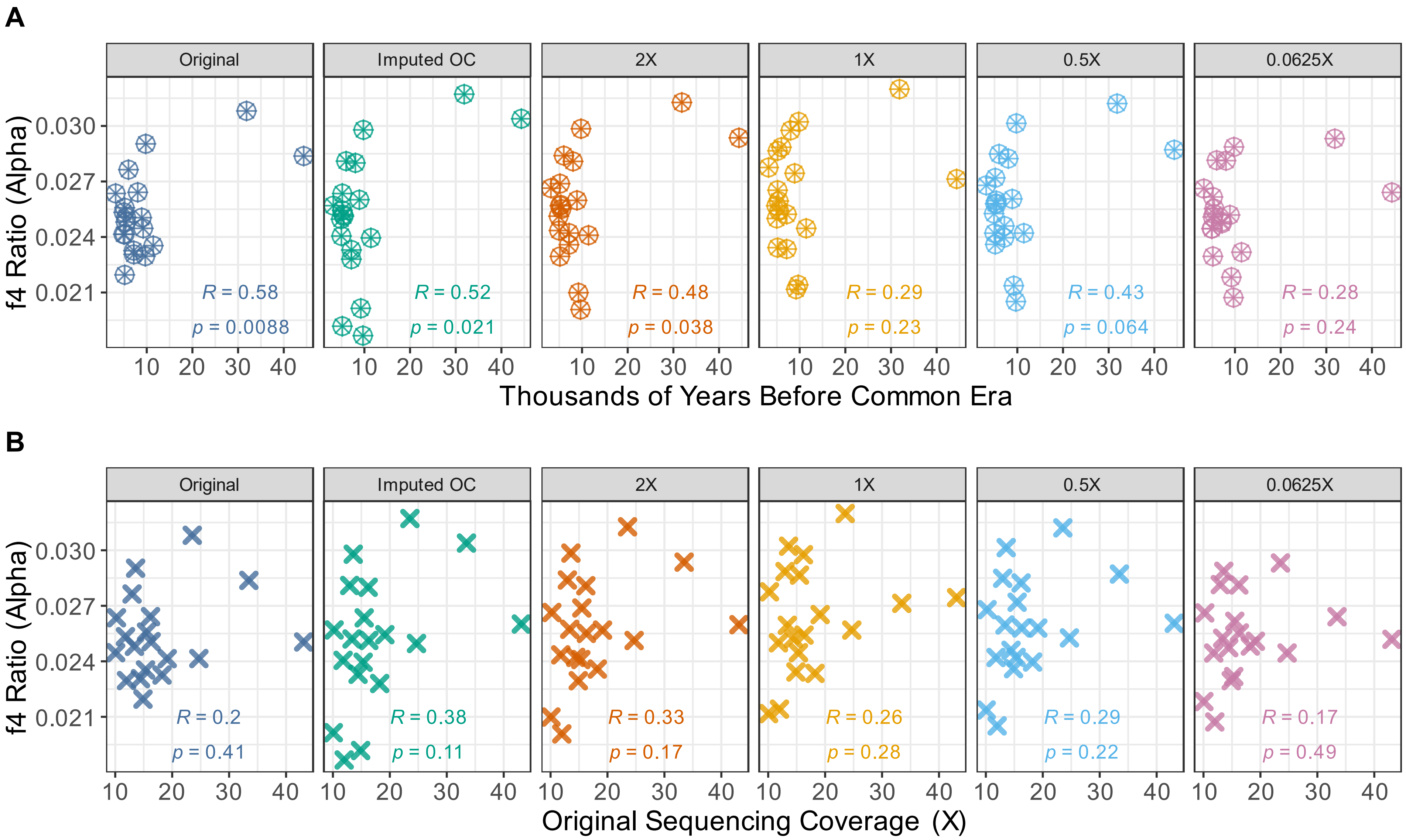
A
Original
Imputed OC
2X
1X
0.5X
0.0625X
f4 Ratio (Alpha)
0.030
0.027
0.024
0.021
R = 0.58
p = 0.0088
R = 0.52
p = 0.021
R = 0.48
p = 0.038
R = 0.29
p = 0.23
R = 0.43
p = 0.064
R = 0.28
p = 0.24
10 20 30 40
Thousands of Years Before Common Era
B
Original
Imputed OC
2X
1X
0.5X
0.0625X
f4 Ratio (Alpha)
0.030
0.027
0.024
0.021
R = 0.2
p = 0.41
R = 0.38
p = 0.11
R = 0.33
p = 0.17
R = 0.26
p = 0.28
R = 0.29
p = 0.22
R = 0.17
p = 0.49
10 20 30 40
Original Sequencing Coverage (X)

FigureS4

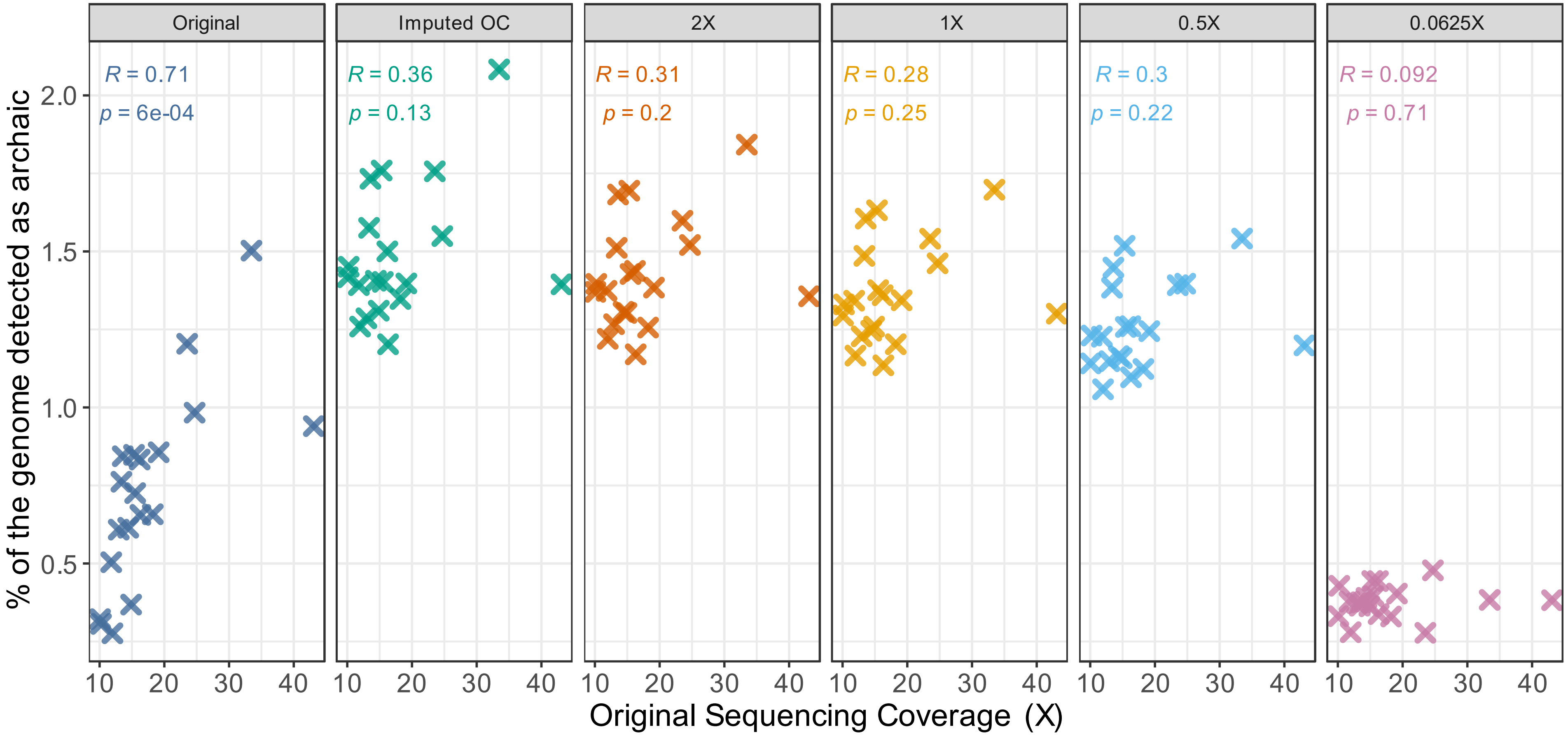

Original
Imputed OC
2X
1X
0.5X
0.0625X
R = 0.71
p = 6e-04
R = 0.36
p = 0.13
R = 0.31
p = 0.2
R = 0.28
p = 0.25
R = 0.3
p = 0.22
R = 0.092
p = 0.71
% of the genome detected as archaic
2.0
1.5
1.0
0.5
10 20 30 40
Original Sequencing Coverage (X)

Figure S5

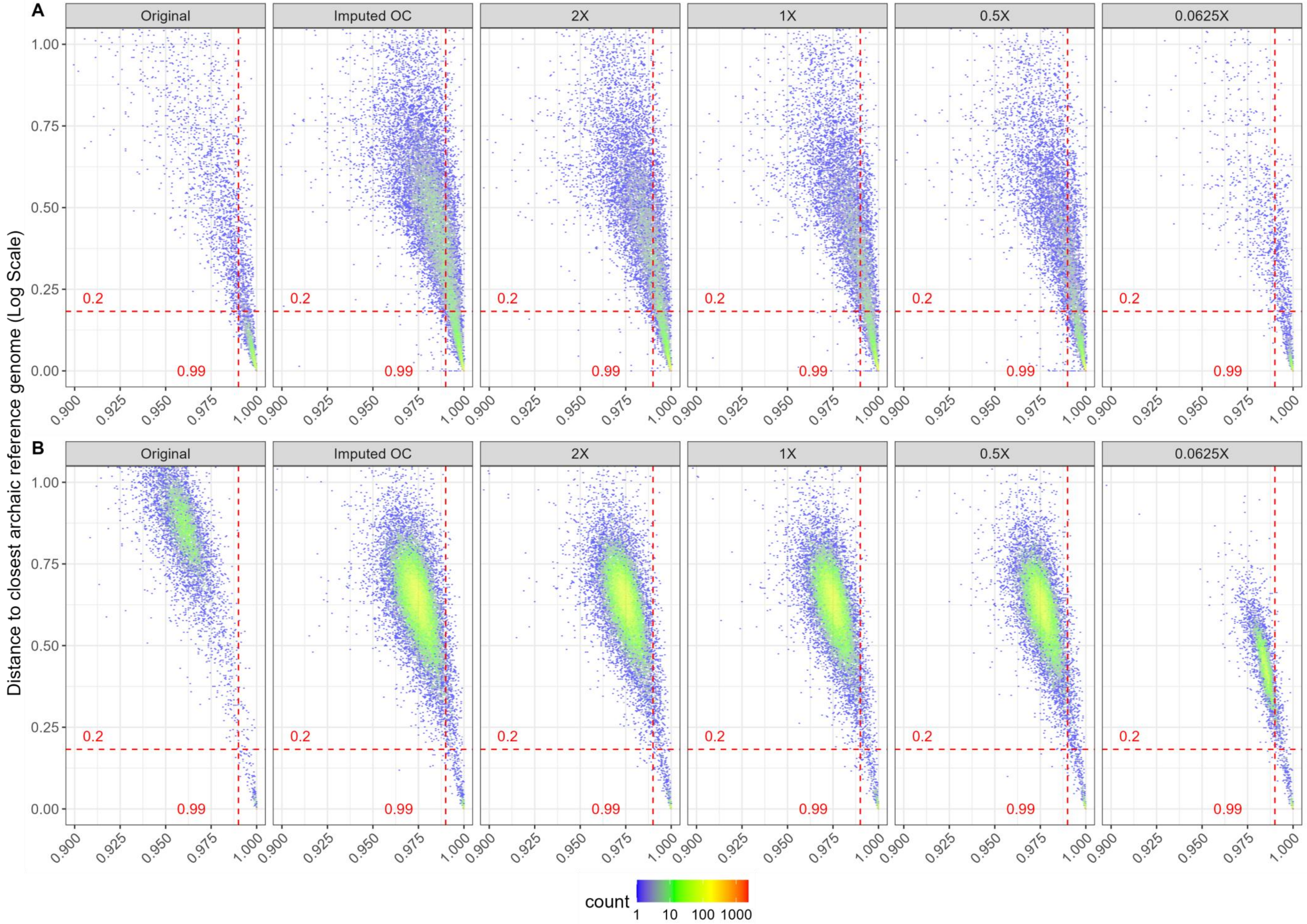

A
B
Original
Imputed OC
2X
1X
0.5X
0.0625X
Distance to closest archaic reference genome (Log Scale)
0.2
0.99
count
1 10 100 1000

### Figure S6

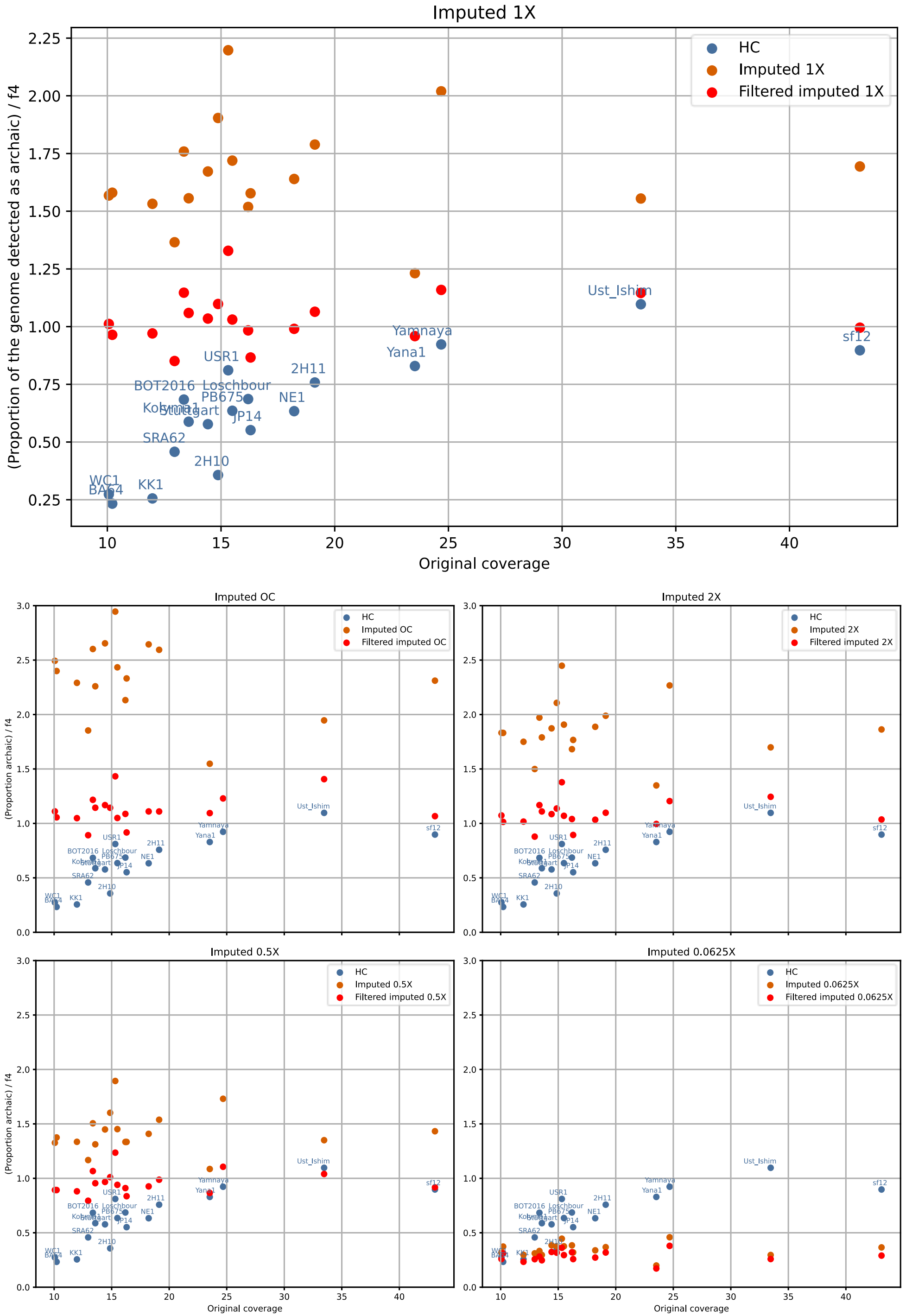

Imputed 1X
HC
Imputed 1X
Filtered imputed 1X
(Proportion of the genome detected as archaic) / f4
Original coverage
2.25
2.00
1.75
1.50
1.25
1.00
0.75
0.50
0.25
10
15
20
25
30
35
40
Ust_Ishim
Yamnaya
Yana1
sf12
USR1
2H11
BOT2016
Loschbour
PB675
NE1
Kolyma1
Stuttgart
JP14
SRA62
2H10
WC1
BA64
KK1
Imputed OC
Imputed OC
Filtered imputed OC
Imputed 2X
Imputed 2X
Filtered imputed 2X
Imputed 0.5X
Imputed 0.5X
Filtered imputed 0.5X
Imputed 0.0625X
Imputed 0.0625X
Filtered imputed 0.0625X
(Proportion archaic) / f4
Original coverage

Figure S7

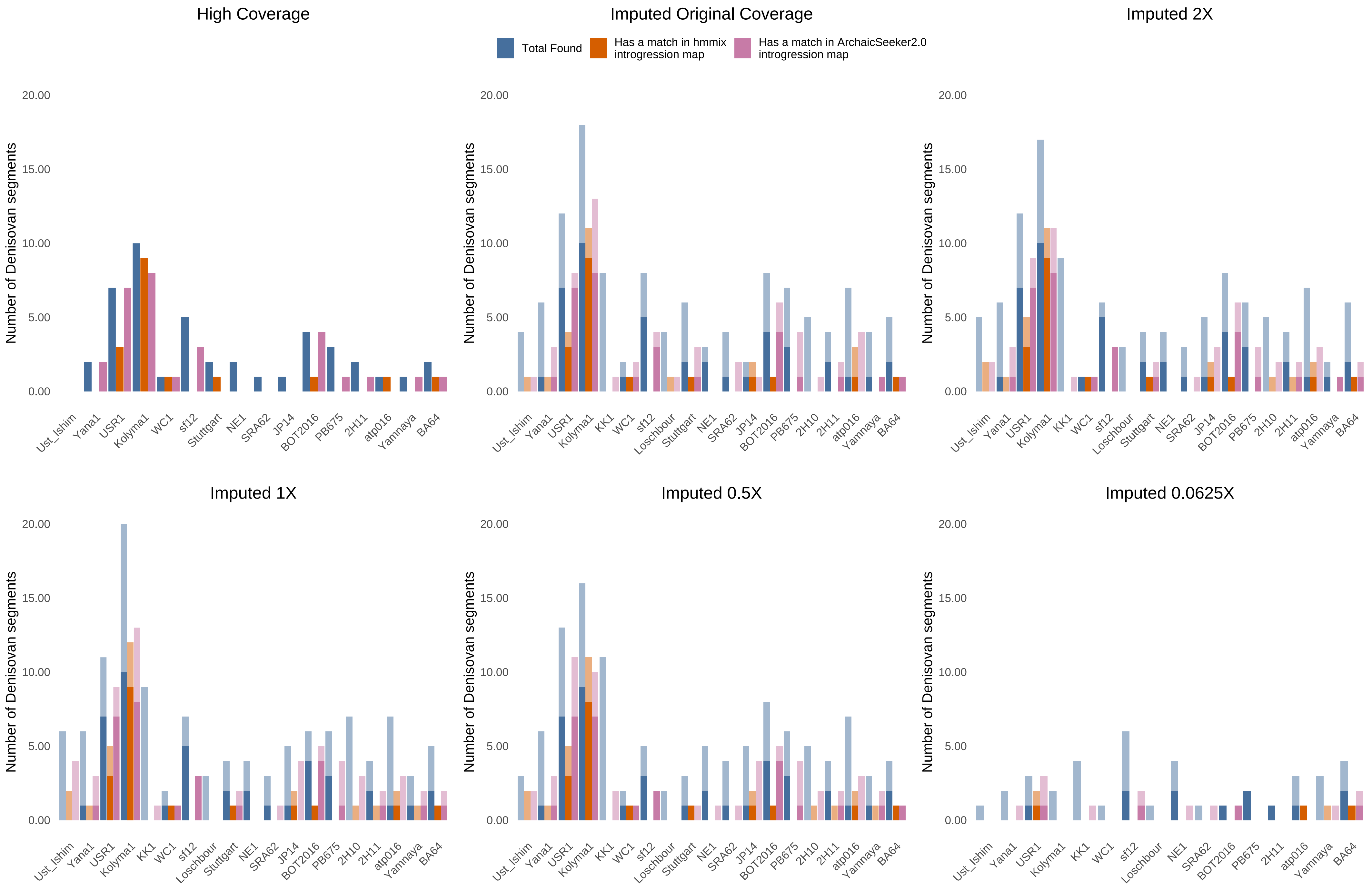

High Coverage
Imputed Original Coverage
Imputed 2X
Imputed 1X
Imputed 0.5X
Imputed 0.0625X
Total Found
Has a match in hmmix introgression map
Has a match in ArchaicSeeker2.0 introgression map
Number of Denisovan segments
0.00
5.00
10.00
15.00
20.00
Ust_Ishim
Yana1
USR1
Kolyma1
KK1
WC1
sf12
Loschbour
Stuttgart
NE1
SRA62
JP14
BOT2016
PB675
2H10
2H11
atp016
Yamnaya
BA64

Figure S8

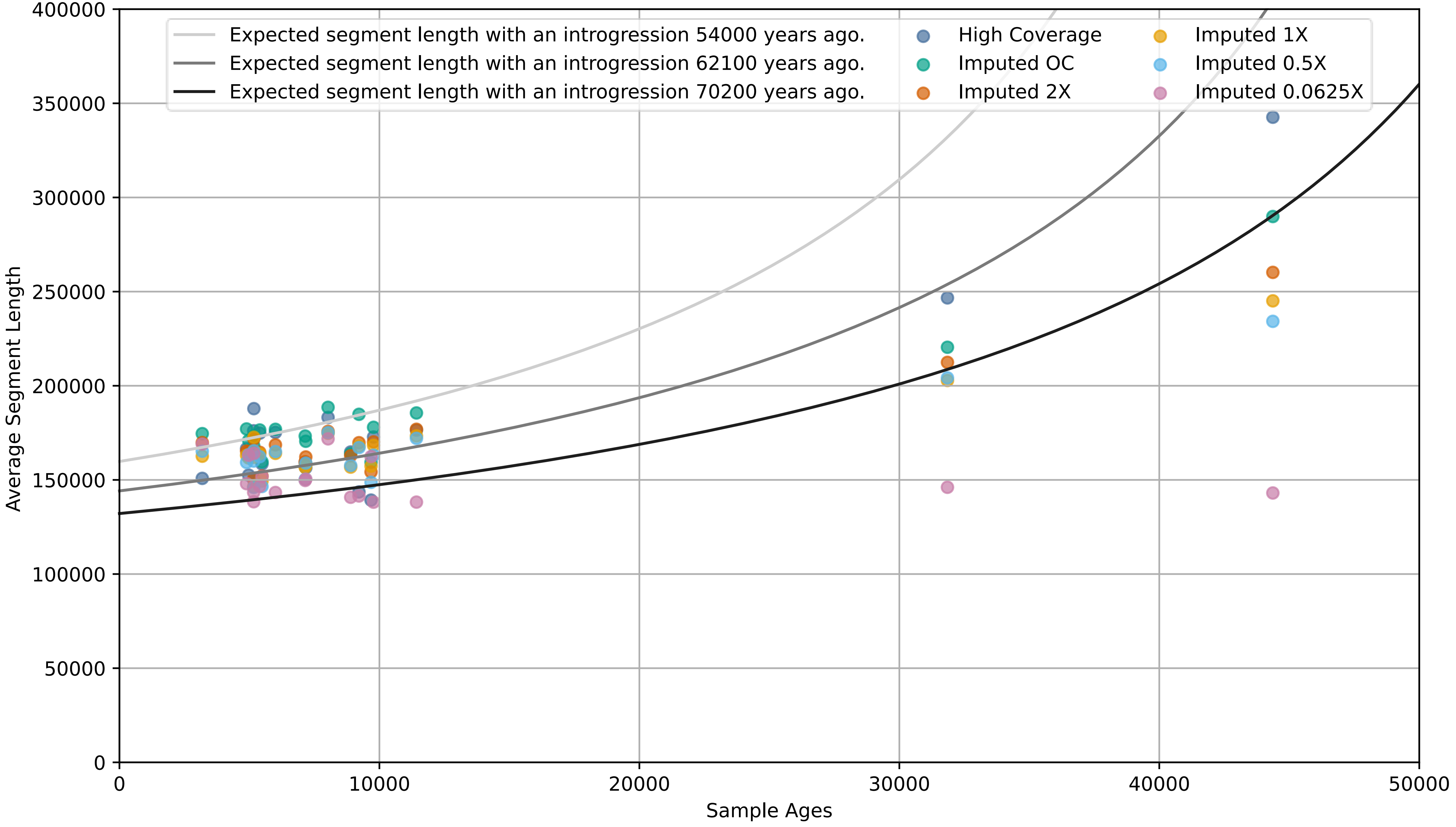
Expected segment length with an introgression 54000 years ago.
Expected segment length with an introgression 62100 years ago.
Expected segment length with an introgression 70200 years ago.
High Coverage
Imputed OC
Imputed 2X
Imputed 1X
Imputed 0.5X
Imputed 0.0625X
Average Segment Length
Sample Ages
0
50000
100000
150000
200000
250000
300000
350000
400000
0
10000
20000
30000
40000
50000

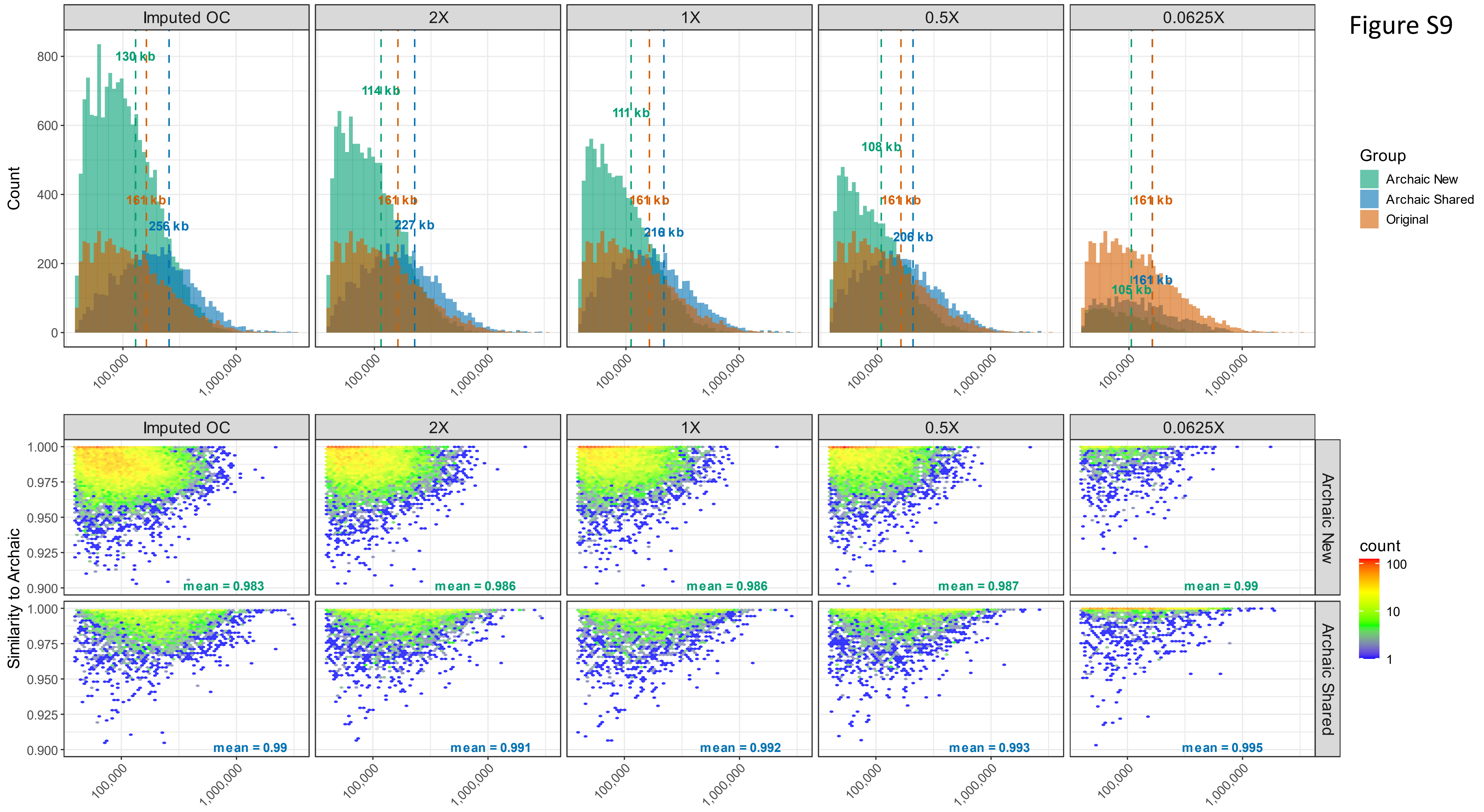
Figure S9
Imputed OC
2X
1X
0.5X
0.0625X
Count
130 kb
161 kb
256 kb
114 kb
227 kb
111 kb
216 kb
108 kb
206 kb
105 kb
Group
Archaic New
Archaic Shared
Original
Similarity to Archaic
mean = 0.983
mean = 0.986
mean = 0.986
mean = 0.987
mean = 0.99
mean = 0.99
mean = 0.991
mean = 0.992
mean = 0.993
mean = 0.995
Archaic New
Archaic Shared
count
100
10
1
100,000
1,000,000

Figure S10

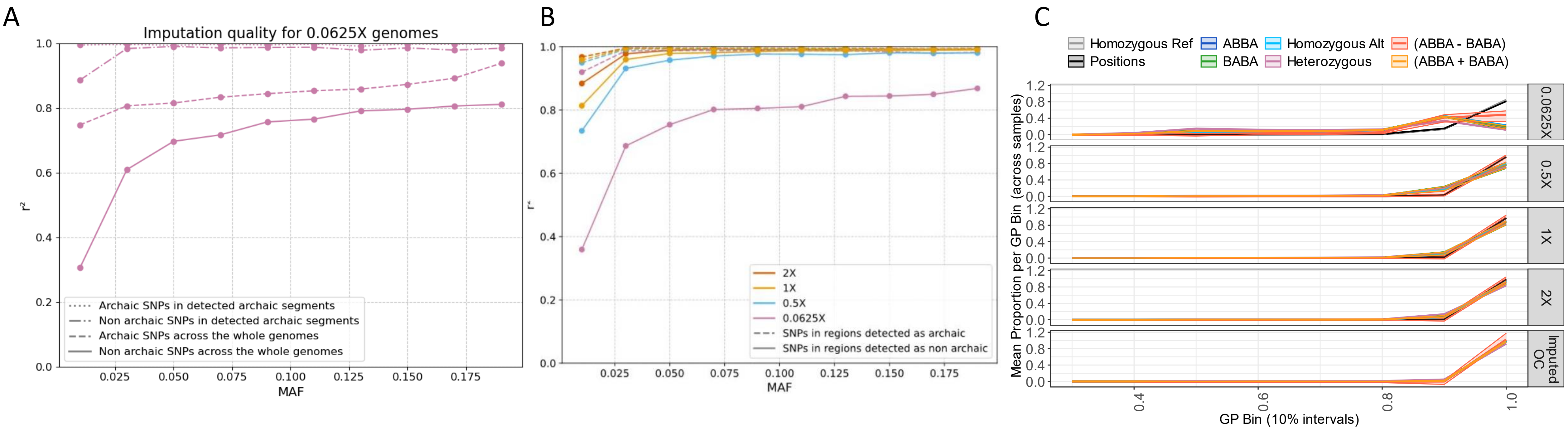
A
Imputation quality for 0.0625X genomes
r²
MAF
Archaic SNPs in detected archaic segments
Non archaic SNPs in detected archaic segments
Archaic SNPs across the whole genomes
Non archaic SNPs across the whole genomes
B
r²
MAF
2X
1X
0.5X
0.0625X
SNPs in regions detected as archaic
SNPs in regions detected as non archaic
C
Homozygous Ref
Positions
ABBA
BABA
Homozygous Alt
Heterozygous
(ABBA - BABA)
(ABBA + BABA)
Mean Proportion per GP Bin (across samples)
GP Bin (10% intervals)
0.0625X
0.5X
1X
2X
Imputed OC

Figure S11

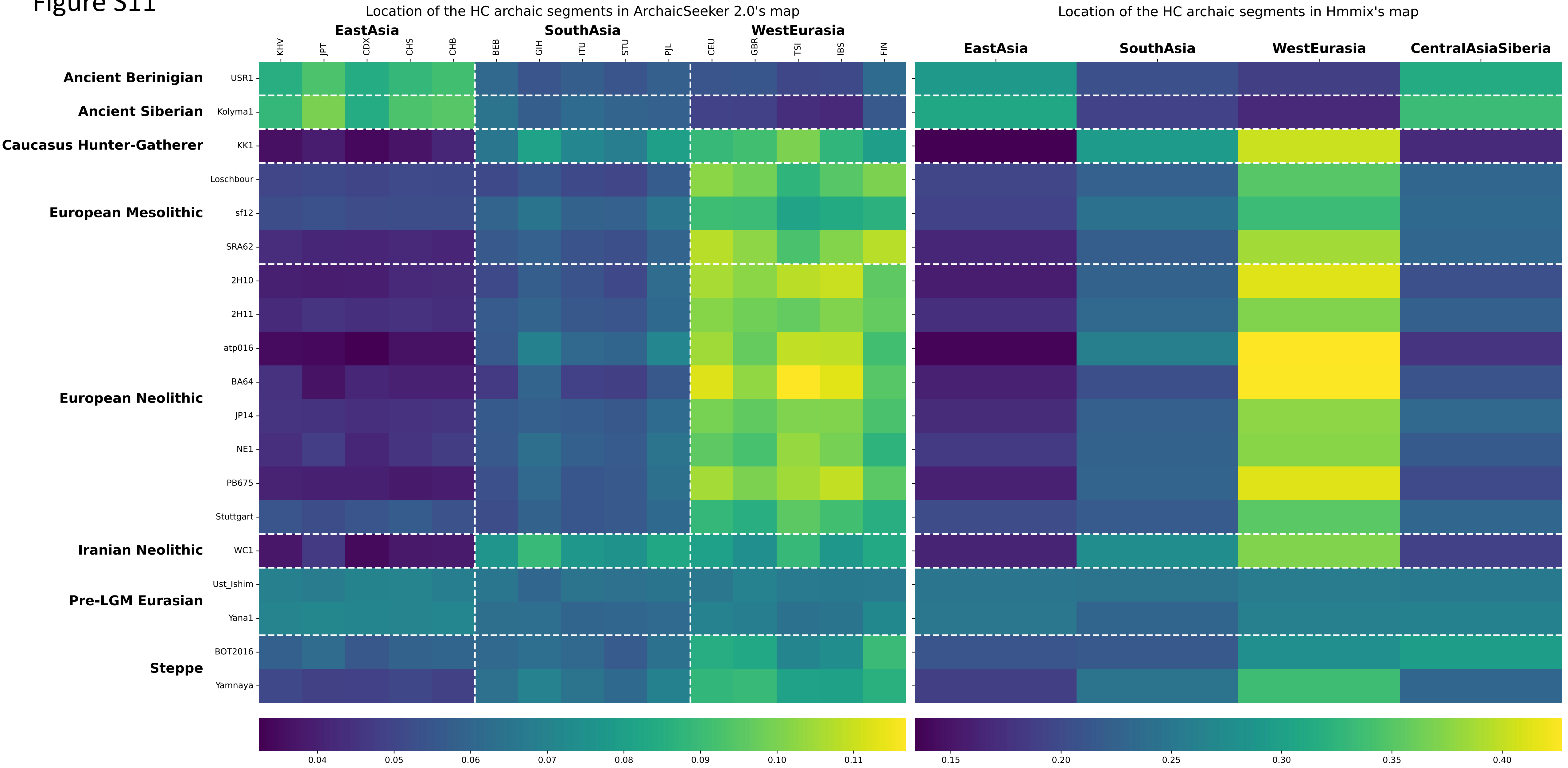
Location of the HC archaic segments in ArchaicSeeker 2.0's map
Location of the HC archaic segments in Hmmix's map
EastAsia
SouthAsia
WestEurasia
CentralAsiaSiberia
KHV
JPT
CDX
CHS
CHB
BEB
GIH
ITU
STU
PJL
CEU
GBR
TSI
IBS
FIN
Ancient Berinigian
Ancient Siberian
Caucasus Hunter-Gatherer
European Mesolithic
European Neolithic
Iranian Neolithic
Pre-LGM Eurasian
Steppe
USR1
Kolyma1
KK1
Loschbour
sf12
SRA62
2H10
2H11
atp016
BA64
JP14
NE1
PB675
Stuttgart
WC1
Ust_Ishim
Yana1
BOT2016
Yamnaya
0.04
0.05
0.06
0.07
0.08
0.09
0.10
0.11
0.15
0.20
0.25
0.30
0.35
0.40

Figure S12

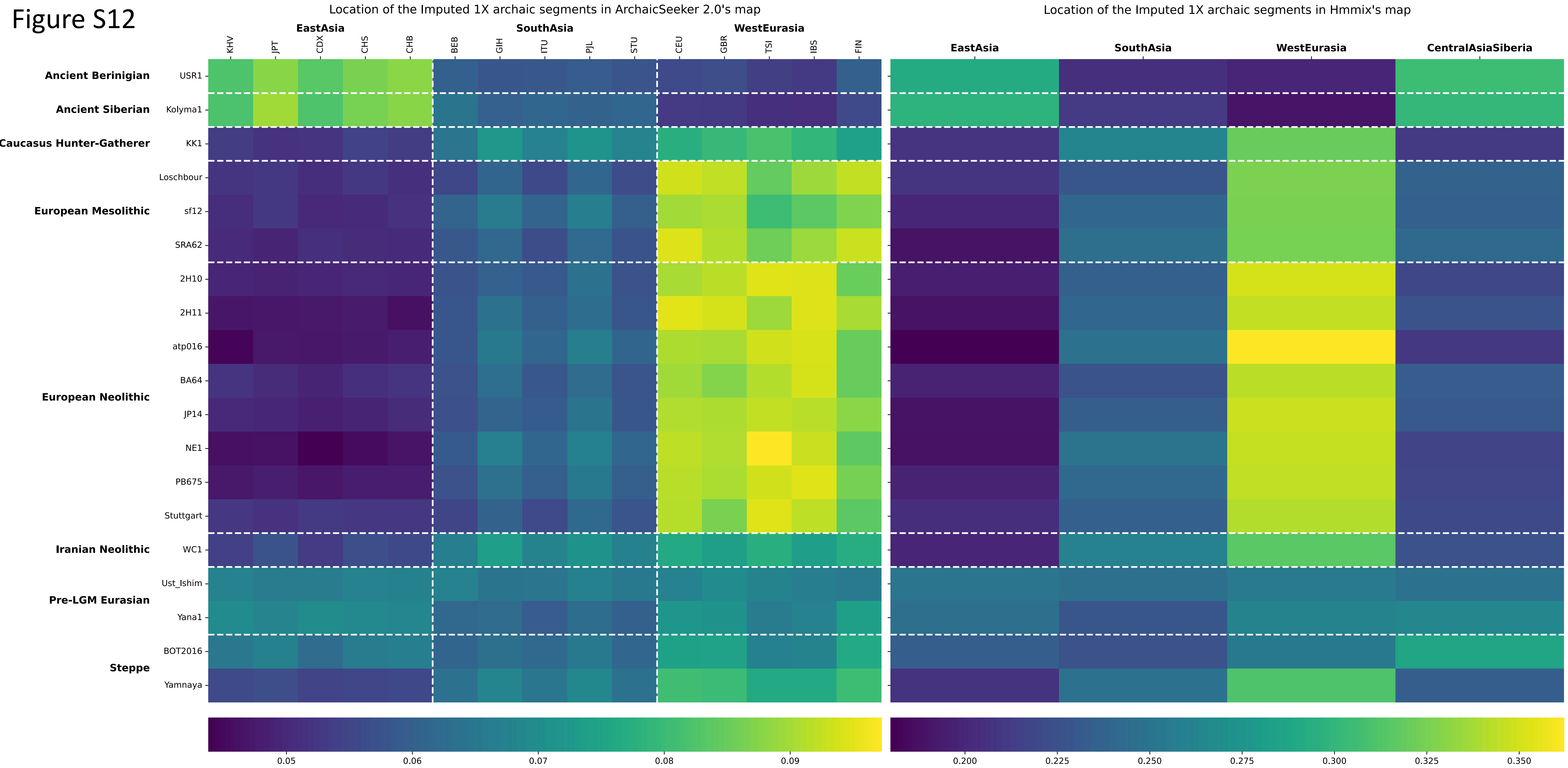

Location of the Imputed 1X archaic segments in ArchaicSeeker 2.0's map
Location of the Imputed 1X archaic segments in Hmmix's map
EastAsia
SouthAsia
WestEurasia
CentralAsiaSiberia
KHV
JPT
CDX
CHS
CHB
BEB
GIH
ITU
PJL
STU
CEU
GBR
TSI
IBS
FIN
Ancient Beringian
Ancient Siberian
Caucasus Hunter-Gatherer
European Mesolithic
European Neolithic
Iranian Neolithic
Pre-LGM Eurasian
Steppe
USR1
Kolyma1
KK1
Loschbour
sf12
SRA62
2H10
2H11
atp016
BA64
JP14
NE1
PB675
Stuttgart
WC1
Ust_Ishim
Yana1
BOT2016
Yamnaya
0.05
0.06
0.07
0.08
0.09
0.200
0.225
0.250
0.275
0.300
0.325
0.350

### Figure S13

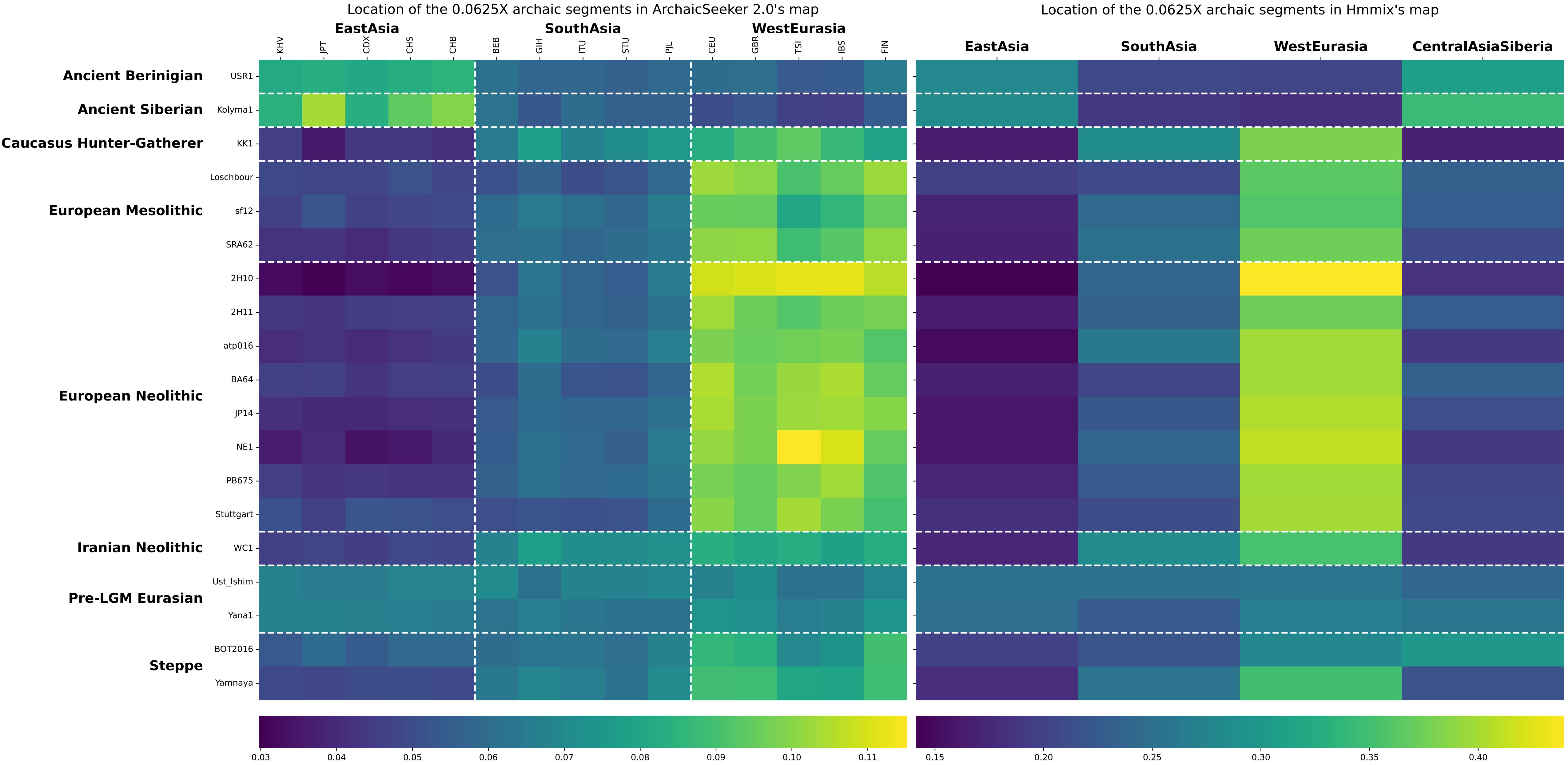

Location of the 0.0625X archaic segments in ArchaicSeeker 2.0's map
Location of the 0.0625X archaic segments in Hmmix's map
EastAsia
SouthAsia
WestEurasia
CentralAsiaSiberia
KHV
JPT
CDX
CHS
CHB
BEB
GIH
ITU
STU
PJL
CEU
GBR
TSI
IBS
FIN
Ancient Berinigian
Ancient Siberian
Caucasus Hunter-Gatherer
European Mesolithic
European Neolithic
Iranian Neolithic
Pre-LGM Eurasian
Steppe
USR1
Kolyma1
KK1
Loschbour
sf12
SRA62
2H10
2H11
atp016
BA64
JP14
NE1
PB675
Stuttgart
WC1
Ust_Ishim
Yana1
BOT2016
Yamnaya
0.03
0.04
0.05
0.06
0.07
0.08
0.09
0.10
0.11
0.15
0.20
0.25
0.30
0.35
0.40

### Figure S14

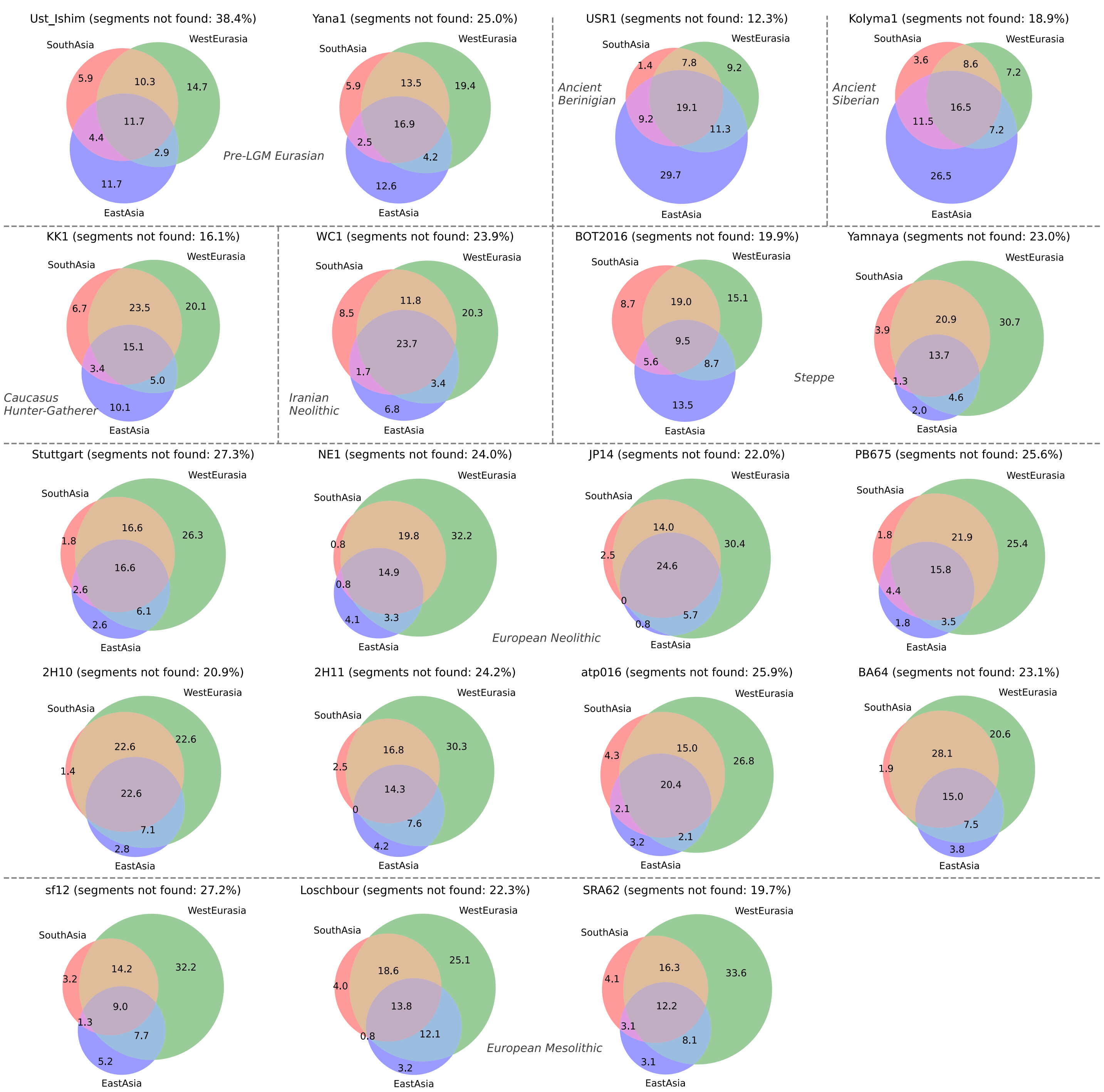

Ust_Ishim (segments not found: 38.4%)
SouthAsia
WestEurasia
EastAsia
5.9
10.3
14.7
11.7
4.4
2.9
11.7
Yana1 (segments not found: 25.0%)
SouthAsia
WestEurasia
EastAsia
5.9
13.5
19.4
16.9
2.5
4.2
12.6
Pre-LGM Eurasian
USR1 (segments not found: 12.3%)
SouthAsia
WestEurasia
EastAsia
1.4
7.8
9.2
19.1
9.2
11.3
29.7
Ancient Berinigian
Kolyma1 (segments not found: 18.9%)
SouthAsia
WestEurasia
EastAsia
3.6
8.6
7.2
16.5
11.5
7.2
26.5
Ancient Siberian
KK1 (segments not found: 16.1%)
SouthAsia
WestEurasia
EastAsia
6.7
23.5
20.1
15.1
3.4
5.0
10.1
Caucasus Hunter-Gatherer
WC1 (segments not found: 23.9%)
SouthAsia
WestEurasia
EastAsia
8.5
11.8
20.3
23.7
1.7
3.4
6.8
Iranian Neolithic
BOT2016 (segments not found: 19.9%)
SouthAsia
WestEurasia
EastAsia
8.7
19.0
15.1
9.5
5.6
8.7
13.5
Steppe
Yamnaya (segments not found: 23.0%)
SouthAsia
WestEurasia
EastAsia
3.9
20.9
30.7
13.7
1.3
4.6
2.0
Stuttgart (segments not found: 27.3%)
SouthAsia
WestEurasia
EastAsia
1.8
16.6
26.3
16.6
2.6
6.1
2.6
NE1 (segments not found: 24.0%)
SouthAsia
WestEurasia
EastAsia
0.8
19.8
32.2
14.9
0.8
3.3
4.1
European Neolithic
JP14 (segments not found: 22.0%)
SouthAsia
WestEurasia
EastAsia
2.5
14.0
30.4
24.6
0
5.7
0.8
PB675 (segments not found: 25.6%)
SouthAsia
WestEurasia
EastAsia
1.8
21.9
25.4
15.8
4.4
3.5
1.8
2H10 (segments not found: 20.9%)
SouthAsia
WestEurasia
EastAsia
1.4
22.6
22.6
22.6
7.1
2.8
2H11 (segments not found: 24.2%)
SouthAsia
WestEurasia
EastAsia
2.5
16.8
30.3
14.3
0
7.6
4.2
atp016 (segments not found: 25.9%)
SouthAsia
WestEurasia
EastAsia
4.3
15.0
26.8
20.4
2.1
2.1
3.2
BA64 (segments not found: 23.1%)
SouthAsia
WestEurasia
EastAsia
1.9
28.1
20.6
15.0
7.5
3.8
sf12 (segments not found: 27.2%)
SouthAsia
WestEurasia
EastAsia
3.2
14.2
32.2
9.0
1.3
7.7
5.2
Loschbour (segments not found: 22.3%)
SouthAsia
WestEurasia
EastAsia
4.0
18.6
25.1
13.8
0.8
12.1
3.2
European Mesolithic
SRA62 (segments not found: 19.7%)
SouthAsia
WestEurasia
EastAsia
4.1
16.3
33.6
12.2
3.1
8.1
3.1

### Figure S15

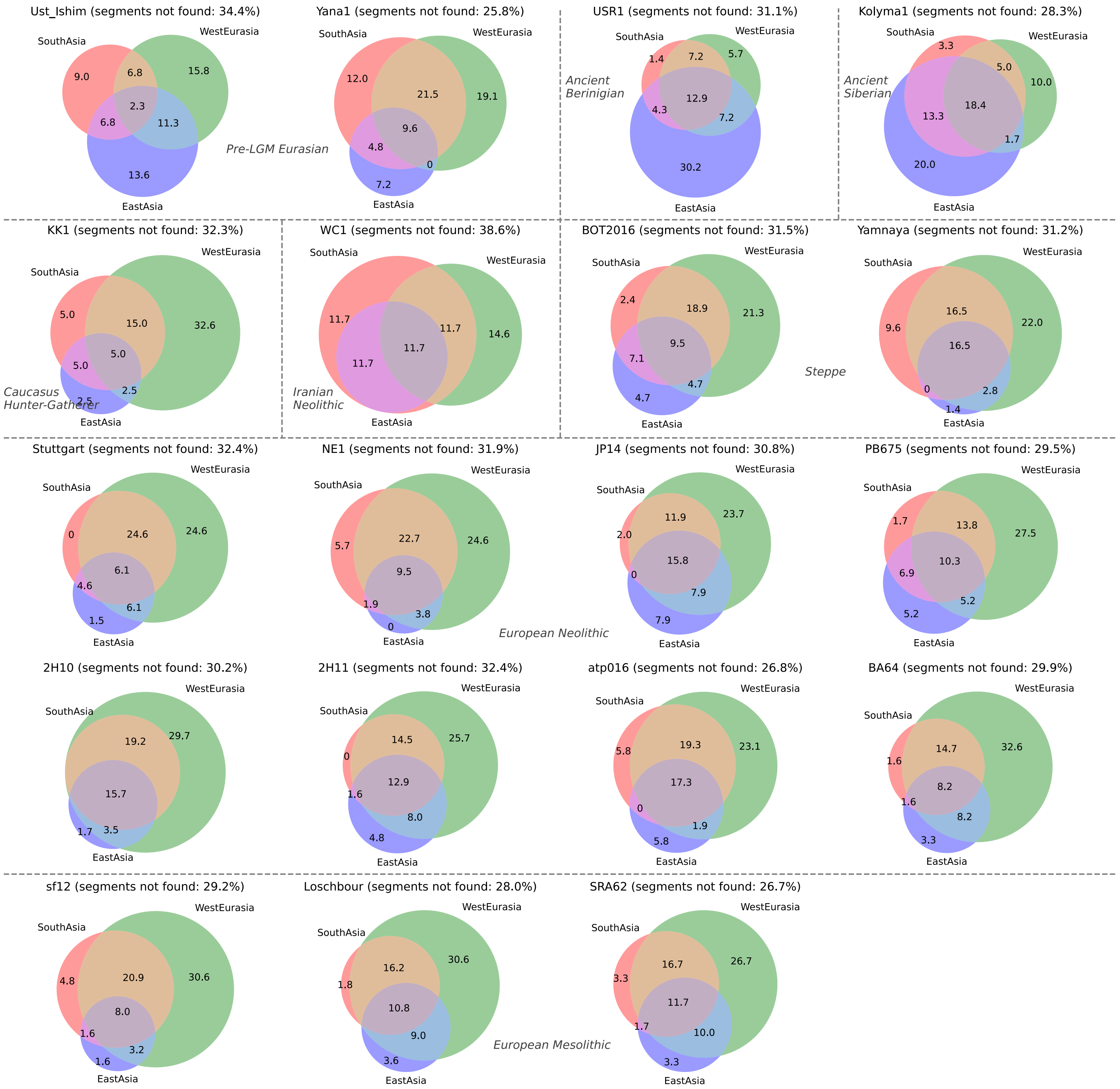

Ust_Ishim (segments not found: 34.4%)
SouthAsia
WestEurasia
EastAsia
9.0
6.8
15.8
2.3
6.8
11.3
13.6
Yana1 (segments not found: 25.8%)
SouthAsia
WestEurasia
EastAsia
12.0
21.5
19.1
9.6
4.8
0
7.2
Pre-LGM Eurasian
USR1 (segments not found: 31.1%)
SouthAsia
WestEurasia
EastAsia
1.4
7.2
5.7
12.9
4.3
7.2
30.2
Ancient Berinigian
Kolyma1 (segments not found: 28.3%)
SouthAsia
WestEurasia
EastAsia
3.3
5.0
10.0
18.4
13.3
1.7
20.0
Ancient Siberian
KK1 (segments not found: 32.3%)
SouthAsia
WestEurasia
EastAsia
5.0
15.0
32.6
5.0
5.0
2.5
2.5
Caucasus Hunter-Gatherer
WC1 (segments not found: 38.6%)
SouthAsia
WestEurasia
EastAsia
11.7
11.7
14.6
11.7
11.7
Iranian Neolithic
BOT2016 (segments not found: 31.5%)
SouthAsia
WestEurasia
EastAsia
2.4
18.9
21.3
9.5
7.1
4.7
4.7
Steppe
Yamnaya (segments not found: 31.2%)
SouthAsia
WestEurasia
EastAsia
9.6
16.5
22.0
16.5
0
2.8
1.4
Stuttgart (segments not found: 32.4%)
SouthAsia
WestEurasia
EastAsia
0
24.6
24.6
6.1
4.6
6.1
1.5
NE1 (segments not found: 31.9%)
SouthAsia
WestEurasia
EastAsia
5.7
22.7
24.6
9.5
1.9
3.8
0
European Neolithic
JP14 (segments not found: 30.8%)
SouthAsia
WestEurasia
EastAsia
2.0
11.9
23.7
15.8
0
7.9
7.9
PB675 (segments not found: 29.5%)
SouthAsia
WestEurasia
EastAsia
1.7
13.8
27.5
10.3
6.9
5.2
5.2
2H10 (segments not found: 30.2%)
SouthAsia
WestEurasia
EastAsia
19.2
29.7
15.7
1.7
3.5
2H11 (segments not found: 32.4%)
SouthAsia
WestEurasia
EastAsia
0
14.5
25.7
12.9
1.6
8.0
4.8
atp016 (segments not found: 26.8%)
SouthAsia
WestEurasia
EastAsia
5.8
19.3
23.1
17.3
0
1.9
5.8
BA64 (segments not found: 29.9%)
SouthAsia
WestEurasia
EastAsia
1.6
14.7
32.6
8.2
1.6
8.2
3.3
sf12 (segments not found: 29.2%)
SouthAsia
WestEurasia
EastAsia
4.8
20.9
30.6
8.0
1.6
3.2
1.6
Loschbour (segments not found: 28.0%)
SouthAsia
WestEurasia
EastAsia
1.8
16.2
30.6
10.8
9.0
3.6
European Mesolithic
SRA62 (segments not found: 26.7%)
SouthAsia
WestEurasia
EastAsia
3.3
16.7
26.7
11.7
1.7
10.0
3.3

Figure S16

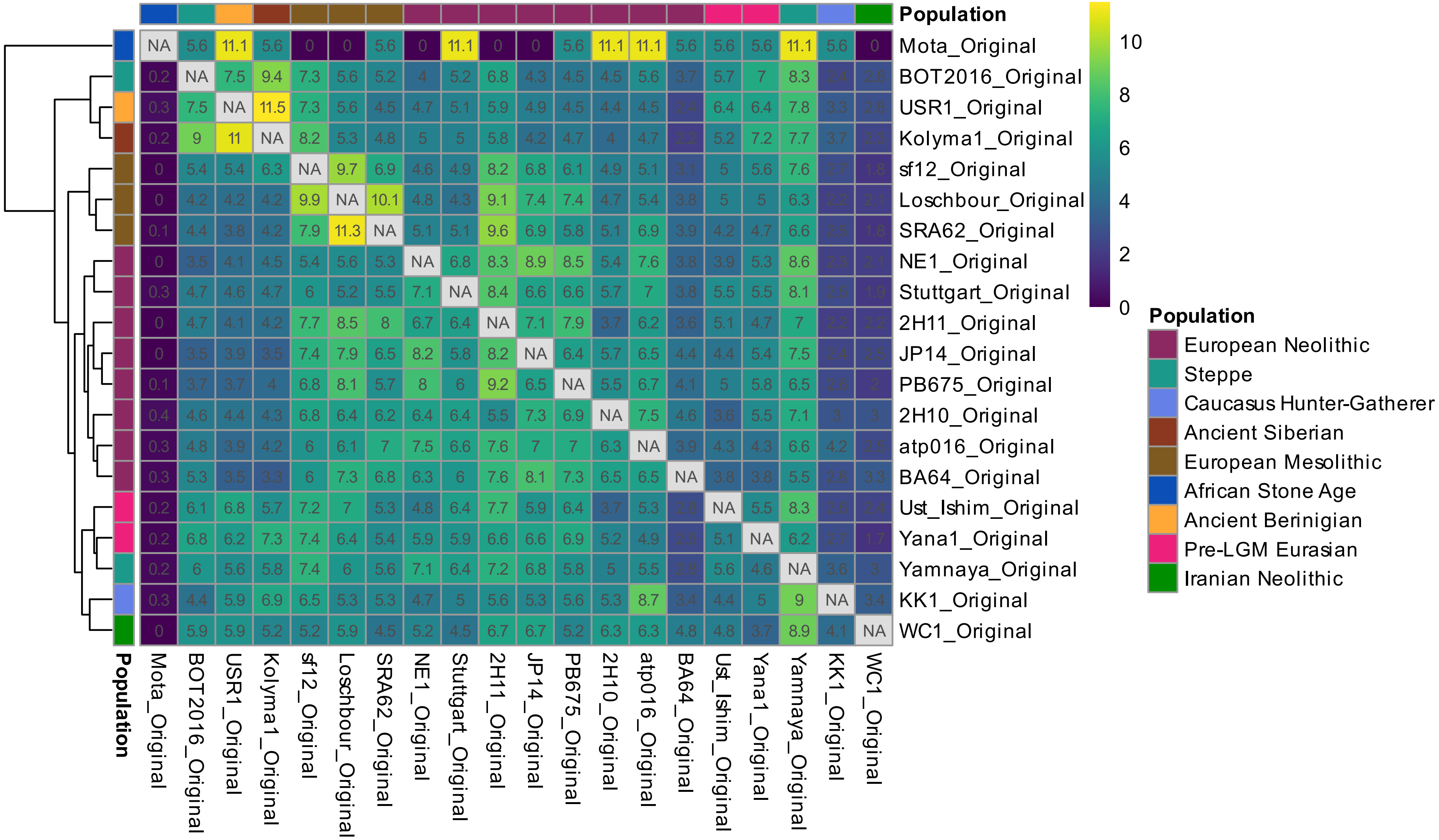

Population
Mota_Original
BOT2016_Original
USR1_Original
Kolyma1_Original
sf12_Original
Loschbour_Original
SRA62_Original
NE1_Original
Stuttgart_Original
2H11_Original
JP14_Original
PB675_Original
2H10_Original
atp016_Original
BA64_Original
Ust_Ishim_Original
Yana1_Original
Yamnaya_Original
KK1_Original
WC1_Original
10
8
6
4
2
0
Population
European Neolithic
Steppe
Caucasus Hunter-Gatherer
Ancient Siberian
European Mesolithic
African Stone Age
Ancient Berinigian
Pre-LGM Eurasian
Iranian Neolithic

Figure S17

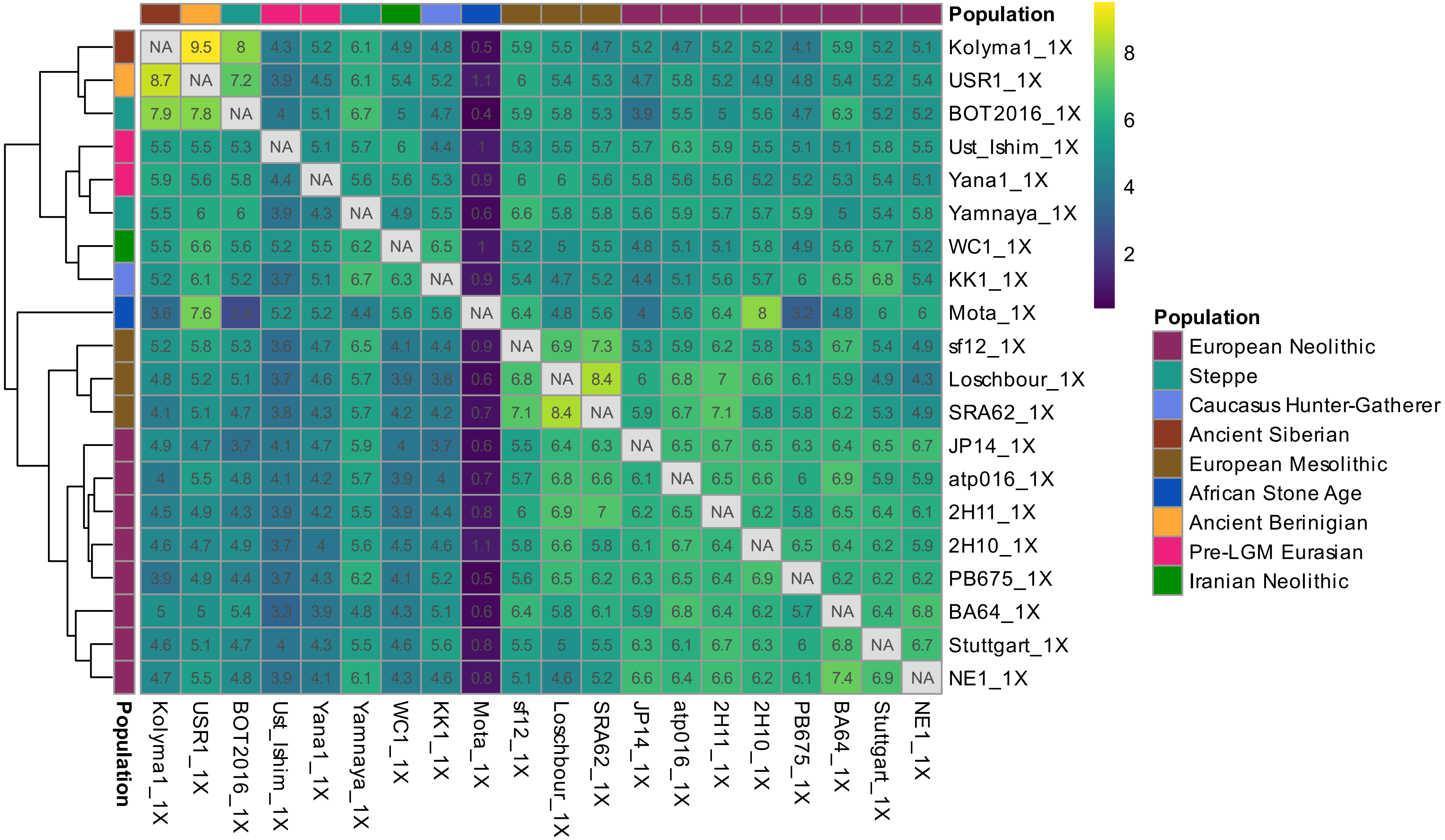
Population
Kolyma1_1X
USR1_1X
BOT2016_1X
Ust_Ishim_1X
Yana1_1X
Yamnaya_1X
WC1_1X
KK1_1X
Mota_1X
sf12_1X
Loschbour_1X
SRA62_1X
JP14_1X
atp016_1X
2H11_1X
2H10_1X
PB675_1X
BA64_1X
Stuttgart_1X
NE1_1X
NA 9.5 8 4.3 5.2 6.1 4.9 4.8 0.5 5.9 5.5 4.7 5.2 4.7 5.2 5.2 4.1 5.9 5.2 5.1
8.7 NA 7.2 3.9 4.5 6.1 5.4 5.2 1.1 6 5.4 5.3 4.7 5.8 5.2 4.9 4.8 5.4 5.2 5.4
7.9 7.8 NA 4 5.1 6.7 5 4.7 0.4 5.9 5.8 5.3 3.9 5.5 5 5.6 4.7 6.3 5.2 5.2
5.5 5.5 5.3 NA 5.1 5.7 6 4.4 1 5.3 5.5 5.7 5.7 6.3 5.9 5.5 5.1 5.1 5.8 5.5
5.9 5.6 5.8 4.4 NA 5.6 5.6 5.3 0.9 6 6 5.6 5.8 5.6 5.6 5.2 5.2 5.3 5.4 5.1
5.5 6 6 3.9 4.3 NA 4.9 5.5 0.6 6.6 5.8 5.8 5.6 5.9 5.7 5.7 5.9 5 5.4 5.8
5.5 6.6 5.6 5.2 5.5 6.2 NA 6.5 1 5.2 5 5.5 4.8 5.1 5.1 5.8 4.9 5.6 5.7 5.2
5.2 6.1 5.2 3.7 5.1 6.7 6.3 NA 0.9 5.4 4.7 5.2 4.4 5.1 5.6 5.7 6 6.5 6.8 5.4
3.6 7.6 2.4 5.2 5.2 4.4 5.6 5.6 NA 6.4 4.8 5.6 4 5.6 6.4 8 3.2 4.8 6 6
5.2 5.8 5.3 3.6 4.7 6.5 4.1 4.4 0.9 NA 6.9 7.3 5.3 5.9 6.2 5.8 5.3 6.7 5.4 4.9
4.8 5.2 5.1 3.7 4.6 5.7 3.9 3.8 0.6 6.8 NA 8.4 6 6.8 7 6.6 6.1 5.9 4.9 4.3
4.1 5.1 4.7 3.8 4.3 5.7 4.2 4.2 0.7 7.1 8.4 NA 5.9 6.7 7.1 5.8 5.8 6.2 5.3 4.9
4.9 4.7 3.7 4.1 4.7 5.9 4 3.7 0.6 5.5 6.4 6.3 NA 6.5 6.7 6.5 6.3 6.4 6.5 6.7
4 5.5 4.8 4.1 4.2 5.7 3.9 4 0.7 5.7 6.8 6.6 6.1 NA 6.5 6.6 6 6.9 5.9 5.9
4.5 4.9 4.3 3.9 4.2 5.5 3.9 4.4 0.8 6 6.9 7 6.2 6.5 NA 6.2 5.8 6.5 6.4 6.1
4.6 4.7 4.9 3.7 4 5.6 4.5 4.6 1.1 5.8 6.6 5.8 6.1 6.7 6.4 NA 6.5 6.4 6.2 5.9
3.9 4.9 4.4 3.7 4.3 6.2 4.1 5.2 0.5 5.6 6.5 6.2 6.3 6.5 6.4 6.9 NA 6.2 6.2 6.2
5 5 5.4 3.3 3.9 4.8 4.3 5.1 0.6 6.4 5.8 6.1 5.9 6.8 6.4 6.2 5.7 NA 6.4 6.8
4.6 5.1 4.7 4 4.3 5.5 4.6 5.6 0.8 5.5 5 5.5 6.3 6.1 6.7 6.3 6 6.8 NA 6.7
4.7 5.5 4.8 3.9 4.1 6.1 4.3 4.6 0.8 5.1 4.6 5.2 6.6 6.4 6.6 6.2 6.1 7.4 6.9 NA
8
6
4
2
Population
European Neolithic
Steppe
Caucasus Hunter-Gatherer
Ancient Siberian
European Mesolithic
African Stone Age
Ancient Berinigian
Pre-LGM Eurasian
Iranian Neolithic

Figure S18

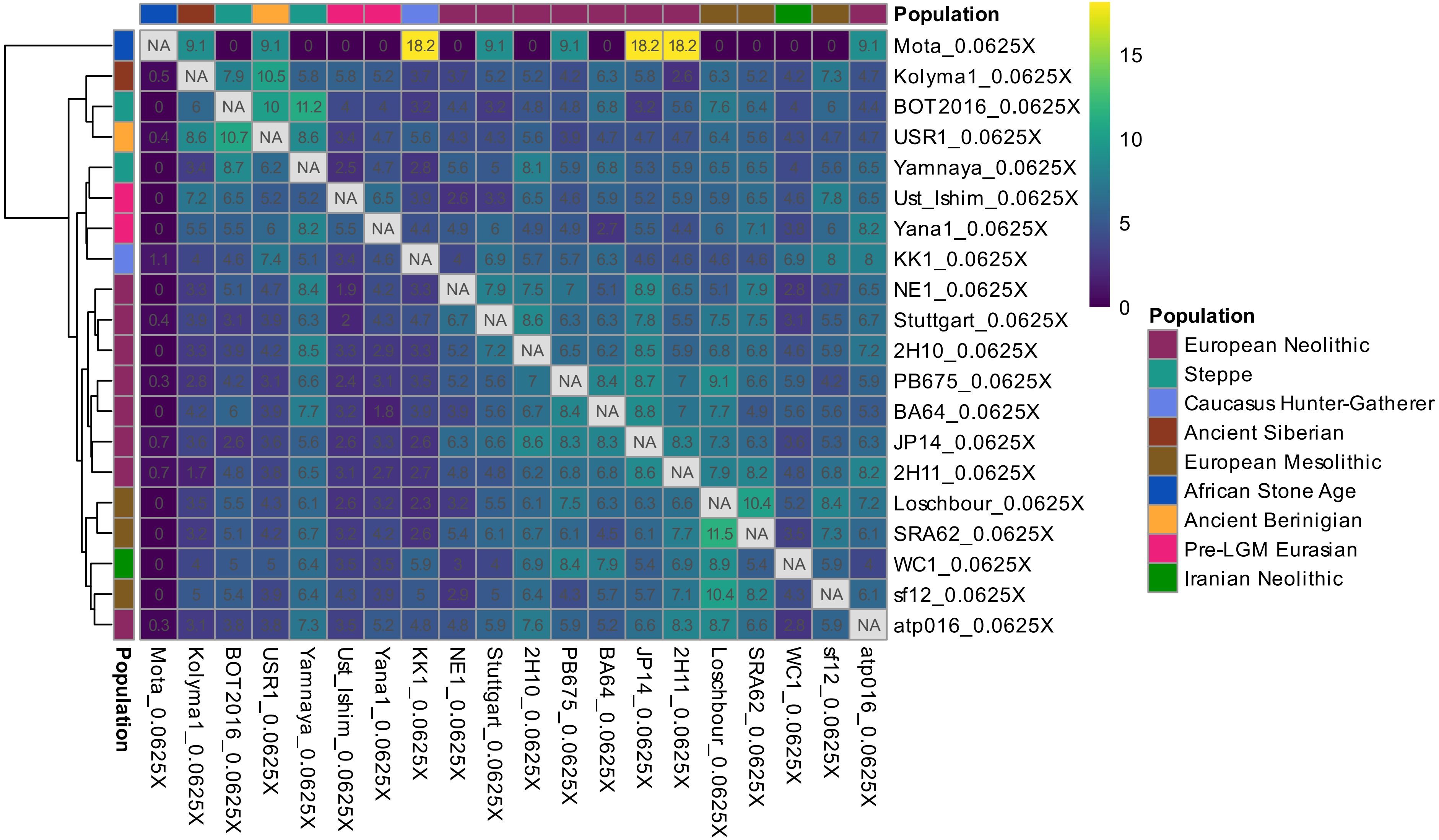

Population
Mota_0.0625X
Kolyma1_0.0625X
BOT2016_0.0625X
USR1_0.0625X
Yamnaya_0.0625X
Ust_Ishim_0.0625X
Yana1_0.0625X
KK1_0.0625X
NE1_0.0625X
Stuttgart_0.0625X
2H10_0.0625X
PB675_0.0625X
BA64_0.0625X
JP14_0.0625X
2H11_0.0625X
Loschbour_0.0625X
SRA62_0.0625X
WC1_0.0625X
sf12_0.0625X
atp016_0.0625X
15
10
5
0
Population
European Neolithic
Steppe
Caucasus Hunter-Gatherer
Ancient Siberian
European Mesolithic
African Stone Age
Ancient Berinigian
Pre-LGM Eurasian
Iranian Neolithic

Figure S19

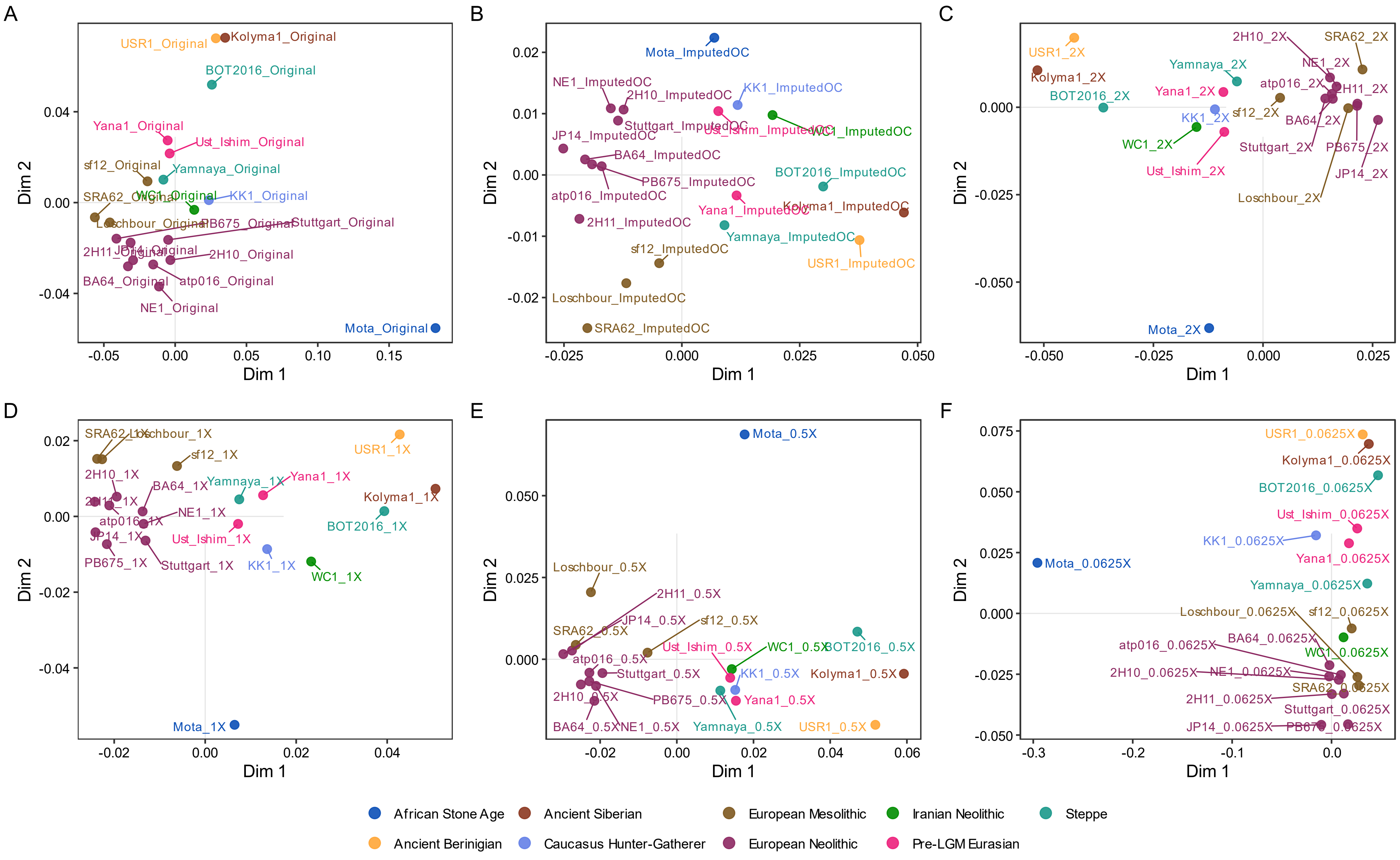
A
USR1_Original
Kolyma1_Original
BOT2016_Original
Yana1_Original
Ust_Ishim_Original
sf12_Original
Yamnaya_Original
SRA62_Original
WC1_Original
KK1_Original
Loschbour_Original
PB675_Original
Stuttgart_Original
2H11_Original
JP14_Original
2H10_Original
BA64_Original
atp016_Original
NE1_Original
Mota_Original
B
Mota_ImputedOC
NE1_ImputedOC
2H10_ImputedOC
KK1_ImputedOC
Stuttgart_ImputedOC
Ust_Ishim_ImputedOC
WC1_ImputedOC
JP14_ImputedOC
BA64_ImputedOC
PB675_ImputedOC
BOT2016_ImputedOC
atp016_ImputedOC
Yana1_ImputedOC
Kolyma1_ImputedOC
2H11_ImputedOC
Yamnaya_ImputedOC
sf12_ImputedOC
USR1_ImputedOC
Loschbour_ImputedOC
SRA62_ImputedOC
C
USR1_2X
2H10_2X
SRA62_2X
Kolyma1_2X
Yamnaya_2X
NE1_2X
BOT2016_2X
Yana1_2X
atp016_2X
2H11_2X
KK1_2X
sf12_2X
BA64_2X
WC1_2X
Stuttgart_2X
PB675_2X
Ust_Ishim_2X
JP14_2X
Loschbour_2X
Mota_2X
D
SRA62_1X
Loschbour_1X
sf12_1X
USR1_1X
2H10_1X
BA64_1X
Yamnaya_1X
Yana1_1X
Kolyma1_1X
2H11_1X
NE1_1X
atp016_1X
BOT2016_1X
JP14_1X
Ust_Ishim_1X
PB675_1X
Stuttgart_1X
KK1_1X
WC1_1X
Mota_1X
E
Mota_0.5X
Loschbour_0.5X
2H11_0.5X
JP14_0.5X
sf12_0.5X
SRA62_0.5X
Ust_Ishim_0.5X
WC1_0.5X
BOT2016_0.5X
atp016_0.5X
Stuttgart_0.5X
KK1_0.5X
Kolyma1_0.5X
2H10_0.5X
PB675_0.5X
Yana1_0.5X
BA64_0.5X
NE1_0.5X
Yamnaya_0.5X
USR1_0.5X
F
USR1_0.0625X
Kolyma1_0.0625X
BOT2016_0.0625X
Ust_Ishim_0.0625X
KK1_0.0625X
Yana1_0.0625X
Mota_0.0625X
Yamnaya_0.0625X
Loschbour_0.0625X
sf12_0.0625X
BA64_0.0625X
atp016_0.0625X
WC1_0.0625X
2H10_0.0625X
NE1_0.0625X
SRA62_0.0625X
2H11_0.0625X
Stuttgart_0.0625X
JP14_0.0625X
PB675_0.0625X
Dim 1
Dim 2
African Stone Age
Ancient Siberian
European Mesolithic
Iranian Neolithic
Steppe
Ancient Berinigian
Caucasus Hunter-Gatherer
European Neolithic
Pre-LGM Eurasian

Figure S20

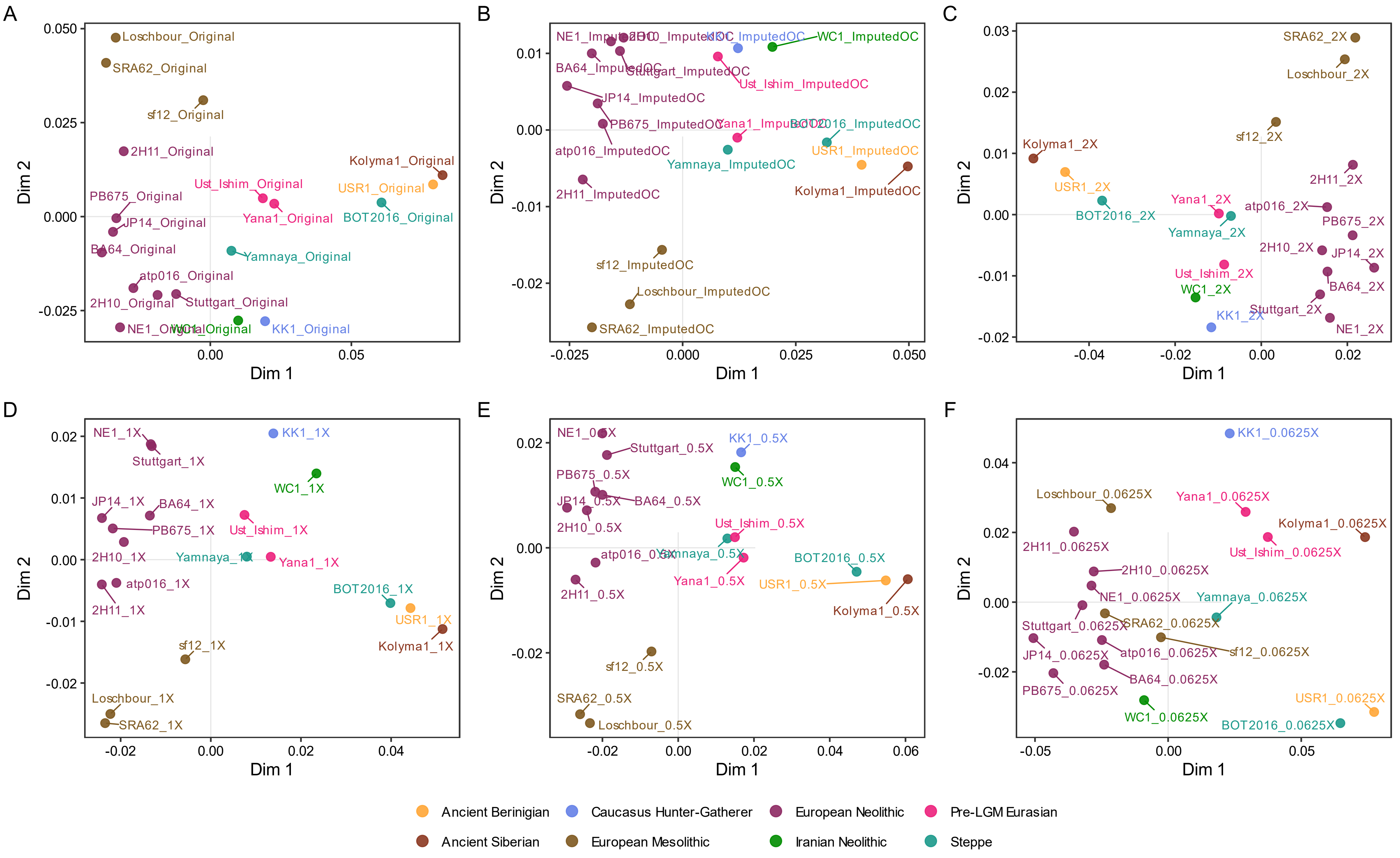
A
Loschbour_Original
SRA62_Original
sf12_Original
2H11_Original
Kolyma1_Original
Ust_Ishim_Original
USR1_Original
PB675_Original
JP14_Original
Yana1_Original
BOT2016_Original
BA64_Original
Yamnaya_Original
atp016_Original
2H10_Original
Stuttgart_Original
NE1_Original
WC1_Original
KK1_Original
B
NE1_ImputedOC
2H10_ImputedOC
KK1_ImputedOC
WC1_ImputedOC
BA64_ImputedOC
Stuttgart_ImputedOC
Ust_Ishim_ImputedOC
JP14_ImputedOC
PB675_ImputedOC
Yana1_ImputedOC
BOT2016_ImputedOC
atp016_ImputedOC
USR1_ImputedOC
Yamnaya_ImputedOC
2H11_ImputedOC
Kolyma1_ImputedOC
sf12_ImputedOC
Loschbour_ImputedOC
SRA62_ImputedOC
C
SRA62_2X
Loschbour_2X
sf12_2X
Kolyma1_2X
USR1_2X
2H11_2X
Yana1_2X
atp016_2X
BOT2016_2X
PB675_2X
Yamnaya_2X
2H10_2X
JP14_2X
Ust_Ishim_2X
BA64_2X
WC1_2X
KK1_2X
Stuttgart_2X
NE1_2X
D
NE1_1X
KK1_1X
Stuttgart_1X
WC1_1X
JP14_1X
BA64_1X
PB675_1X
Ust_Ishim_1X
2H10_1X
Yamnaya_1X
Yana1_1X
atp016_1X
2H11_1X
BOT2016_1X
USR1_1X
sf12_1X
Kolyma1_1X
Loschbour_1X
SRA62_1X
E
NE1_0.5X
Stuttgart_0.5X
KK1_0.5X
PB675_0.5X
WC1_0.5X
JP14_0.5X
BA64_0.5X
2H10_0.5X
Ust_Ishim_0.5X
atp016_0.5X
Yamnaya_0.5X
BOT2016_0.5X
Yana1_0.5X
USR1_0.5X
2H11_0.5X
Kolyma1_0.5X
sf12_0.5X
SRA62_0.5X
Loschbour_0.5X
F
KK1_0.0625X
Loschbour_0.0625X
Yana1_0.0625X
Kolyma1_0.0625X
2H11_0.0625X
Ust_Ishim_0.0625X
2H10_0.0625X
NE1_0.0625X
Yamnaya_0.0625X
Stuttgart_0.0625X
SRA62_0.0625X
JP14_0.0625X
atp016_0.0625X
sf12_0.0625X
BA64_0.0625X
PB675_0.0625X
USR1_0.0625X
WC1_0.0625X
BOT2016_0.0625X
Dim 1
Dim 2
Ancient Beringian
Caucasus Hunter-Gatherer
European Neolithic
Pre-LGM Eurasian
Ancient Siberian
European Mesolithic
Iranian Neolithic
Steppe

Figure S21

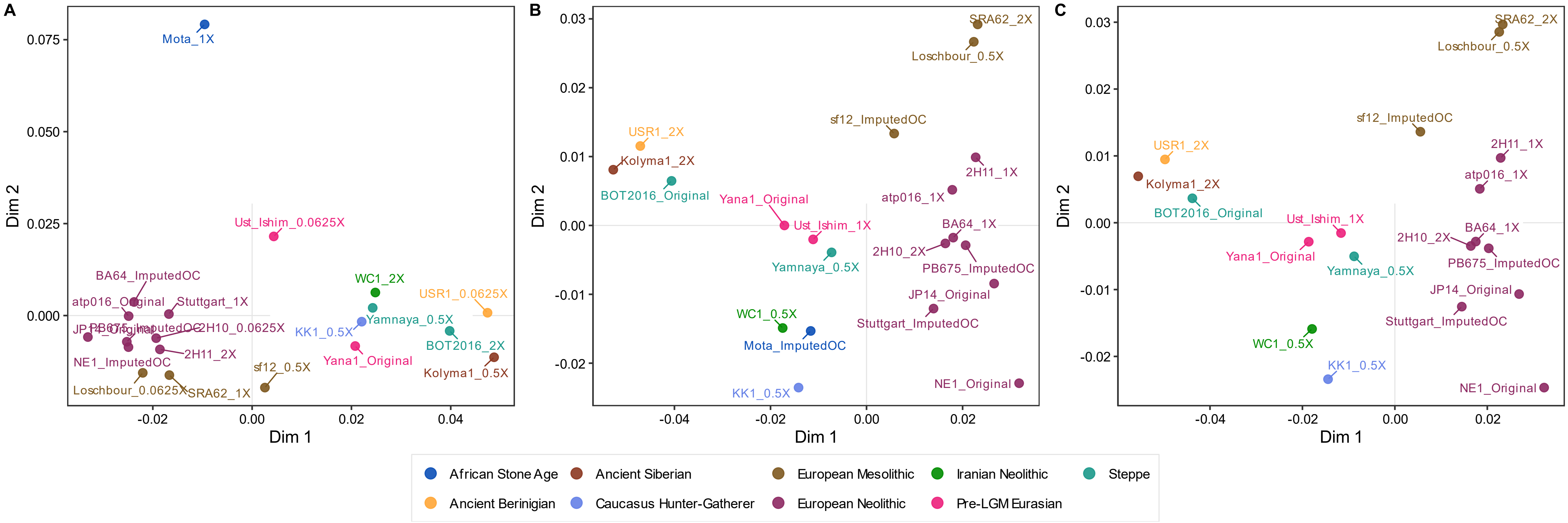
A
B
C
Dim 1
Dim 2
Mota_1X
Ust_Ishim_0.0625X
BA64_ImputedOC
atp016_Original
Stuttgart_1X
JP14_Original
PB675_ImputedOC
2H10_0.0625X
2H11_2X
NE1_ImputedOC
Loschbour_0.0625X
SRA62_1X
sf12_0.5X
WC1_2X
USR1_0.0625X
KK1_0.5X
Yamnaya_0.5X
BOT2016_2X
Yana1_Original
Kolyma1_0.5X
SRA62_2X
Loschbour_0.5X
sf12_ImputedOC
USR1_2X
Kolyma1_2X
BOT2016_Original
Yana1_Original
Ust_Ishim_1X
2H11_1X
atp016_1X
BA64_1X
2H10_2X
PB675_ImputedOC
Yamnaya_0.5X
JP14_Original
Stuttgart_ImputedOC
WC1_0.5X
Mota_ImputedOC
NE1_Original
KK1_0.5X
African Stone Age
Ancient Siberian
European Mesolithic
Iranian Neolithic
Steppe
Ancient Berinigian
Caucasus Hunter-Gatherer
European Neolithic
Pre-LGM Eurasian

Figure S22

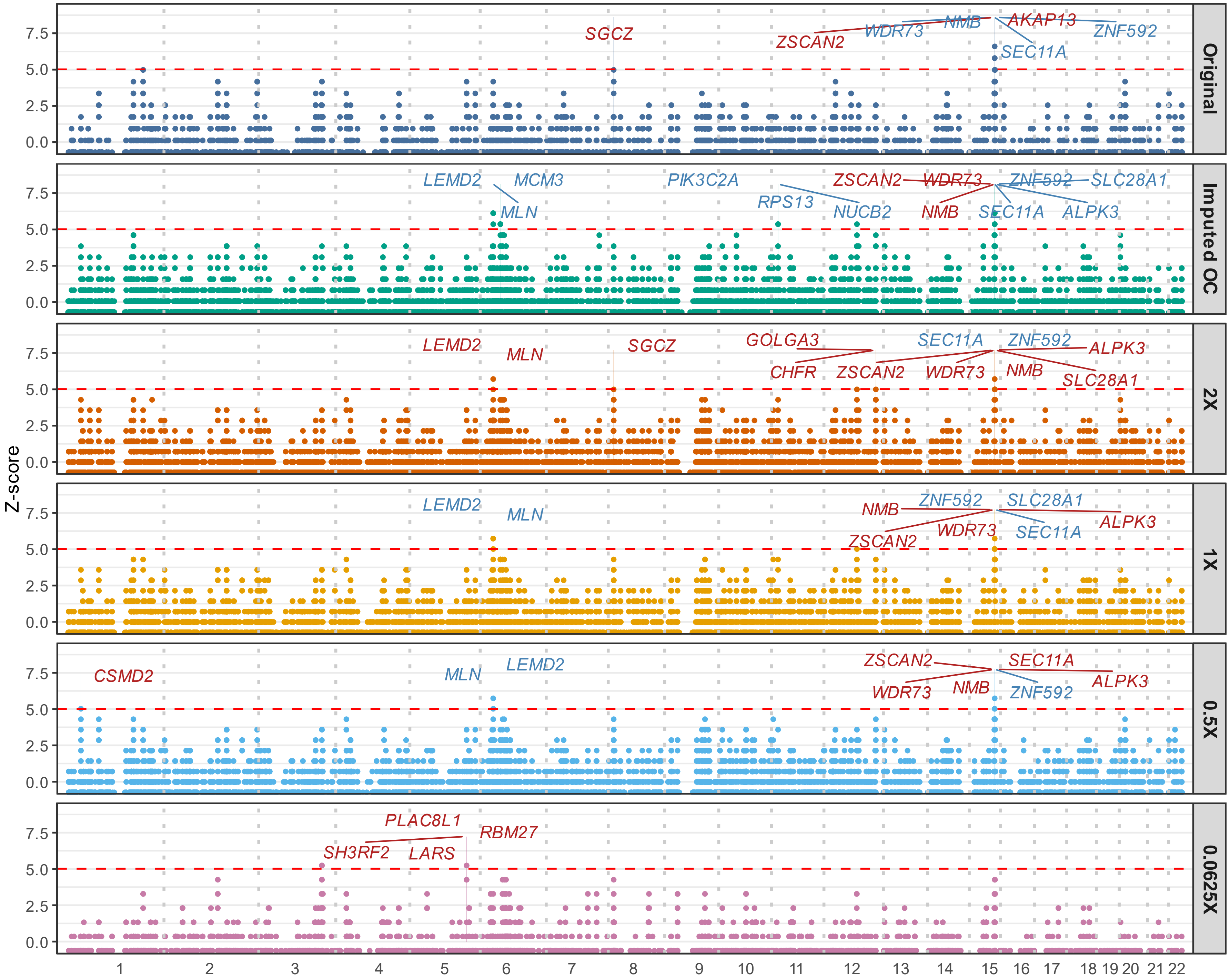

Original
SGCZ
ZSCAN2
WDR73
NMB
AKAP13
ZNF592
SEC11A
Imputed OC
LEMD2
MCM3
MLN
PIK3C2A
RPS13
NUCB2
ZSCAN2
WDR73
NMB
ZNF592
SLC28A1
SEC11A
ALPK3
2X
LEMD2
MLN
SGCZ
GOLGA3
CHFR
ZSCAN2
SEC11A
WDR73
ZNF592
NMB
ALPK3
SLC28A1
1X
LEMD2
MLN
NMB
ZNF592
SLC28A1
WDR73
ZSCAN2
SEC11A
ALPK3
0.5X
CSMD2
MLN
LEMD2
ZSCAN2
WDR73
NMB
SEC11A
ZNF592
ALPK3
0.0625X
PLAC8L1
SH3RF2
LARS
RBM27
Z-score
0.0
2.5
5.0
7.5
1 2 3 4 5 6 7 8 9 10 11 12 13 14 15 16 17 18 19 20 21 22

### Figure S23

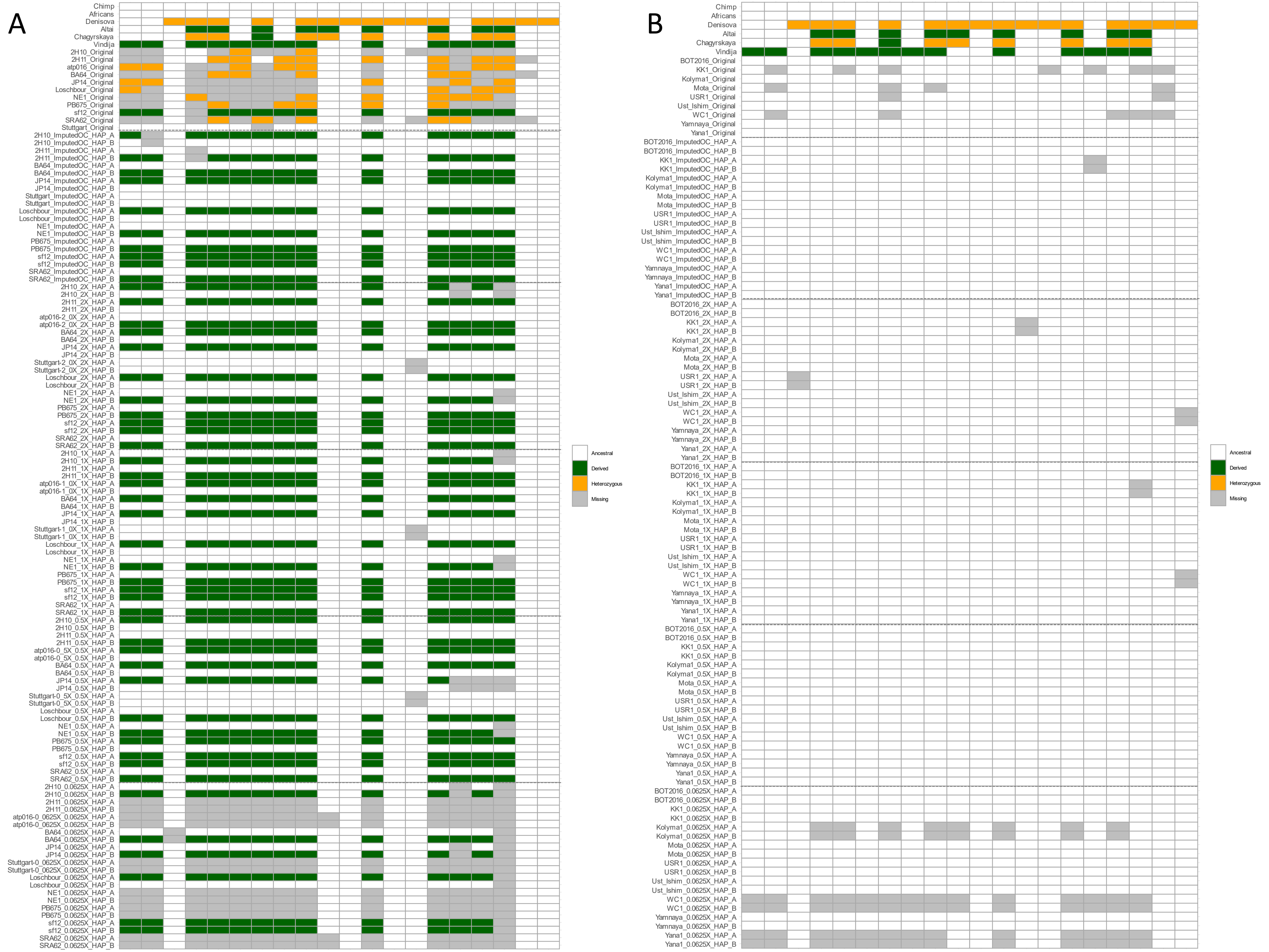
A
B
Chimp
Africans
Denisova
Altai
Chagyrskaya
Vindija
Ancestral
Derived
Heterozygous
Missing
