## Supplementary figures and images for "Archaic ancestry inference in imputed ancient human genomes"

### Figures1-4

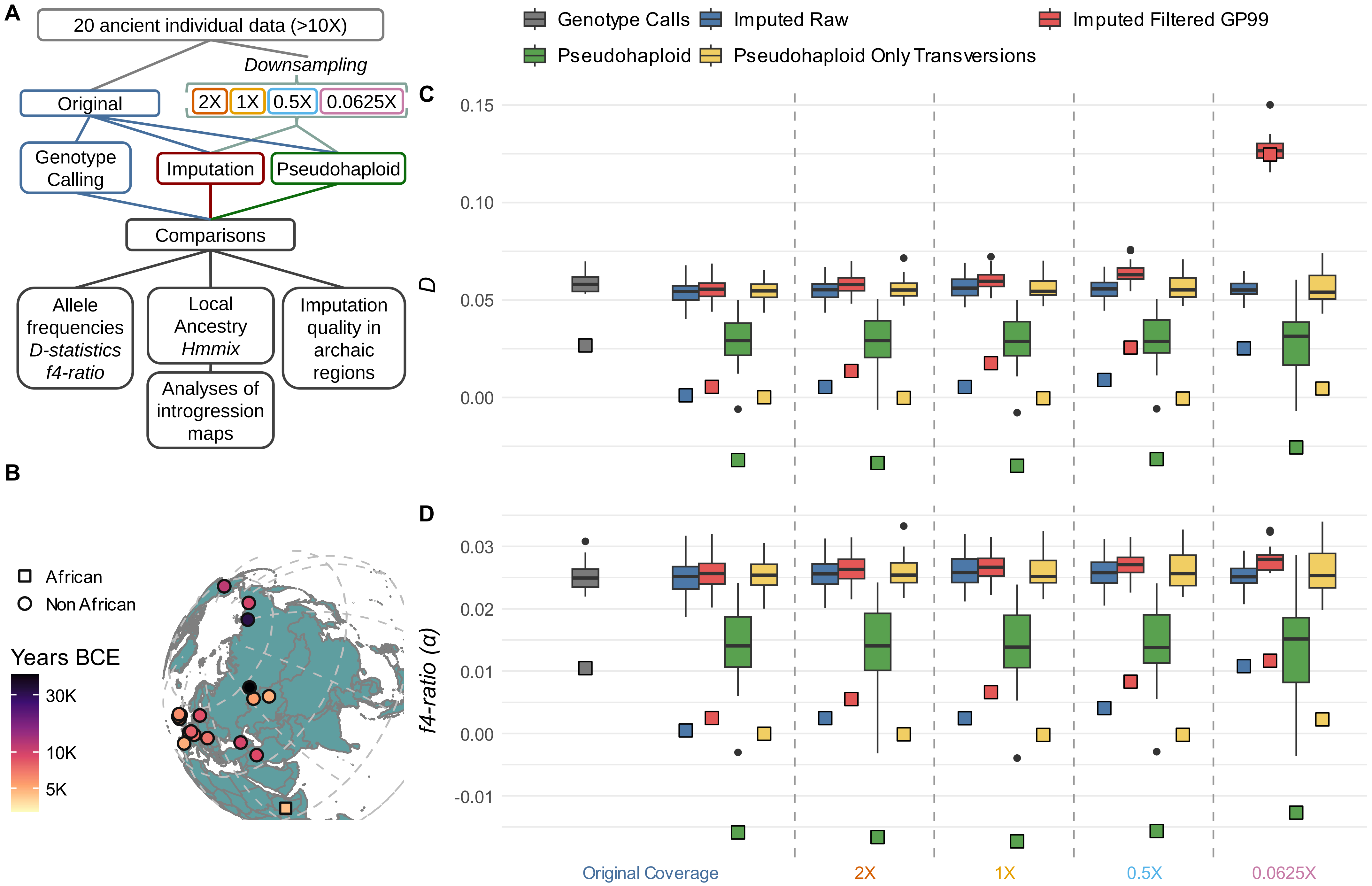

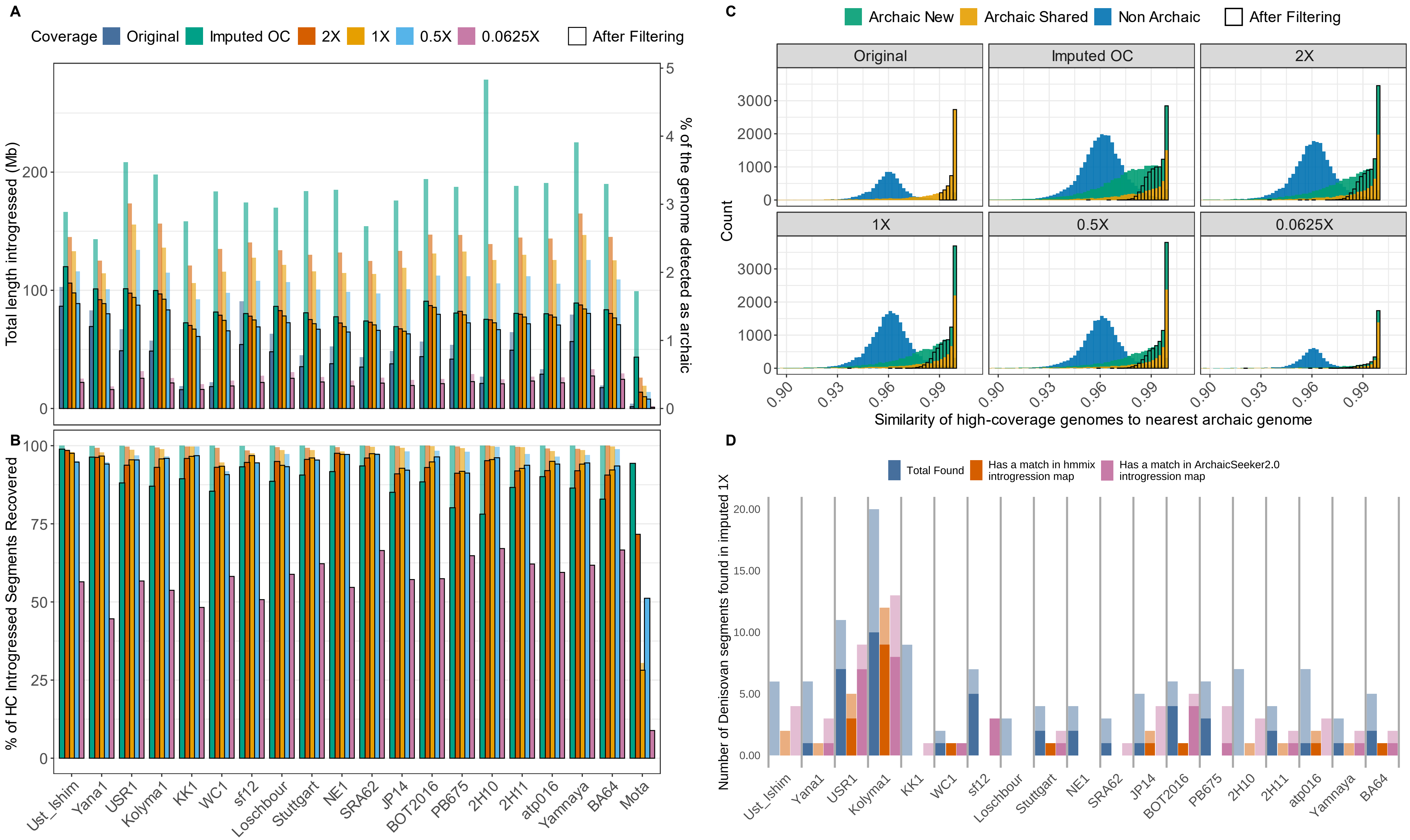

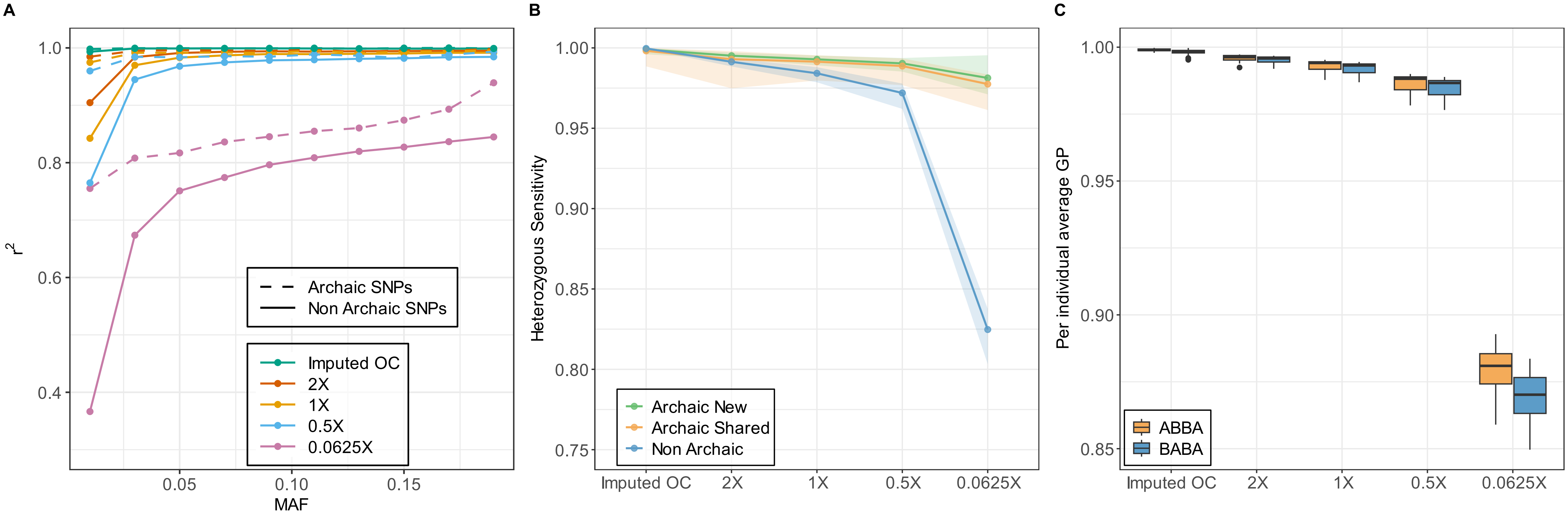

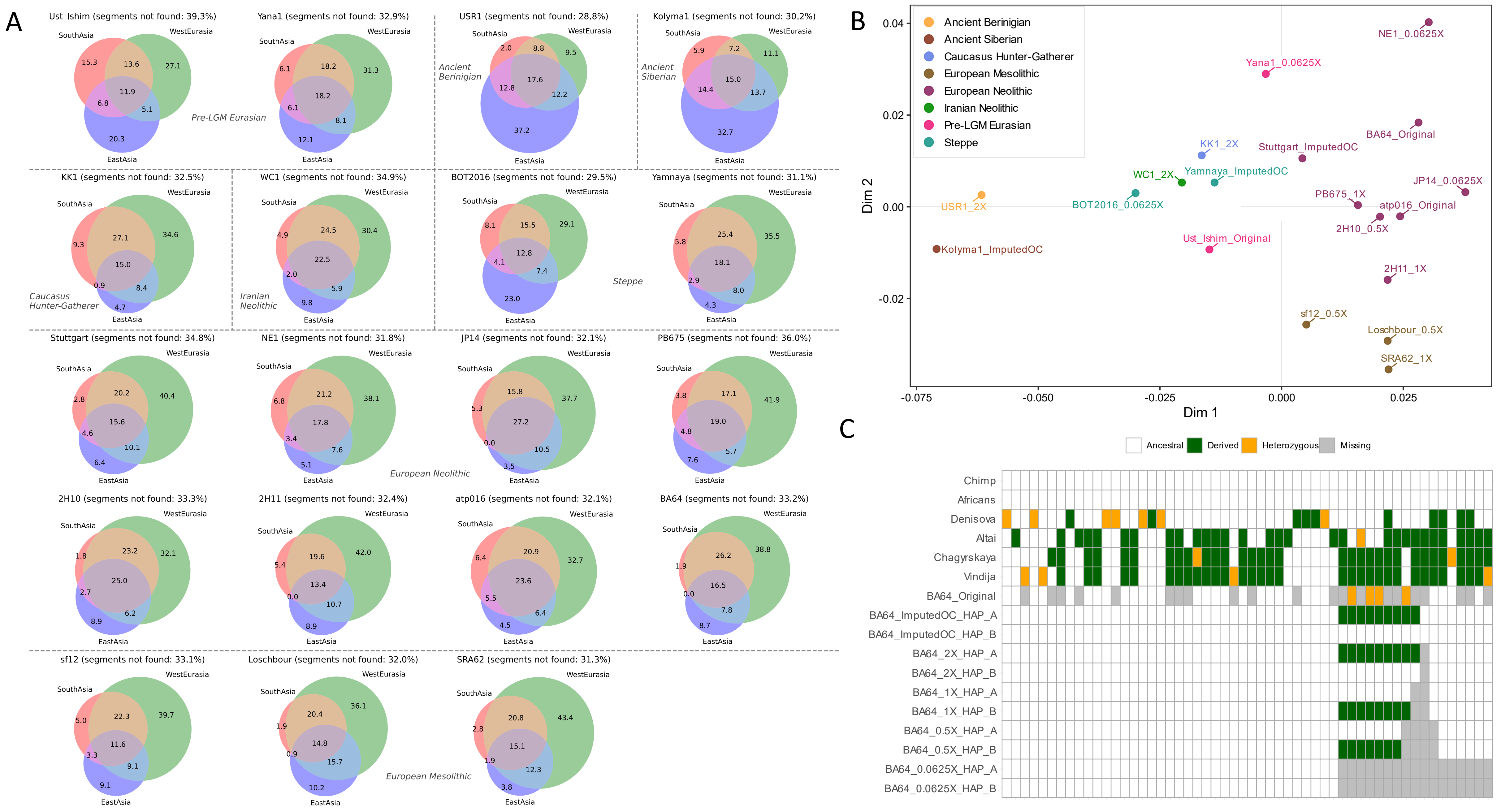
